# Modeling Alternative Conformational States in CASP16

**DOI:** 10.1101/2025.09.02.673835

**Authors:** Namita Dube, Theresa A. Ramelot, Tiburon L. Benavides, Yuanpeng J. Huang, John Moult, Andriy Kryshtafovych, Gaetano T. Montelione

## Abstract

The CASP16 Ensemble Prediction experiment assessed advances in methods for modeling proteins, nucleic acids, and their complexes in multiple conformational states. Targets included systems with experimental structures determined in two or three states, evaluated by direct comparison to experimental coordinates, as well as domain–linker–domain (D–L–D) targets assessed against statistical models from NMR and SAXS data. This paper focuses on the former class of multi-state targets. Ten ensembles were released as community challenges, including ligand-induced conformational changes, protein–DNA complexes, a trimeric protein, a stem-loop RNA, and multiple oligomeric states of a single RNA. For five targets, some groups produced reasonably accurate models of both reference states (best TM-score >0.75). However, with the exception of one protein–ligand complex (T1214), where an apo structure was available as a template, predictors generally failed to capture key structural details distinguishing the states. Overall, accuracy was significantly lower than for single-state targets in other CASP experiments. The most successful approaches generated multiple AlphaFold2 models using enhanced multiple sequence alignments and sampling protocols, followed by model quality based selection. While the AlphaFold3 server performed well on several targets, individual groups outperformed it in specific cases. By contrast, predictions for one protein–DNA complex, three RNA targets, and multiple oligomeric RNA states consistently fell short (TM-score <0.75). These results highlight both progress and persistent challenges in multi-state prediction. Despite recent advances, accurate modeling of conformational ensembles, particularly RNA and large multimeric assemblies, remains a critical frontier for structural biology.

## Introduction

Biomolecules, including proteins and nucleic acids, are not static objects, but rather exist as conformational distributions in dynamic equilibrium. These dynamics often underpin their functions. While recent advances in structure prediction have demonstrated outstanding success in modeling dominant conformational states of proteins, RNAs, DNAs, and their complexes [1,2], there have been few critical assessments of modeling alternative or multiple conformational states.

The Critical Assessment of Protein Structure Prediction (CASP) provides a platform for “blind” assessment of protein structure models, in which the experimental reference structure is not available to the predictor groups. To address the important challenge of modeling multiple conformational states, the 2022 CASP15 community experiment included, for the first time, computing multiple conformations for protein and RNA structures [3]. The results of this first CASP ensembles experiment were mixed, with full or partial success in reproducing both conformational states for four of nine targets. Across these successfully-modeled “ensemble” targets, enhanced sampling using variations of AlphaFold2 [4] was the most effective approach.

The CASP16 Ensemble Prediction experiment included two types of targets: those assessed in the traditional way by comparing predicted models directly to experimentally determined coordinates, and domain–linker–domain (D–L–D) targets in which submitted model sets were evaluated against statistical models derived from experimental NMR and SAXS data. The first type is the focus of this paper, while the second is discussed in another paper in this special issue of PROTEINS [5]. The CASP16 ensembles experiment was also guided by discussions in the on-line CASP Special Interest Group on Modeling Conformational Ensembles [6,7].

In the CASP experiment, we define "*ensembles*" as collections of two or more structural conformations adopted by the same macromolecular sequence, or sometimes by alternative conformations stabilized through ligand binding or small sequence variations. These ensembles range in complexity from discrete two-state systems, such as open/closed lids regulating access to enzyme active sites, to broad distributions of conformations typical of intrinsically disordered proteins (IDPs) or intrinsically disordered regions (IDRs). We define "*conformational states*" as distinct molecular conformations that have been, or could in principle be, distinguished by experimental data [8]. This definition is useful in practice because it postulates predicted states to be experimentally testable, even if they have not yet been observed. Importantly, it also encompasses alternative conformational states that may be populated under certain conditions but have not yet been distinguished experimentally. Such states still meet our definition, provided they are in principle experimentally resolvable.

A total of 12 targets were identified from experimental submissions for the CASP16 ensemble assessment, representing a diverse set of biomolecular systems including membrane proteins, protein monomers, protein-DNA complexes, RNA molecules, and protein oligomers. Ten of these targets (excluding the domain-linker-domain system mentioned above) are assessed in this study; some were single-chain substructures of other larger multi-chain targets. For each target, predictors were told the number of states in the reference ensemble. For five targets, some predictors generated models that are reasonably accurate for both states (best model TM score > 0.75). For the other five, all predictors failed to achieve accurate models (TM < 0.75) of one or more states. These outcomes demonstrate both the potential of current methods and the substantial challenges that remain in accurate multi-state conformational modeling

## Results

The CASP Ensemble Prediction experiment assessed the accuracy of collections of alternative conformational states of biomolecules for several types of ensembles (**Table 1**) including: (i) hinges, (ii) lids / cryptic ligand binding sites, (iii) conformational rearrangements, (iv) intrinsically disordered regions (IDRs) of interdomain linkers, and (v) variations in oligomeric state (or quaternary structure). In CASP16, "blind" targets, not accessible to predictors, corresponding to each of these classes were available from experimental NMR, X-ray crystallography, and/or cryogenic electron microscopy (cryoEM) studies. Other types of ensembles indicated in **Table 1**, including fold switching (FS) and intrinsically disordered proteins (IDPs), were not available in the CASP16 target set.

**Table 1.** Classes of alternative conformational states and conformational modulators.

| Alternative Conformational States | Modulators |
| --- | --- |
| Hinges (HG) | Boltzmann Distributions (BD)<br>• Temperature<br>• Pressure<br>• “Invisible states” |
| Lids / Cryptic Sites (LC) |  |
| Rearrangements (RA) | Ligand Binding (LB) |
| Fold Switching (FS) | Biomolecule Binding (BB) |
| Intrinsically disordered proteins (IDPs) | pH change (PH) |
| Intrinsically disordered regions (IDRs), including linkers between domains | Mutation (MP) |
|  | Post translational modifications (PT) |
| Oligomer State (OS) | Post translational modifications (PT) |

Each of these classes of ensembles (**Table 1**) arises from conformational equilibria that are determined by thermodynamics. The simplest systems exhibit detectable alternative states populated according to Boltzmann distributions governed by temperature and/or pressure. These include both significantly populated multiple conformational states that can be detected simultaneously in the same sample [for example as observed for dengue virus protease [9]], to very lowly populated states (sometimes referred to as “invisible states”), which can be detected by specific NMR spectroscopy experiments such as relaxation dispersion [10] or chemical exchange by saturation transfer (CEST) [11]. Alternative conformational state distributions may also be modulated by binding to partners, including small ligands or other biomolecules. They may also arise (or shift in population) due to changes in the pH (affecting the ionization states of some residues), point mutations, sequence extensions or truncations, or posttranslational modifications. In the CASP16 Ensembles Prediction experiment, the CASP community was asked to predict both kinds of multiple conformational ensembles; those in which multiple states are present at equilibrium in a single sample and other cases where the alternative conformation is induced to form by ligand or biomolecular partner binding. While the relative populations of states is also a critical feature of the ensemble, aside from the protein A domain-linker-domain experiment discussed elsewhere [5] the relative conformer populations were not requested from predictors, and no significant effort was made by predictor groups to estimate them.

The 12 targets of the CASP16 Ensembles experiment, each consisting of two or more conformational states, are listed in **Table 2**. These targets are classified by *oligomer class* as defined in previous CASP experiments [1,2], number of residues (nres), experimental method used for structure determination, number of models requested from predictors, and number of participating groups. For target T1214holo predictors were asked to submit 5 models, which were assessed by finding the model best matching to the ligand-bound state. For target R1203v1/v2 predictors were asked to submit 5 models, each assessed against both reference states. In later stages of the experiment, predictors were instructed to submit 5 models for each state. For these targets, some groups submitted models for only one state, so the number of groups submitting models for both states is slightly lower than the total number submitting for the first state. **Table 2** also summarizes the total number of models submitted by all predictor groups, the *ensemble class* as defined in **Table 1**, and a general comment about the nature of the ensemble. T-type, M-type, and R-type targets are single chain protein or homomers, heteromultimers (protein-nucleic acid), and RNA targets, respectively. Ensemble predictions for targets T1214, M1228 (and its protein substructure T1228), M1239 (and its protein substructure T1239), T1249, T1294, R1203, R1253, and R1283 are discussed in the following sections. Targets T1200 and T1300 are domain-linker-domain (D-L-D) targets, in which the domains are treated as static conformations with relative orientations and distances determined by the conformation of the interdomain linker, an IDR. An assessment of these predictions against experimental data including NMR residual dipolar coupling (RDC) obtained using internal orientation by a paramagnetic metal [12,13] and small angle X-ray scattering (SAXS) [14,15], are described elsewhere [5].

**Table 2:**
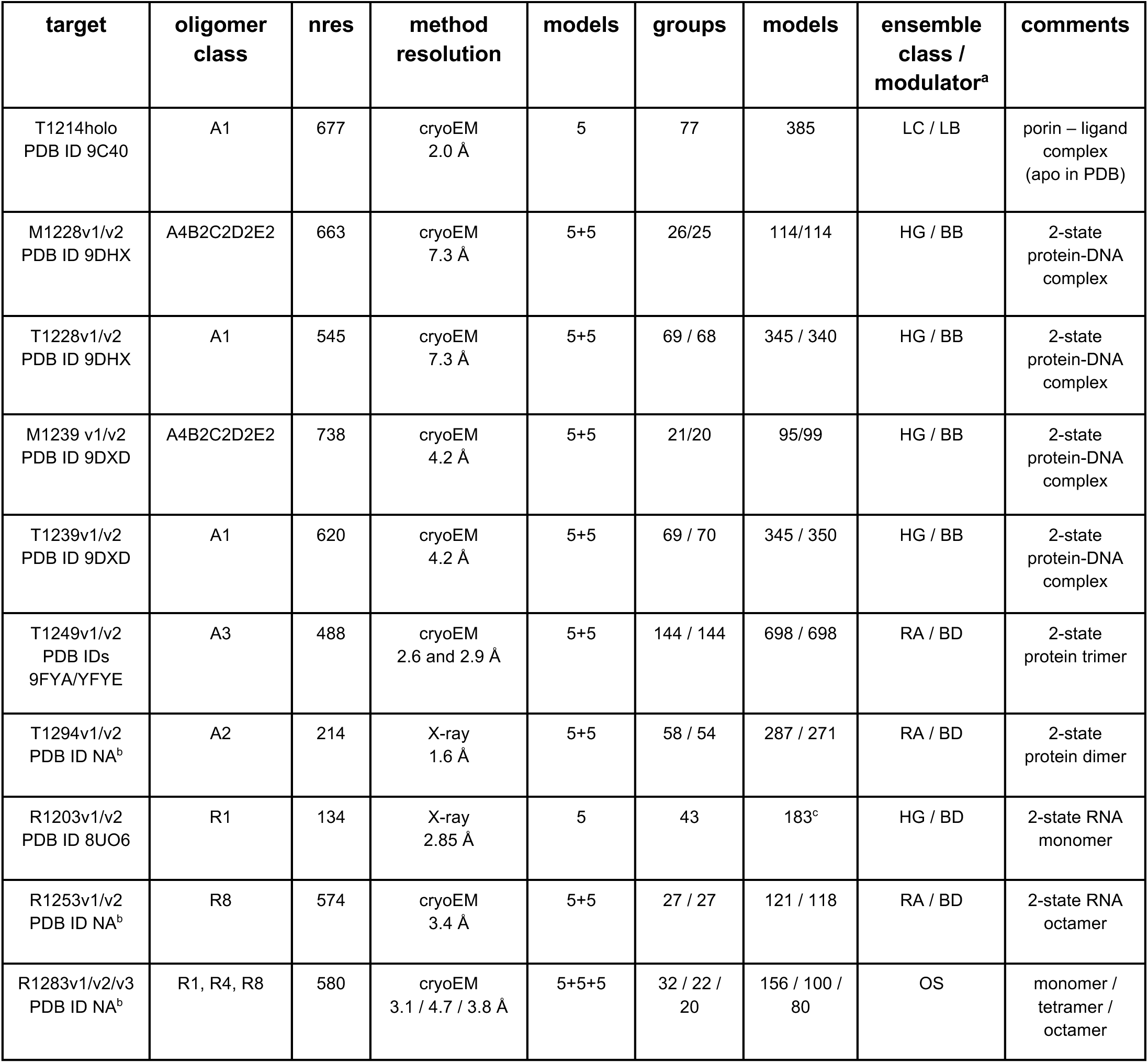
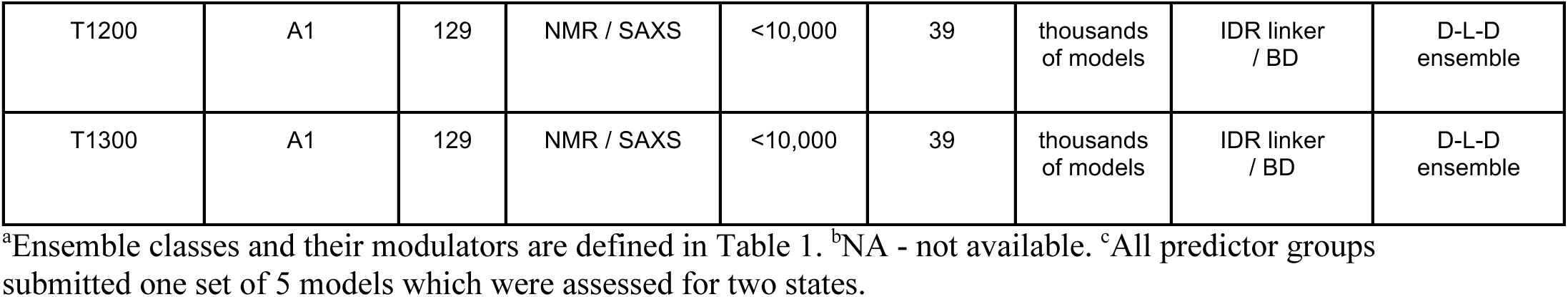
Features of CASP16 Ensemble Targets.

| target | oligomer class | nres | method resolution | models | groups | models | ensemble class / modulator<sup>a</sup> | comments |
| --- | --- | --- | --- | --- | --- | --- | --- | --- |
| T1214holo<br>PDB ID 9C40 | A1 | 677 | cryoEM<br>2.0 Å | 5 | 77 | 385 | LC / LB | porin – ligand complex<br>(apo in PDB) |
| M1228v1/v2<br>PDB ID 9DHX | A4B2C2D2E2 | 663 | cryoEM<br>7.3 Å | 5+5 | 26/25 | 114/114 | HG / BB | 2-state<br>protein-DNA complex |
| T1228v1/v2<br>PDB ID 9DHX | A1 | 545 | cryoEM<br>7.3 Å | 5+5 | 69 / 68 | 345 / 340 | HG / BB | 2-state<br>protein-DNA complex |
| M1239 v1/v2<br>PDB ID 9DXD | A4B2C2D2E2 | 738 | cryoEM<br>4.2 Å | 5+5 | 21/20 | 95/99 | HG / BB | 2-state<br>protein-DNA complex |
| T1239v1/v2<br>PDB ID 9DXD | A1 | 620 | cryoEM<br>4.2 Å | 5+5 | 69 / 70 | 345 / 350 | HG / BB | 2-state<br>protein-DNA complex |
| T1249v1/v2<br>PDB IDs<br>9FYA/YFYE | A3 | 488 | cryoEM<br>2.6 and 2.9 Å | 5+5 | 144 / 144 | 698 / 698 | RA / BD | 2-state<br>protein trimer |
| T1294v1/v2<br>PDB ID NA<sup>b</sup> | A2 | 214 | X-ray<br>1.6 Å | 5+5 | 58 / 54 | 287 / 271 | RA / BD | 2-state<br>protein dimer |
| R1203v1/v2<br>PDB ID 8UO6 | R1 | 134 | X-ray<br>2.85 Å | 5 | 43 | 183<sup>c</sup> | HG / BD | 2-state RNA monomer |
| R1253v1/v2<br>PDB ID NA<sup>b</sup> | R8 | 574 | cryoEM<br>3.4 Å | 5+5 | 27 / 27 | 121 / 118 | RA / BD | 2-state RNA octamer |
| R1283v1/v2/v3<br>PDB ID NA<sup>b</sup> | R1, R4, R8 | 580 | cryoEM<br>3.1 / 4.7 / 3.8 Å | 5+5+5 | 32 / 22 / 20 | 156 / 100 / 80 | OS | monomer / tetramer / octamer |
| T1200 | A1 | 129 | NMR / SAXS | <10,000 | 39 | thousands of models | IDR linker / BD | D-L-D ensemble |
| T1300 | A1 | 129 | NMR / SAXS | <10,000 | 39 | thousands of models | IDR linker / BD | D-L-D ensemble |

<sup>a</sup>Ensemble classes and their modulators are defined in Table 1. <sup>b</sup>NA - not available. <sup>c</sup>All predictor groups submitted one set of 5 models which were assessed for two states.

The overall results of ensemble prediction performance in CASP16 are summarized in **Table 3**. With the exception of target T1214, for which the apo structure was available from the PDB, no prediction groups provided high-accuracy models (e.g., GDT_TS > 0.75) typical of recent CASP experiments. In this study, we used different structural-similarity metrics for the different targets, as appropriate to the target class. “Successful” modeling was defined as correctly modeling the distinct overall fold topologies and/or spatial arrangements of substructures of the multiple conformational states observed experimentally with global TM scores > 0.75. Many groups “successfully” modeled the two alternative conformational states of several target sets. Among these, several groups did about equally well by our assessment metrics. However, no one group was consistently top-tier across all (or even most) targets. This is largely attributable to the wide range of target types, ranging from integral membrane proteins with ligand-induced conformational changes, to multistate protein-nucleic acid complexes, to multiple conformations of a short RNA stem-loop. Accordingly, no effort was made to compare the performance of all groups across all targets.

**Table 3.** Summary of Ensemble Prediction Performance in CASP16.

| Target | States Observed | Best (Score) | Prediction Outcomes and Challenges |
| --- | --- | --- | --- |
| T1214_holo | v1 (holo)<br>v2 (apo) | v1 TM 0.99, GDT_TS 0.96 | Ligand-induced holo state captured with high accuracy by many predictor groups. Templates of ligand bound porins exist with low sequence identity (< 22%). Apo form available to predictors as PDB ID 6V81. |
| M1228 | v1 (trans)<br>v2 (cis) | v1 TM 0.80, GDT_TS 0.37<br>v2 TM 0.81, GDT_TS 0.36 | Overall fold topologies are correct; low GDT_TS and DockQ scores result from domain/interface misalignments. Domain D3 lacked templates, and the relative positions of D3/D4 are poorly modeled. |
| T1228 | v1 (V- shaped)<br>v2 (compact-shaped) | v1 TM 0.83, GDT_TS 0.63<br>v2 TM 0.83, GDT_TS 0.52 | Low GDT_TS reflects domain rotation and orientation errors, especially for domains D3/D4 with hinge-like transitions; v2 (compact-shaped) was least accurate across predictors. |
| M1239 | v1 (cis)<br>v2 (trans) | v1 TM 0.81, GDT_TS 0.30<br>v2 TM 0.68, GDT_TS 0.29 | Overall v2 (trans) arrangement was captured, but v1 (cis) predictions did not model the asymmetric A<sub>2</sub>B<sub>2</sub> tetramer with matching D4-D4 coiled coil interdomain interactions. |
| T1239 | v1 (V-shaped)<br>v2 (compact-shaped) | v1 TM 0.92, GDT_TS 0.72<br>v2 TM 0.77, GDT_TS 0.48 | State v1 (V-shaped) predictions were accurate considering low-resolution of reference structure; v2 (compact-shaped) was least accurate, mainly attributable to inaccurate D4 domain orientation and packing. |
| T1249 | v1 (closed)<br>v2 (open) | v1 TM 0.98, GDT_TS 0.72<br>v2 TM 0.97, GDT_TS 0.48 | TM-scores were high; but only moderate interface recovery for v1 (closed); no groups achieved high <DockQ> for v2 (open). No group excelled in accurately modeling interfaces of both states. |
| R1203 | v1 (open)<br>v2 (closed) | v1 TM 0.64, GDT_TS 0.60<br>v2 TM 0.72, GDT_TS 0.63 | RNA stem-loop modeled, but neither state was accurately predicted due to errors in predicting loop placement and noncanonical base pairs. |

The performance of CASP16 predictor groups on each of the ten ensemble target groups (excluding the D-L-D targets) are summarized in the following sections. Detailed descriptions of methods used by the top performing groups for each target, based on the CASP16 predictor abstracts, are presented as **Supplementary Text**.

### T1214: Ligand-induced conformational change in an integral membrane protein transporter

Target T1214 challenged predictors to model ligand-induced conformational changes that form a cofactor-binding site. This 677-residue *Escherichia coli* TonB-dependent transporter, PqqU, undergoes localized rearrangements in extracellular loops 7 and 8 upon binding the redox cofactor pyrroloquinoline quinone (PQQ), forming a well-ordered β-hairpin that encloses the ligand in an internal cavity (**Figure 1**) [16]. At the time of CASP16, the 2.96 Å X-ray crystal structure of the apo form (PDB ID: 6V81) was available while the ∼2.0 Å holo structure had only recently been determined by cryoEM in the lab of R. Grinter (Monash University, Clayton, Australia) [16]. Predictors were asked to submit five models for the holo state.

**Fig 1.**
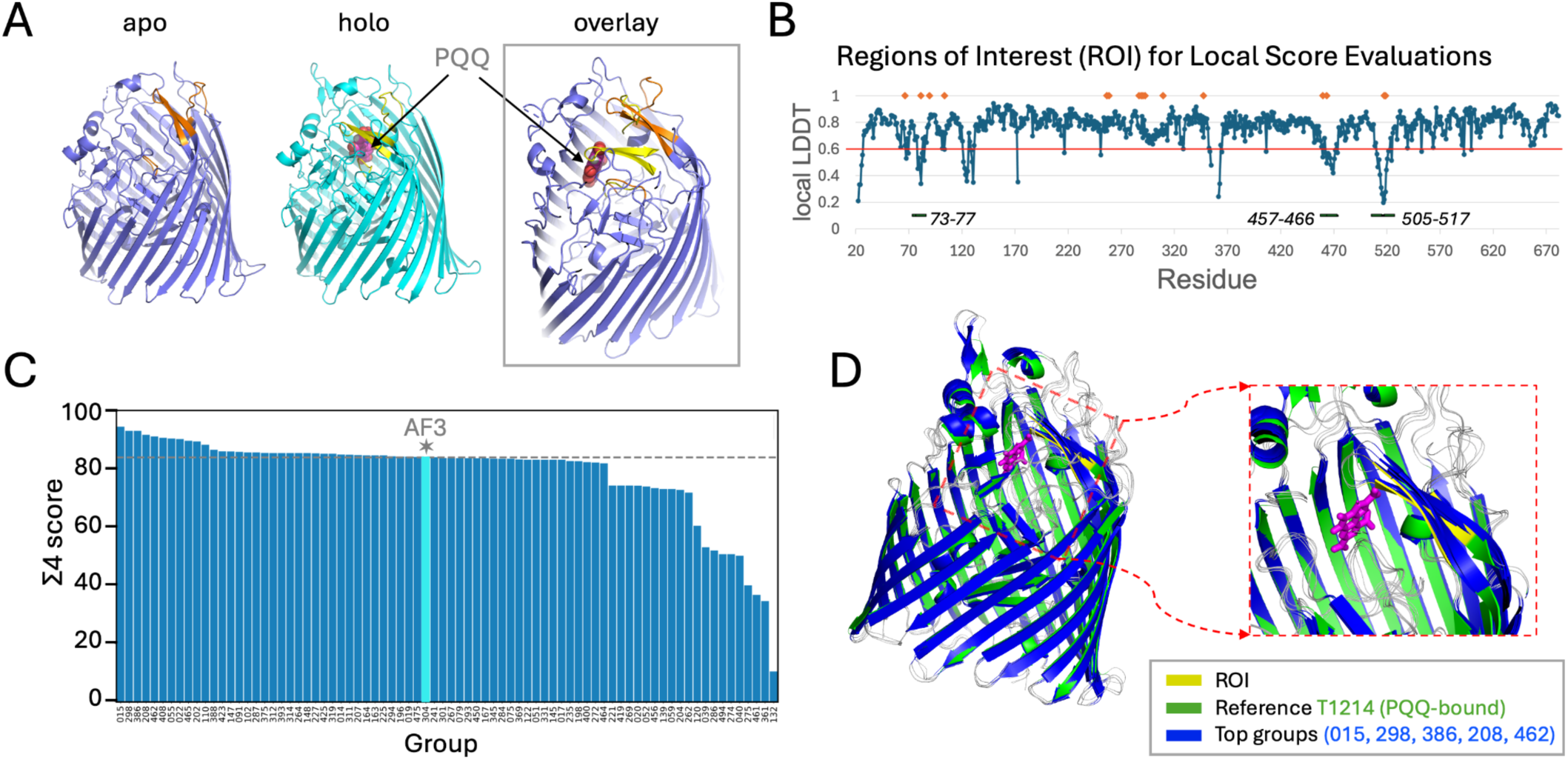
Structural comparison and scoring analysis of T1214_holo models. (A) Target T1214v1 (holo structure) and PDB ID 6V81 (apo structure) with structurally distinguishing ROIs highlighted in orange on the apo structure and yellow on the PQQ-bound structure, with overlay illustrating the key conformational changes upon ligand binding. (B) Per-residue local LDDT plot between apo and holo structures indicating ROIs, identified as segments with at least 3 sequential residues having LDDT < 0.6 and at least one residue < 3.0 Å (orange diamonds) from the PQQ ligand, are indicated with a black horizontal line and residue ranges. (C) Ranking of predictor groups based on the global/local Σ4 score of the best of the five models submitted relative to the reference structure. An expanded version of panel C is presented in **Supplementary Figure S1**. (D) Structural superposition of top-scoring model for each of the top 5 ranked groups onto the T1214 (PQQ-bound) reference (green) with PQQ ligand (magenta). The predicted models are shown in blue, with loops as thin gray lines. Close-up view of the ligand-binding region with ROIs highlighted (yellow) on the reference structure.

Because most of the structure remains unchanged between apo and holo forms, we developed a hybrid global/local scoring approach that emphasizes conformational changes near the PQQ site. Regions of Interest (ROIs) were defined as contiguous segments that both deviated significantly between apo and holo structures and were in proximity to the ligand. These captured the major ligand-induced shifts while excluding distant, poorly defined surface loops unlikely to be functionally relevant (**Fig. 1A-B**). Using these ROI definitions, we computed vaious composite scores Σ1 through Σ4 that combined global (GDT_TS, LDDT, RMSD) and local (LDDT and Cα displacements within ROIs) metrics. Each of these Σ scores, described in the Methods section, ranges from 0 to 100, with 100 indicating a perfect global and local (within ROIs) match between predicted and experimental models. Among the four Σ scores assessed, Σ4 was found to be the most discriminating and was chosen to rank predictor groups (**Fig. 1C-D**).

As illustrated in **Figure 1C**, fifty-seven groups had Σ4 scores of > 80/100 for their best-scoring holo-state model. Eight groups exceeded Σ4 scores of 90 indicating excellent local and global accuracy, and higher than the Σ4 score of 84.17 for AF3-server (group 304). The five top-performing groups were PEZYFoldings (015), ShanghaiTech-Ligand (386), ShanghaiTech-human (298), falcon2 (208), and Zheng (462). A complete ranking of all predictor groups, based on best scoring model per group, is presented in **Supplementary Table S1**. Some groups, such as AF3-server (group 304), ShanghaiTech-human (298), ShanghaiTech-Ligand (386), modeled the ligand explicitly, whereas others, including PEZYFoldings (015), falcon2 (208) and Zheng (462), reported that their predictions were generated without including the ligand.

PEZYFoldings used an AlphaFold2-Multimer-based pipeline augmented with extensive sequence searches, MSA construction, and comprehensive template selection, followed by OpenMM relaxation and a custom rescoring method. Although they did not include the ligand in their reported modeling protocol, they still captured binding-pocket conformations consistent with the holo state, suggesting that MSA-driven context alone provided sufficient information to guide the conformational shift. The available low sequence identity templates (< 22%) include ligand-bound porins that may have impacted the modeling, such as FecA–citrate (PDB 1KMP) and FoxA–Fe-Qercetin (PDB 6I97), which contain similar bent β-hairpin motifs (**Supplemental Table S8**). Overall, the T1214 holo structure was predicted with high accuracy by many groups, showing that ligand-induced conformational changes can be captured even without explicit ligand modeling.

### M1228: Conformational diversity in a DNA-bound multimeric complex

Large serine integrases (LSIs) mediate site-specific DNA recombination reactions that move phage or mobile genetic element DNAs into and out of host bacterial genomes [17]. Target M1228 represents an LSI from *B. subtilis* phage SPbeta that forms a DNA-bound homotetrameric complex composed of four identical protein subunits and four DNA duplexes (2 short and 2 long). The structure was determined by cryoEM in the laboratory of P. Rice (University of Chicago) [18]. Two distinct conformational states of the complex were resolved in the 7.3 Å resolution electron density map, designated M1228v1 (state V1, cis) and M1228v2 (state V2, trans) (**Figure 2A**). These two states differ in their quaternary architecture. In M1228v1, the protein subunits (protomers) 1A and 1B are related to 1A’ and 1B’ by a 180° rotation about the x-axis. In contrast, in M1228v2, protomers 2A and 2B are similarly related to 2A’ and 2B’, but via a 180° rotation about the y-axis (**Figure 2A, Supplementary Figure S2**).

**Fig. 2.**
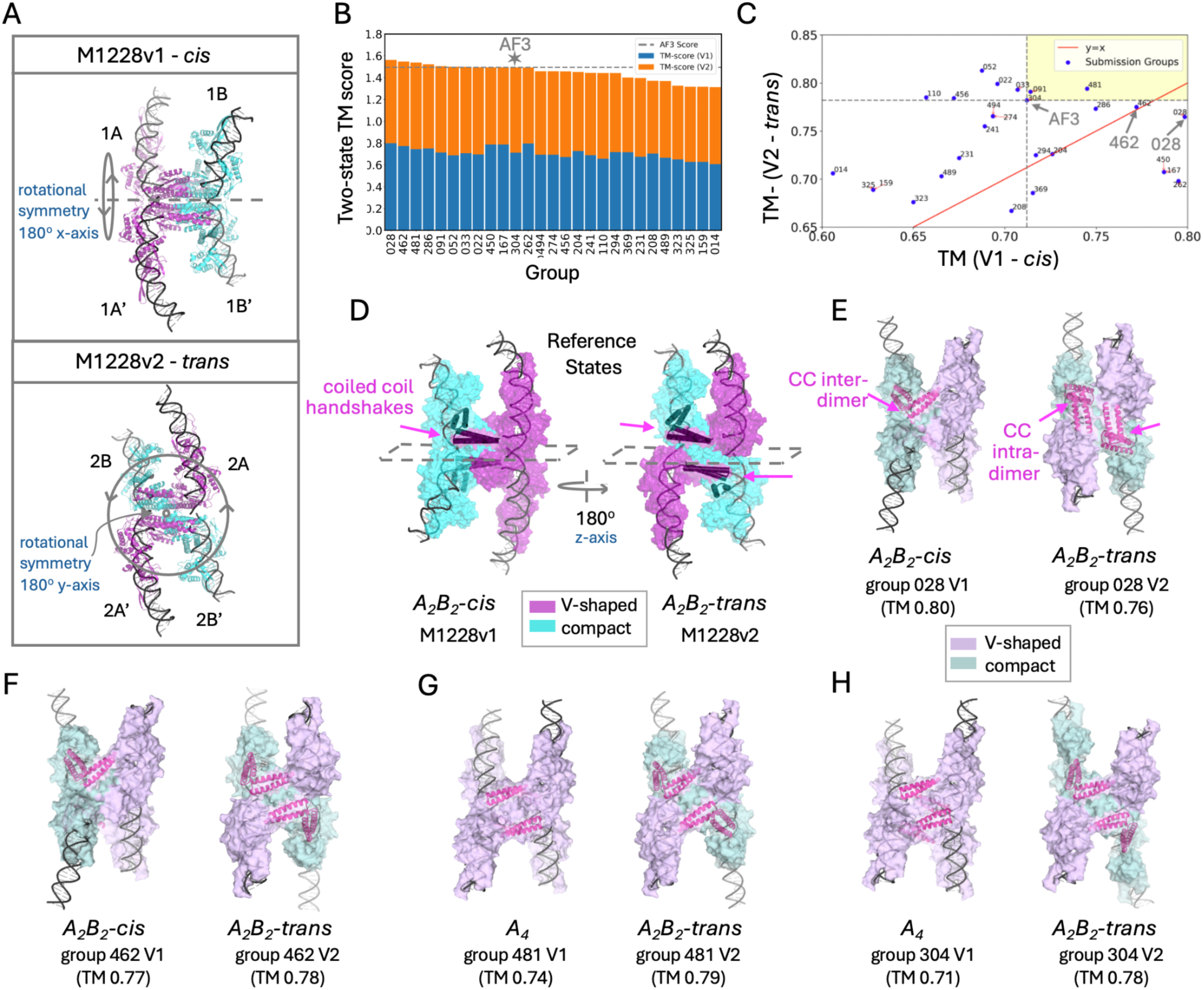
Two-state model evaluation and structural transitions in M1228. (A) CryoEM reference structures M1228v1 (cis) and M1228v2 (trans) represent two quaternary states of the protein-DNA assembly, each containing four protein protomers and four DNA duplexes arranged with distinct rotational symmetries. Protomers are colored by conformation: magenta for V-shaped (A; 1A, 1A’ in V1, and 2A and 2A’ in V2) and cyan for the compact (B; 1B, 1B’ in V1, and 2B, 2B’ in V2). Within each state, primed chains (e.g., 1A′) are related to their unprimed partners (e.g., 1A) by a 180° rotation. (B) Stacked bar chart of two-state performance, ranking of predictor groups by the sum of TM-scores for their best V1 (blue) and V2 (orange) models. AF3-server (304) is indicated by a star and hashed horizontal line. (C) Scatterplot showing each group’s best TM-score to V1 (x-axis) vs V2 (y-axis). Points closer to the upper right of the diagonal line indicate accurate modeling of both states. (D) Quaternary assemblies of V1 and V2, related by a 180° rotation of the bottom half relative to the top half, reconfiguring inter-subunit and protein-DNA interfaces (gray dashed box marks the mid-plane). (E-H) Space-filling representation of the best V1and V2 predictions from groups NKRNA-S (028), Zheng (362), Vfold (481), and AF3-server (304). Protomer state conformational stoichiometry is listed as A_2_B_2_ or A_4_, and protomer conformations are shaded lighter magenta/cyan than in D; Domain 4 helices are shown as pink cartoons, and their coiled coil (CC) handshake interactions are shown. Panels B and C are expanded for readability in **Supplemental Figure S3**.

Specifically, the two alternate assemblies are related by a 180° rotation of the lower half of the assembly relative to the upper half (**Figure 2D**). States V1 (cis) and V2 (trans) can interconvert by such rotation, which is thought to be part of the mechanism of gene recombination by exchanging DNA strands to enable alternate DNA connectivity [18-20]. Each half of the tetramer can be considered an asymmetric dimer, containing one “V-shaped” and one “compact-shaped” protomer (see target T1228 below for further discussion). While the interfaces within each half are conserved, the flat hydrophobic interface between halves allows the lower dimer to swivel relative to the upper dimer (**Figure 2D**). Key structural features illustrated in **Figure 2D** are the protomer conformational stoichiometry, *A*2*B***_2_** (where *italicized* in this designation refers to *conformational stoichiometry*), and the functionally-important interfacial “handshake” interaction between the ends of the antiparallel coiled-coil (CC) motifs of domains D4 of the V-shaped and compact-shaped protomer conformers [18]. In both states, the longer DNA duplexes (33- and 35-base pairs) bind to V-shaped protomers, while the shorter duplexes (24- and 25-base pairs) bind to compact-shaped protomers. These structures are interpreted as evidence for a conformational switching mechanism that reconfigures DNA connectivity like a railway switch [18].

The challenge was to model both the V1 (cis) and V2 (trans) tetrameric assemblies, capturing symmetry, protomer conformations, and the large-scale structural change. FoldSeek [21] confirmed that no close sequence templates (< 20% identity) were available in the PDB. However, a few smaller tetrameric protein–DNA complexes with high symmetry were identified as distant structural analogs, including PDB IDs 1ZR2 and 2GM4 [19,22] (see **Supplementary Table S8** for details). Participants were provided the full 663-residue protein sequence and two duplex DNA sequences and asked to submit up to five models for each state.

For target M1228, as in other CASP ensemble targets, predictors submitted two sets of models (up to five per set) intended to represent distinct conformational states. As the state labels assigned by predictors were arbitrary, we reassigned them by scoring each set against both reference structures using TM-scores. The highest-scoring model across all comparisons determined the initial assignment (e.g., assigning predicted set A to V1); the other predicted set was then assigned to the remaining reference (e.g., set B to V2). The sum of TM-scores from the best model in each set gave “two-state scores” that were used to rank predictor groups (**Methods; Figure 3)**. This procedure can be generalized to more than two reference states when multiple structures are available. TM-score was selected over GDT_TS scores because the highest GDT_TS values for both states were below 0.4, whereas TM-scores exceeded 0.75 for the best models. TM-score is more tolerant of structural variations and better reflects overall fold correctness, while GDT_TS penalizes even small domain movements or misalignments, producing uniformly low values that prohibit identification of correct models for large multi-domain proteins.

**Fig. 3.**
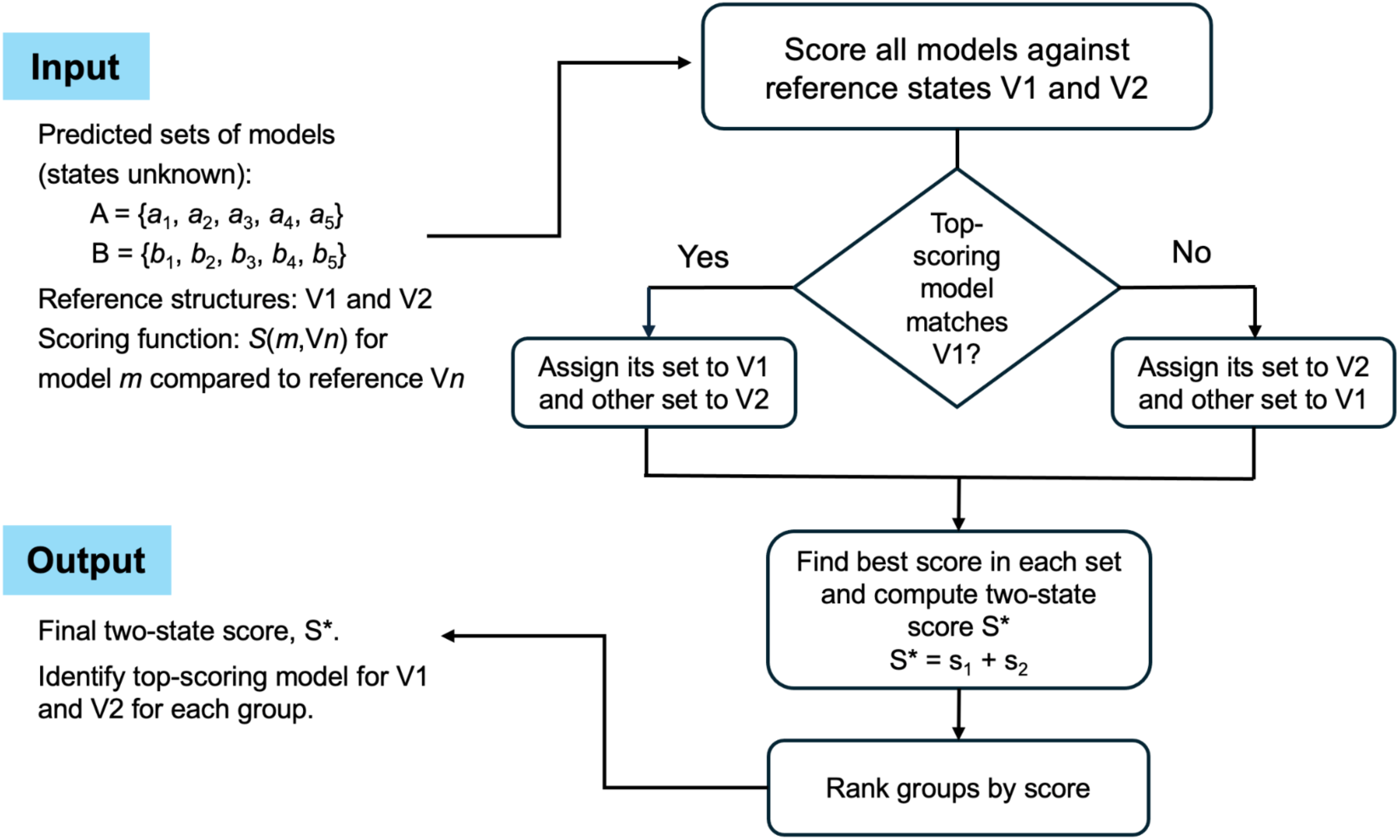
Algorithm for two-state scoring. The algorithm assigns the two predicted ensembles, designated sets A and B, to their best-matching experimental reference states (V1 and V2). The structural scoring functions *S*(*m*, V*n*) comparing predictor model, *m*, to references state V*n*, include TM-score, GDT-TS, DockQ, and composite global/local scores 𝛴 (defined in the Methods), where higher values indicate a better match. Scores s_1_ and s_2_ are from the top-scoring model within each set relative to its assigned reference state.

Based on TM-score analysis (**Figure 2B-C, Supplementary Table S2)**, most groups predicted the V2-trans *A*_2_*B*_2_ conformational state more accurately (14 / 27), with eight capturing inter-dimer coiled coil (CC) handshakes between D4 domains, including groups Zheng (462), Vfold (481), and AF3-server (304) (shown in **Figure 4F-G, right**). Fewer groups predicted the V1-cis *A*_2_*B*_2_ state; only four groups did so correctly, including NKRNA-s (028) and Zheng (462) (**Figures 2E–F, left**). However, NKRNA-s predicted an intra-dimer CC handshake (between protomers along the non-contiguous DNA chain), whereas Zheng reproduced the experimental inter-dimer handshake. Many groups (17) produced symmetric *A*_4_ models (with all V-shaped protomers) that have antiparallel CC interactions that are not consistent with the reference structures (e.g., **Figure 2G–H, left**).

**Fig. 4.**
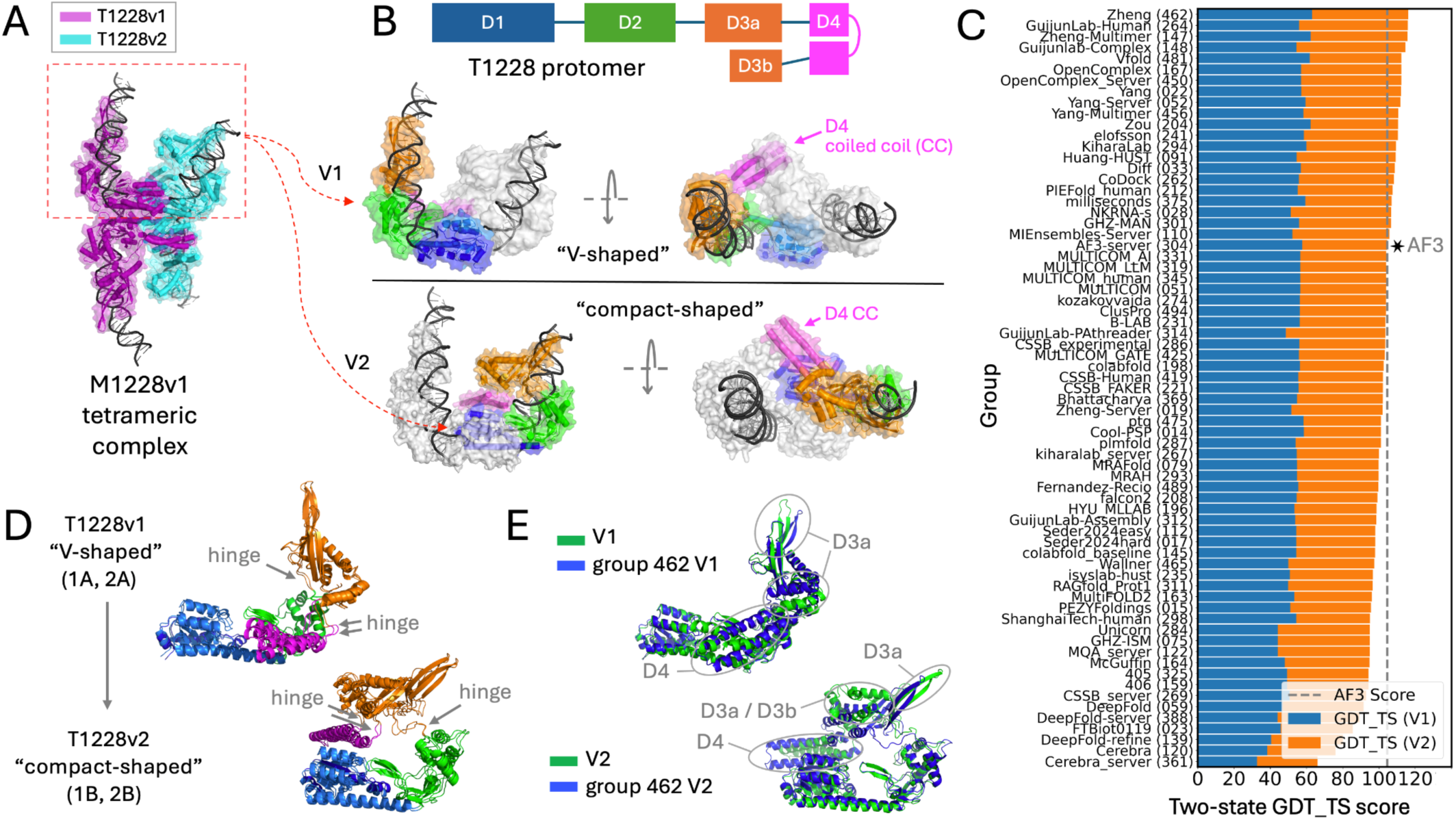
Two-state structural analysis and model evaluation for target T1228. (A) Reference structures T1228v1 (magenta) and T1228v2 (cyan) represent two distinct subunit conformations of the M1228v1 tetramer bound to two strands of dsDNA. (B) Domain organization of target T1228 along the primary sequence: D1 (blue), D2 (green), D3a-b (orange), and D4 (pink). Protomer subunits of M1228v1 (top half) are shown colored by domain for T1228 state V1, “V-shaped”, (top) and state V2, “compact shaped” (bottom). (C) Combined two-state GDT_TS scores for the best-performing models from each group. The AF3-server (304) result is indicated by a star and vertical dashed line. (D) The T1228v1 “V-shaped” subunits (1A, M1228v1; 2A, M1228v2) are highly similar (global LDDT = 0.93), as are the T1228v2 “compact-shaped” subunits (1B, M1228v1; 2B, M1228v2; LDDT = 0.87). For evaluation, chain 1A (M1228v1) was used as the reference state T1228v1 (V1) and chain 1B (M1228v1) for state T1228v2 (V2), given their close similarity to these corresponding protomers in the same state. The large conformational change between T1228v1 and v2 involves rotation about hinge regions (arrows). (E) Comparison of Zheng group 462 models (blue) with the reference states (green) V1 and V2 of T1228. Panel C and **t**he two state-scatter plot of GDT_TS scores for the best V1 and V2 models submitted by each group are shown in larger format in **Supplemental Figure S4**.

The best-scoring two-state ensembles (based on rankings shown in **Figure 2B**) came from groups NKRNA-s (028), Zheng (462) and Vfold (481). These two-state ensembles, each with two-state scores slightly higher than AF3, are shown in space filling representation in **Figure 2E-G**. Other groups achieved similar two-state performance (≥ 1.49), including Huang-HUST (091), Yang-Server (052), Diff (033), Yang (022), OpenComplex_server (450), OpenComplex (167), AF3-server (304), and CoDock (262). CoDock and Yang-Server achieved the best scores for one state, but not the alternate, lowering their combined scores. Modeling of the DNA duplexes was not formally assessed, since DNA placement was poor: no predictors reproduced the correct binding of long duplexes to V-shaped protomers and short duplexes to compact ones (**Figure 2E-H**). Despite reasonably high TM-scores, the relatively low Global DockQ scores (< 0.37) indicate low accuracy of these models and confirm poor DNA-protein interface packing in general (**Supplementary Figure S3D-F).**

Overall, the Zheng (462) group had the best performance with target M1228. They employed a symmetry-aware modeling strategy incorporating deep learning-based docking to recover both cis and trans quaternary arrangement and conformational stoichiometry (*A*_2_*B*_2_), details of the CC handshake, and even recapitulated DNA chain swapping behavior. However, protein-DNA interactions were not positioned correctly, and the short and long DNA segments were bound to the incorrect protomer states. Group NKRNA-s (028) had the (marginally) best performance based on two-state TM scores, but did not provide correct details for the CC handshake. They developed DeepProtNA for modeling of protein-DNA and protein-RNA complexes. MSAs were built using large metagenomic databases, and structural templates were obtained using methods which can process complex assemblies. AlphaFold2 variants with extended decoy generation and different dropout settings were used for modeling. The best two-state scoring models of other top performing groups (e.g. Vfold (481) and AF3-server (304)) do not have correct conformational stoichiometry (*A*_2_*B*_2_) for state V1. The top-performing server groups were Yang-Server (052), OpenComplex_server (450), and AF3-server (304). Additional details of the methods used by these and other top-performing groups are summarized in **Supplementary Text**.

### T1228 Protomer States

Target T1228 is the 545-amino acid single-chain subunit of the M1228 complex, which consists of four such protomers bound to DNA (**Figure 4A**). Each T1228 protomer contains four domains (D1 - D4; D3 comprises two segments, D3a and D3b) (**Figure 4A,B)** and adopts two pairs of distinct conformational states within the assembly: a “V-shaped” state (T1228v1, state V1) and a “compact shaped” state (T1228v2, state V2).

The two states differ by domain rotations about hinge regions in interdomain linkers (**Figure 4B,D**). A key modeling challenge was the lack of close structural homologs: none of the available templates for domains D1–D4 had more than ∼ 20% sequence identity. The best alignments were for domain D1, including smaller tetrameric DNA-bound complexes related to M1228. Additional templates, such as PDB IDs 4KIS and 6DNW, covered D2 and the first part of D3 (DNA-binding region). These resemble state V1 more closely than V2, but are not dimeric or tetrameric. By contrast, D4 templates were restricted to short alignments with coiled-coil containing proteins (**Supplementary Table S8**). These biases likely explain why most predictors, including AF3-server (304), modeled V1 better than V2 (**Supplementary Figure S4, bottom**). In particular, in V2 predictions domain D4 was well folded but often mispositioned within the complex.

As discussed above, within M1228 each asymmetric dimer pairs one V-shaped and one compact-shaped protomer. The absence of homologous templates for the remainder of D3, together with hinge flexibility, made accurate placement of D3/D4 especially difficult to model. As a result, the GDT_TS scores were modest for both conformers: all V2 state predictions had GDT_TS < 60, and only three groups exceeded 60 for V1, with Zheng (462) highest at 63. These moderate values indicate reasonable accuracy given the low resolution (7.3 Å) of the cryoEM reference structure, but are lower than typically observed for single-state large multi-domain proteins in recent CASP studies using AI-based methods.

As illustrated in **Figure 4C** and **Supplementary Table S3**, **t**he top five T1228 predictor groups were Zheng (462), GuijunLab-Human (264), Zheng-Multimer (147), Guijunlab-Complex (148), and Vfold (481). For Zheng and others, a persistent limitation was accurate modeling of the D3/D4 region (**Figure 4E**). Several groups had two-state scores higher than AlphaFold3-server (304). Interestingly, although group NKRNA-s (028) had the highest two-state TM-score for M1228, they predicted a different D4 orientation of the V-shaped state V1, which is reflected in the lower GDT_TS score ranking of only 19th place in the two-state GDT_TS scoring for T1228v1. The best scoring server groups included GuijunLab_Server (450), OpenComplex_Server (167), Yang_Server (052), and MIEnsembles-Server (110), all of which had GDT_TS two-state scores (marginally) higher than AlphaFold3-server (304).

Overall, the best-performing group for T1228 was Zhang (262). Although many more groups (69) submitted multistate T1228 predictions than M1228 (27), groups Zheng (462) and Vfold (481) were the two top-ranked groups for T1228 that also submitted predictions for M1228. The methods used by Zhang (262) are outlined above. Vfold (481) used AlphaFold3 to predict structural candidates, d AMBER-based energy minimization for refinement, and a knowledge-based scoring function to evaluate the quality of models. GuijunLab-Human (264) and Guijunlab-Complex (148), which also performed well for T1228, utilized a strategy focused on paired MSA construction using deep learning, together with AlphaFold-Multimer and DeepAssembly [23] to generate a series of multimer structures. These were then ranked using in-house multi-model quality assessment methods. GuijunLab-Human (264) used AlphaFold2, AlphaFold3, and HDOCK [24] to predict monomer and complex structure. These models were then used to determine inter-chain distances which were input into DeepAssembly [23] to predict the complex structures. Additional details of the methods used by these and other top-performing groups are summarized in **Supplementary Text**.

### M1239 and T1239: Conformational diversity in a second DNA-bound multimeric complex

Target M1239 is a second *B. subtilis* phage SPβ LSI homotetrameric protein-DNA complex provided for CASP16 by the P. Rice laboratory (University of Chicago) [18]. Each 622-residue subunit is homologous to those of M1228, but includes an additional 85 C-terminal residues. The cryoEM structure (4.2 Å, PDB 9DXD) revealed two conformational states, M1239v1 (state V1, trans) and M1239v2 (state V2, cis), resembling those of M1228 but modified by the longer C-terminus (**Figure 5A,B, Supplementary Fig. S5A**). Specifically, 57 of the additional residues form a third non-contiguous segment (D3c), enlarging domain D3 relative to M1228. In the “compact-shaped” protomer, D4 extends further to make additional contacts in the minor groove of a second DNA duplex (**Figure 5B**). Aside from these additional DNA-binding interactions, binding of the longer DNA duplex to the V-shaped (V1) protomer and shorter DNA duplex to the compact-shaped (V2) protomer were similar to the M1228 complex. The tetramer-stabilizing inter-dimer D4-D4 CC handshake between the upper and lower asymmetric protomers was retained in both states (**Figure 5A**) and is important for stabilization of the active tetramer over the inactive dimer [25].

**Fig. 5.**
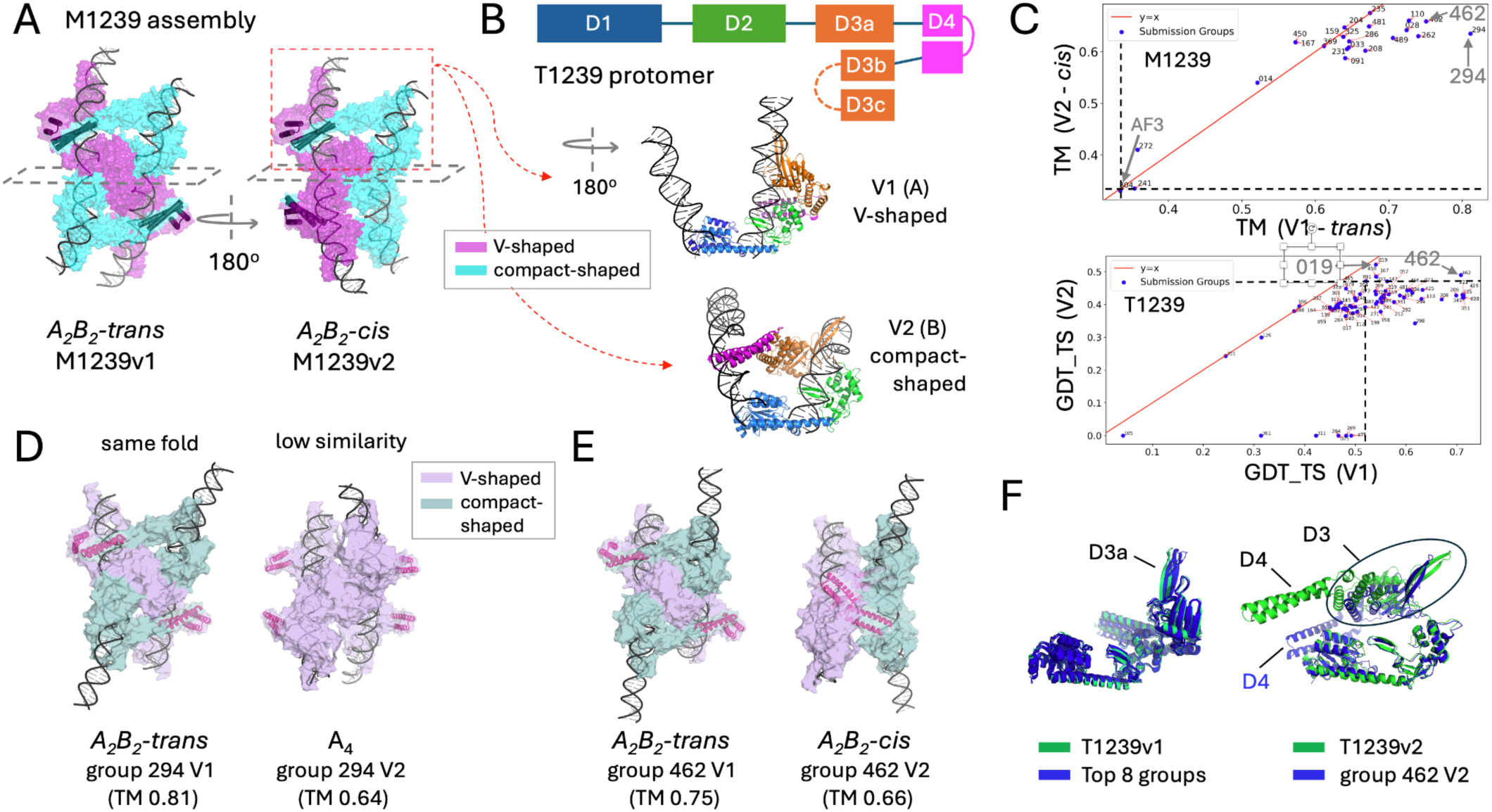
Conformational states of M1239 and T1239 and two-state structure analysis for both. (A) Reference structures M1239v1 and v2, representing two distinct conformational states V1 (trans) and V2 (cis). (B) Top: T1239 schematic with domain color coding used in structural representations: domains D1 (blue), D2 (green), D3a-c (orange), and D4 (pink). Dashed region in D3 indicates residues missing in the electron density (residue range 536 - 564). Bottom: Structural comparison of T1239 references states: V1 V-shaped and V2 compact-shaped. (C) Scatterplots of two-state TM-scores for M1239v1/v2 and two-state GDT_TS scores for T1239v1/v2. (D) Comparison of top-scoring KiharaLab (294) models to each reference state. Even for the best-scoring models, similarity to M1239 V2 was low (TM = 0.64). (E) Comparison of group Zheng (462) models to each reference state. Similarity to M1239 V2 was also low. (F) Superpositions of predicted models from eight top-scoring groups for V1 (blue, left) and group 462 for V2 (blue, right) onto reference states (green). Plots from panel C are shown at larger scale, along with the corresponding M1239 TM-score and T1239 GDT_TS plots, in **Supplemental Figures S6D and S7D**.

As with M1228, for M1239 predictors submitted five models per state and accuracy was assessed by two-state TM-scoring (**Figure 5C top, Supplementary Figure S6**, **Supplementary Table S4)**. The top performer based on two-state TM scores, KiharaLab (294), predicted the tetrameric V1 trans topology well (TM 0.81) but had incorrect conformational stoichiometry (*A*4) for the V2 cis state (TM 0.64) (**Figure 5D**). Other high-ranking groups, including Zheng (462), MIEnsembles-Server (110), CoDock (262), and NKRNA-s (028), also matched V1 with modestly high TM scores (> 0.70), reproducing the asymmetric dimer arrangement, *A*_2_*B*_2_ conformational stoichiometry, compact protomer - DNA contacts, and inter-dimer CC handshakes. For the V2 state only Zheng (462) recapitulated the cis architecture with correct *A*_2_*B*_2_ protomer state stoichiometry. However, their CC handshake was not modeled between the same protomers as in the experimental structure (**Figure 5E, right**). Most predictor groups generated four symmetric protomer subunits for V2 instead of the correct *A*_2_*B*_2_ conformational stoichiometry, either all V-shaped (*A*_4_) as shown for KiharaLab (**Figure 5D, right**) or all compact-shaped (*B*_4_), lacking CC handshakes and distal DNA–minor groove contacts. For all groups, the best models for both V1 and V2 had very low GDT_TS scores (< 32 and < 30, respectively) and global DockQ scores (<0.40) (**Supplementary Fig. S5C–F**), reflecting substantial discrepancies in protein–DNA and protomer–protomer interfaces.

To further evaluate intra-protomer domain orientations, we analyzed the two-state target T1239, which had broader participation (76 groups vs. 21 for M1239). T1239 mirrors the T1228 architecture for domains D1–D4 but includes an added D3c segment and a flexible 29-residue D3b–D3c loop not resolved in the cryoEM map (**Figure 5A,B**). As in T1228, protomers adopt either a V-shaped (V1) or compact-shaped (V2) conformation. Two-state GDT_TS scoring (S**upplementary Figure S7 and Table S5**) revealed the same trend as for T1228: generally accurate predictions of V1 (GDT_TS > 0.7) by Zheng (462), isyslab-hust (235), CSSB_FAKER (221), CSSB-Human (419), NKRNA-s (028), MULTICOM_human (345), and MULTICOM (051)), but poor agreement (GDT_TS < 0.5) with V2 due to widespread D3/D4 misplacements. Zheng achieved the highest two-state GDT_TS, with only minor D3/D4 deviations in V1 but larger mismatches for the compact V2 protomer (**Figure 5F**). Details of the methods used by these top-performing groups are summarized in **Supplementary Text**.

Overall, most groups failed to predict the trans-to-cis tetrameric transition of M1239, instead producing symmetric tetramers as the second state. The absence of inter-dimer CC handshakes likely contributed to inaccurate D3/D4 placement in the T1239 protomers. Even for the highest scoring models, the functionally important inter-dimer CC contacts do not match the experimental structures. As detailed in the accompanying CASP Special Issue paper [25], modeling of DNA-binding domain interactions in this active tetramer was often inaccurate and was not directly evaluated in our study. AF3-server (304) predictions were dimers, not tetramers, and are therefore not a suitable baseline for evaluating tetramer predictions. Some predicted tetramers did not match either experimental state, potentially reflecting alternative assemblies rather than the two that have been observed experimentally. In conclusion, the M1239/T1239 studies further underscore the difficulty of modeling flexible state transitions with large-scale interdomain motions coupled to multi-assembly DNA interactions.

### T1249: Ligand-induced interface switching in a trimeric protein complex

Target T1249 is a viral spike protein complex from the Sabiá virus (SABV), a Clade-B arenavirus that causes Brazilian hemorrhagic fever. CryoEM structures solved at 2.6 Å (closed) and 2.9 Å (open) capture the native state and the conformation adopted during cell entry. These targets were provided for CASP16 by R. Diskin (Weizmann Institute of Science) [26]. The trimeric complex consists of three identical 488-residue monomers; however, the reference coordinates and sequence provided to predictors corresponded only to the cryoEM–modeled region (residues 59–415, A3). The compact, tightly packed closed state (V1) contrasts with the open state (V2), which was induced by acidic pH together with tight binding to an unidentified metal ion [26]. In the trimer, three His157 side chains, one from each protomer, coordinate this metal ion, shifting the equilibrium toward the open state and likely stabilizing it during cell entry. The trimeric conformers differ primarily in the packing interfaces between otherwise identical protomers, indicating that the conformational change is a concerted rigid-body rearrangement with a substantial impact on inter-protomer packing (**Figure 6A**). Although homologous spike protein templates were identified by FoldSeek, high-confidence matches (50–60% sequence identity) were limited to a central region of T1249. Templates identified for the N-terminal ∼180 residues, which includes the H157 metal-binding site that modulates the state equilibrium, shared only low sequence identity (< 20 %) (**Supplementary Table S8**). Many of the same templates were identified for both V1 and V2, but most had substantially higher scores for V1 (Δbits > 100; **Supplementary Table 8**), indicating better matches to this state. This likely contributed to the generally higher accuracy of V1 predictions compared to V2.

**Fig. 6.**
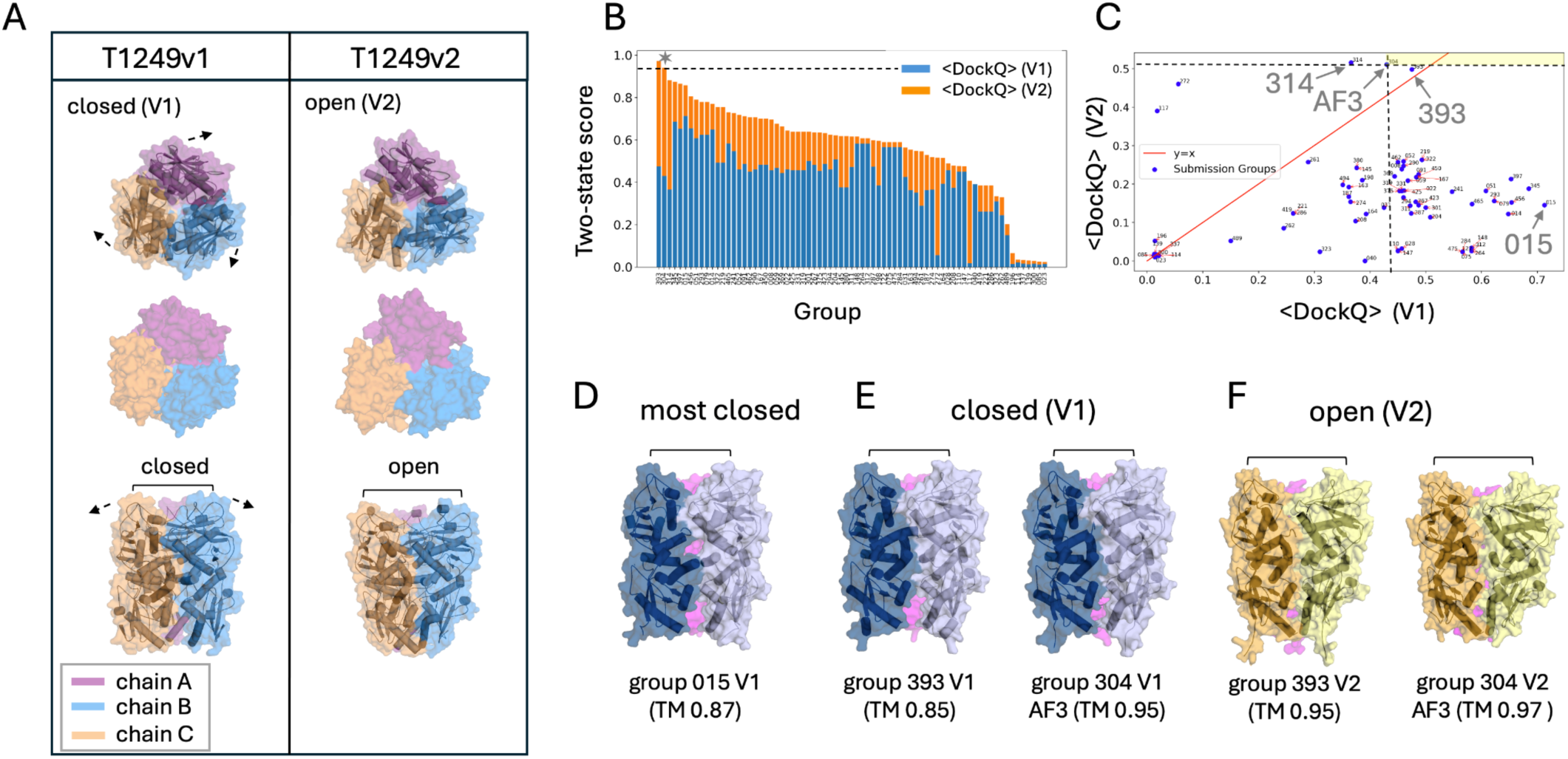
Two-state structure and interface scoring of homotrimeric T1249 models. (A) Comparison of closed (V1) and open (V2) states, shown in ribbon and surface views, with chains of the trimers colored distinctly. (B) Two-state DockQ scores averaged across the three interfaces for the best model of each group, sorted by performance. The performance of AF3-server (304) is indicated by a star and horizontal hashed line. (C) Scatterplot comparing <DOCKQ> scores for states V1 and V2. (D) The top scoring closed state model is from group PEZYFoldings (015). (E) Best closed (V1) structures from groups GuijunLab-QA (393) and AF3-server (304). (F) Same as panel E for open (V2) structures. See **Supplemental Figure S8** for expanded versions of panels B and C.

Predictors submitted five models for each state. Many groups provided models with high TM scores for both states (> 0.85), reflecting the trimer topology and similarity in the monomer folds for most targets. By contrast, DockQ scores [27], which evaluate inter-protomer contacts, provided a more sensitive measure of the biologically relevant conformational change. Therefore, assessment was done using the two-state DockQ score averaged across the three protomer–protomer interfaces, (<DOCKQ>). Scores range from 0 (poor) to 1 (perfect interface match). A ranked list of two-state <DOCKQ> scores for all groups is provided in **Supplementary Table S6**. As shown in **Figure 6B** and **Supplementary Figure S8,** no group excelled with both states, but several produced models with moderate agreement to the V1 state. While most groups struggled to accurately model the packing of the V2 state, the three top-scoring groups GuijunLab-QA (393), AF3-server (304) and GuijunLab-PAthreader (314) had relatively balanced performance for both V1 and V2, albeit moderate (< 0.5) <DOCKQ> scores (**Figure 6C**).

Notably, these three groups also had the best two-state TM-scores (> 0.95 for both V1 and V2). Group PEZYFoldings (015) performed best on V1 <DOCKQ> scores (∼0.7) and captured the wider interface (**Figure 6D**), but scored poorly for V2, while group GuijunLab-PAthreader (314) had best agreement with state V2. Notably, groups GuijunLab-QA (393) and AF3-server (304) recapitulated the positioning of H157 side chains in V2, poised for metal coordination, consistent with the experimental open-state structure, while in their V1 models this feature was absent, consistent with the experimental closed-state structure. Structural comparisons and averaged DockQ scores (all < 0.50) illustrate that even the best-performing groups recovered interface details of both states with only limited accuracy (**Figure 6C, E,F)**.

The five top-performing groups for T1249 based on two-state average interface <DOCKQ> scores were GuijunLab-QA (393), AF3-server (304), GuijunLab-PAthreader (314), MULTICOM_Human (345), and smg.ulaval (397). Among these, GuijunLab-PAthreader (314), GuijunLab-QA (393), AF3-server (304), and MULTICOM_Human (345) reported careful handling of trimeric protein units, albeit with different emphases. GuijunLab groups focused heavily on model quality assessment and structure-guided MSA enhancement. AF3-server prioritized efficient stoichiometry prediction with practical interventions for large systems, while MULTICOM_Human (345) excelled in massive model generation and quality-ranking frameworks combining AI and human expertise. No abstract or methodological details were provided for the smg_ulaval group. Common strategies included using AlphaFold2/3, enhanced MSA protocols, and model quality ranking using pLDDT and interface-focused metrics. Differences lay in the degree of manual intervention, and the integration of structural clustering and deep learning-based MQA models. Additional details of these methods are outlined in the **Supplementary Text**.

### R1203: HIV-1 Rev Response Element Stem-Loop II (SLII) conformational switching

Target R1203, provided by K. Choi (Univ. of Texas), is a 134-nucleotide RNA monomer derived from the HIV-1 Rev Response Element (RRE) stem-loop II (SLII). SLII is critical for the viral life cycle as it facilitates Rev-mediated nuclear export of incompletely spliced viral transcripts [28]. The structure was determined by X-ray crystallography at 2.85 Å resolution, and the asymmetric unit revealed two distinct conformations: an “open” state with an expanded major groove, and a “closed” state with more stabilized loops. These states differ primarily in the number and position of non-canonical base pairs (**Figure 7A**) [29]. In the open state, the three-way junction is thought to facilitate initial high-affinity Rev binding at the unpaired base U65 in the major groove, which then promotes subsequent cooperative binding and oligomerization.

**Fig. 7.**
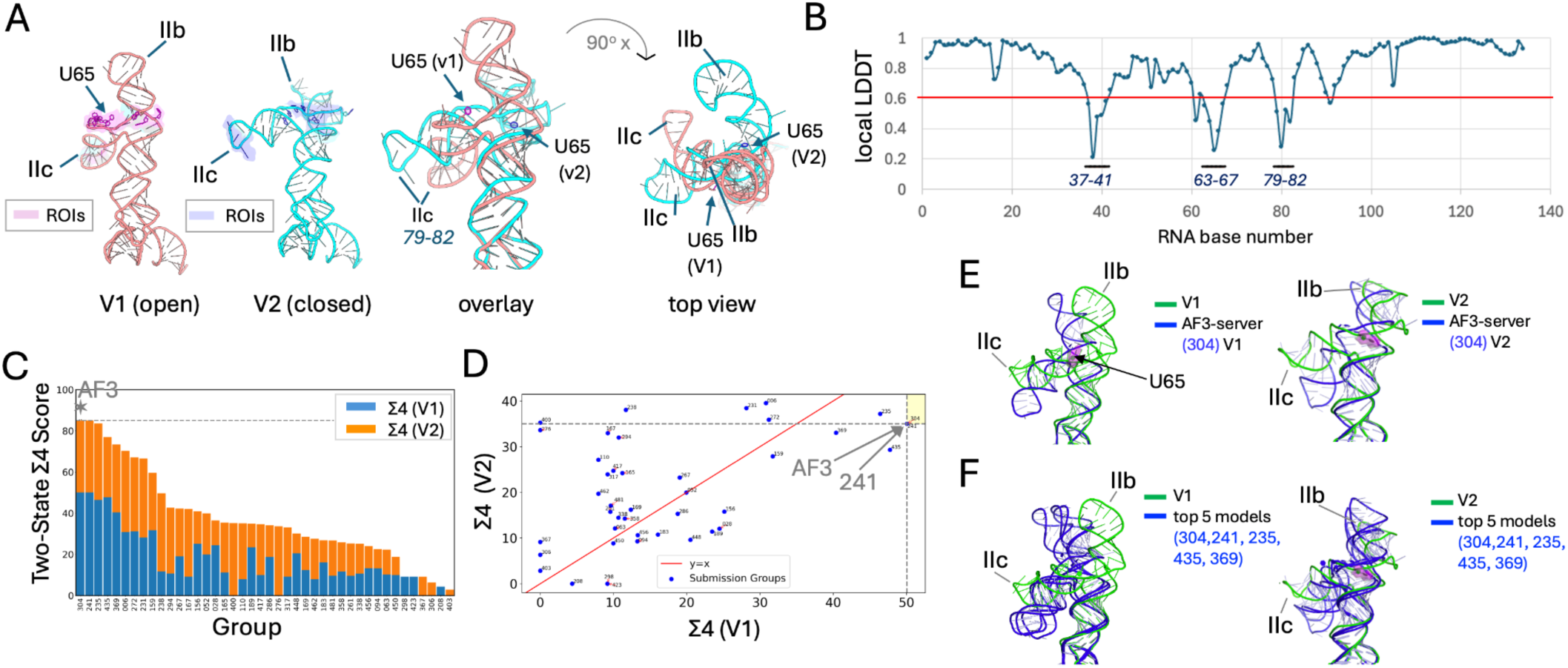
Two-state analysis of RNA models for target R1203. (A) Structural comparison of open (V1) and closed (V2) conformational states of R1203, highlighting loop regions IIb and IIc and superpositions from two different views. The unpaired nucleotide U65 in the open major groove is indicated. (B) Per-nucleotide local LDDT profile for V1 compared to V2 indicating the largest structural differences between states. ROI’s (local LDDT < 0.6) regions were used in the Σ4 score and are highlighted in panels A (magenta) and B (blue). (C) Two-state Σ4 scores for each group’s best models. (D) Scatter plot of Σ4 scores for V1 and V2 across top-scoring group models. (E) Structural alignment AF3-server (304) models with V1 and V2. (F) Alignment of the best five Σ4 scoring models from each group to V1 or V2, showing loop rearrangements and the poor local agreement for the predictions. Panels C and D are shown in larger format in **Supplemental Figure S9**.

CASP16 predictors submitted five models, and each was evaluated against both the open (V1) and closed (V2) conformations observed experimentally. As with T1214, predictors were scored using the two-state Σ4 metric, which combines global and local accuracy, with the local component focused on ROIs that differ most significantly between states. These ROIs, highlighted in **Figure 7A,B**, correspond to loop and groove segments implicated in Rev binding. Score distributions showed a small cluster of groups that modestly outperformed the rest (**Figure 7C,D, Supplementary Table S9**). The top five groups, AF3-server (304), elofsson (241), isyslab-hust (235), RNAFOLDX (435) and Bhattacharya (369) had higher scores for state V1 than for V2. In more than half the cases, the same model would have ranked highest for both states if it had been allowed (data not shown), indicating that predictors struggled to distinguish between open and closed states. No submission exceeded a Σ4 score of 51 for V1 or 40 for V2, indicating overall poor performance. The top TM-scores for each state were 0.64 and 0.72, values that indicate the overall fold was captured but fall short of the > 0.75 threshold typical of high-accuracy models and defined here as "successful". The IIb loop and exposure of unpaired U65 were consistently mispredicted, and even the top-scoring group, AF3-server, failed to reproduce the IIb loop orientation or non-canonical base pairing in either conformation (**Figure 7E,F**). Overall, R1203 was especially challenging, revealing limitations in current RNA prediction methods for modeling conformational transitions, three-way junctions, and non-canonical base pairs, as noted in other CASP16 RNA target evaluations [30].

While no groups were "successful" - based on our criteria defined above - with this target, the top-performing groups were AF3-server and elofsson (241) using human-refined AF3 predictions. Most other top-performing groups built on AlphaFold2, but did not perform quite as well as the AlphaFold3-server. RNAFOLDX (435) utilized a distinct convolutional neural network for modeling, with performance rivaling AF3-server.

### Unsuccessfully modeled targets

Despite the successful (five cases) and promising but unsuccessful (two other cases) results in predicting structures of the two alternative states observed experimentally for the 7 protein-ligand, protein-DNA, trimeric protein, and RNA targets outlined above, CASP16 predictors were unsuccessful in modeling structures of 3 additional target sets, *viz* T1294v1/v2, R1253v1/v2, and R1283v1/v2/v3. Target T1294, a homodimeric isocyanide hydratase protein, was provided with two structural states (V1 and V2), but careful visualization revealed only subtle differences between them, with minimal domain-level rearrangement. The assessors’ consensus was that these structural transitions were too minor to represent meaningful alternative conformations. Consequently, this target was excluded from detailed assessment. Most predictor submissions showed near-identical models for both states, and no group successfully distinguished or justified a biologically relevant structural transition. RNA targets R1253 and R1283 were also very challenging. These were large RNA molecules (574 and 580 nucleotides respectively) provided in multiple oligomeric states monomer (A1), tetramer (A4), and octamer (A8)-requiring accurate modeling of RNA tertiary structure as well as quaternary assembly, often in the absence of homologous templates or well-characterized structural motifs. Across all groups, predictions failed to recover global fold, domain architecture, or base-pairing pattern in any of the states. The highest global GDT_TS scores for monomeric forms were below 20, and for higher-order assemblies, scores dropped below 1. Superpositions with experimental structures revealed extensive misfolding and poor modeling of key structural junctions and tertiary contacts. These outcomes reflect the current limitations of RNA modeling approaches, especially when faced with large, flexible RNAs undergoing oligomerization. No participating method succeeded in capturing the large-scale structural reorganization required for tetrameric or octameric assembly. These findings are consistent with broader trends observed in the CASP16 RNA structure assessment [30], highlighting the need for more advanced tools capable of handling RNA multimerization and conformational heterogeneity. Within this context, R1253 and R1283 were among the most challenging RNA targets in this experiment.

## Discussion

Overall, CASP16 predictors were “successful” in modeling key structural features of both alternative conformational states for five of these ten CASP16 ensemble modeling targets. This includes target T1214, for which the structure of one state (apo) was available in the PDB, and predictors were challenged to model structural details of a second (holo) state. However, no one group dominated in these performance rankings across all targets. This is attributable to the fact that the successful targets spanned a wide range of target types, including ligand binding to an integral membrane protein (target T1214_holo), a large and complex tetrameric protein-dsDNA complex (target M1228), the corresponding subunits of protein-dsDNA complexes (targets T1228 and T1239), and a trimeric protein (T1249). Predictors were not successful in predicting the tetrameric protein-dsDNA complexes (target M1239), the observed oligomerization states of protein target T1294, the two conformations with specific base-pairing of the RNA stem loop (R1203), or the structural ensembles reported for RNA targets R1253 and R1283. In the case of the M1239 protein-DNA complex, successful modeling was challenged by long-range interactions between the longer D4 domain and dsDNA.

TM-score and GDT-TS V1 vs V2 scatter plots for several of the most-successfully modeled targets are shown in **Supplementary Figure S11**. These plots illustrate the range of TM- and GDT_TS scores obtained for the most successful target pairs, excluding target T1214 for which only one state (holo) was modeled. While some predictors submitted “best” models with relatively high TM scores (as high as 0.98), for GDT-TS the scores are generally lower (< 0.72). In nearly all cases, detailed analysis of these models demonstrates the significant challenges encountered by most groups in predicting key structural details, such as the coiled-coil handshake of targets M1228 and M1239, or key non-canonical base pairing interactions of target R1203.

Most of the successful methods used in these ensemble predictions leveraged AlphaFold2 and/or AlphaFold3 initial models and/or used these methods as the primary inference engines. In several cases, and particularly for the stem-loop RNA target R1203, AF3-server (304) provided top, or near-top, two-state ensemble predictions. AF3 was released just shortly before the CASP prediction season, and only a few predictors took advantage of AF3 models. Improvements relative to the baseline AF3-server were generally driven by improved methods for generating MSAs, comprehensive template selection, and final ranking of models.

Among the predictor groups, the most broadly successful with this ensemble target set were the Zhang group, which for example predicted accurate models of the T1214holo structure, key details of the coiled-coiled handshake of target M1228, and did well on several of the other ensemble targets. However, even these best-performing methods provided models with GDT-TS and DockQ scores lower than those obtained for typical CASP16 single-state targets (**Supplementary Figure S11**).

One significant challenge in predicting alternative conformational states is the tendency for ML-based modeling methods to be biased toward conformational states present in the training set, impacting their ability to model novel conformational states [31-41]. For example, *Cfold*, a structure prediction network trained on a conformational split of the PDB that excludes alternative states for protein structure pairs solved in two conformations [36], predicted > 50% of the known alternative conformations with high accuracy (TM-score > 0.75), but failed on the remaining pairs, unable to correctly model the alternative state not present in its training data. Lazou et al. observed successful predictions by AF2 of both open/closed states of cryptic binding sites in only six of 16 proteins tested [39]. In studies of the SLC-class of integral membrane proteins, we have also observed a strong preference for previously reported conformations that persist despite using several enhanced sampling methods [40]. While in some cases, the AI has learned enough to model alternative conformational states not included in the training data, in other cases memorization can be hard to overcome. However, while memorization can prevent modeling of alternative states using methods based on deep learning [34,36,38-40], at least for some of the predicted two-state ensembles of CASP16, methods were able to overcome this bias.

In the course of the study, several additional important concerns were raised. First is the distinction between conformational changes that are induced by ligands, binding partners, or sequence alterations, versus multiple conformational states that coexist within the same biomolecular sample, potentially reflecting dynamic conformational equilibria. The former generally require modeling both in the presence and absence of the modulating factor, while the latter can potentially be approached as alternative states of a single system, where binding shifts the equilibrium by conformational selection. Interestingly, in the case of the ligand-induced conformational change observed for target T1214, the *E. coli* TonB-dependent transporter PqqU bound to PQQ, some predictor groups were able to model the holo form without inclusion of the ligand. However, it may not always be possible to model the alternative conformational state without including the binding partner. Second is the challenge of assessing prediction models against experimental structures. The accuracy of any assessment is limited by the quality of the experimental model itself; in some cases it may be preferable to assess predictions directly against experimental data, such as cryoEM density maps or NMR NOESY peak lists. Third is the challenge of predicting relative conformer populations. Predictors may sometimes identify conformational states that are correct, biologically relevant, and energetically favorable, but absent or only lowly-populated under the experimental conditions used to determine the structure. Conversely, experimental models typically represent only a subset of the conformational ensemble, and conformations may be stabilized by conditions of data collection, or, in the case of X-ray crystallography, by the crystal lattice. These issues underscore the need for assessment strategies that account for conformational ensembles [5], rather than relying solely on individual experimental structures.

### Conclusions

CASP16 demonstrated the power and limitations of AlphaFold2/3 in modeling alternative conformational states, and the key conclusion that successful prediction of alternative conformations often depends on the availability of suitable templates and informative MSAs which were essential for modeling of fold rearrangements or domain repositioning in some targets. However, overall accuracy of these ensemble targets was lower than typically observed for single-state targets, and performance varied widely depending on target complexity. Multi-conformational RNA molecules proved especially challenging, with no cases successfully predicted. Importantly, relative state populations were generally not requested or attempted, and remain unassessed. These results highlight that, despite recent advances in AI-based structure prediction and the significant interest by the structural biology community in predicting multiple conformational states and conformational ensembles of proteins, nucleic acids, and their complexes, accurate modeling of multiple conformational states—particularly for RNA and large multimeric assemblies—remains a fundamental challenge and an important focus for future CASP experiments.

## Methods

### Structure comparison and alignment

Most structure quality statistics were computed by infrastructure developed and maintained by the CASP Prediction Center [42,43]. Predictor group identifiers were anonymized following CASP guidelines. Structural superpositions were performed using US-align [44], complemented by PyMOL [45] for visual inspection and analysis. Structural similarity scores backbone atom root mean squared deviation (RMSD), Template Modeling (TM), Global Distance Test (GDT_TS) [46] and LDDT [47] were used for global structure comparisons. Interface features were compared to reference structures using DockQ scores [27].

### Composite global-local scoring

To assess both the global fold accuracy and local conformational correctness of submitted models, we implemented a composite scoring strategy tailored to ensemble and multi-state targets. Scores from multiple global and local metrics were normalized to a 0 – 100 scale. For global accuracy, we applied several complementary measures. TM-score was used to evaluate overall topological similarity, while GDT_TS was used to capture both global and local structural agreement with the experimental structures. Global LDDT provided a superposition-free measure of distance preservation across all atom pairs. In addition, the global RMSD was included to capture the magnitude of atomic displacements, particularly useful for large rigid-body movements.

To complement global assessments, we also evaluated local structural accuracy using per-residue measures. These included "local" residue-level LDDT and a “local GDT_TS” approximation based on Cα–Cα deviations reported as LGA Dev values. LGA Dev, derived from the Local–Global Alignment algorithm, quantifies per-residue distance deviations from the reference and serves as a local analog to GDT_TS. To place both global and local measures on a comparable scale, we applied sigmoid-weighted scoring functions that convert raw RMSD and LGA Dev values into intuitive scores ranging from 0 to 100. These functions emphasize near-native conformations while smoothly down-weighting larger deviations (**Supplementary Figure S10**).

For RMSD, we used:

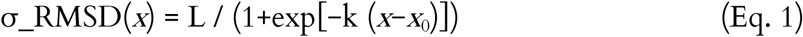

where *x* is the unweighted backbone RMSD (Å), Parameters were fit to yield scores of 100, 95, 80, 50, and 0 at 0, 1, 2, 4, and 8 Å, respectively, giving L = 105, k = −0.77, and *x*_0_ = 3.77.

For per-residue deviations, we applied the same functional form:

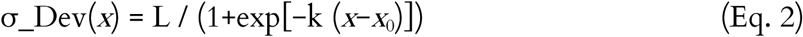

where *x* is the per-residue LGA Dev value (Å), and using identical parameters. Eq. 1 thus captures global RMSD differences, while Eq. 2 applies the same transformation to local deviations, allowing direct comparison of global and local conformational accuracy on a consistent 0–100 scale.

#### Composite global-local score definitions

To comprehensively evaluate both global and local structural accuracy across diverse targets, we defined four composite scoring schemes, each combining a subset of normalized structural metrics: GDT_TS, LDDT, σ_RMSD (Eqn 1), local LDDT, and σ_Dev (Eqn 2). Each metric was scaled to a 0–100 range. The definitions of composite scores are as follows:

- Σ1: All five metrics included, each contributing equally (weight = 1.0).
- Σ2: Same as Σ1, but excludes σ_RMSD to reduce the influence of global displacement in cases where local accuracy is of primary interest.
- Σ3: Applies a binary threshold (δ) to GDT_TS, LDDT, and σ_RMSD: scores ≥ 80 contribute 100; scores < 80 contribute 0. Local LDDT and σ_Dev are retained at full value (weight = 1.0).
- Σ4: Similar to Σ3, but thresholding (δ) is applied only to GDT_TS and LDDT; σ_RMSD, local LDDT, and σ_Dev remain fully weighted.

These variations allow scores to focus on both overall global accuracy with emphasis on local differences between the alternative states.

We initially evaluated all four composite scores (Σ1–Σ4) for target T1214. Among them, Σ4 provided the clearest separation of group performance and was therefore selected for ranking both T1214 and R1203. For other multi-domain targets (T1228, T1239, M1228, and M1239), we used global metrics only, including TM, GDT_TS, and LDDT. For target T1249, evaluation focused on the DockQ interface score. For RNA targets such as R1203, local measures (σ_Dev, lDDT) within loop regions (e.g., loop IIb, loop IIc) were emphasized to capture conformational rearrangements relevant to biological function (e.g., Rev binding and key non-canonical base pairing).

#### Two-state model evaluation

For ensemble targets with two experimentally determined conformational states, we evaluated the ability of predictor groups to accurately capture *both* states using a standardized Two-State Score (Σ*) procedure. Each group submitted up to ten models per target—typically five per state, but with unknown state assignments. All submitted models were compared to both reference conformations (designated as reference states v1 and v2) using structural metrics such as GDT_TS, LDDT, and composite scoring functions (see scoring section above). To avoid bias from the user-supplied state labels, we employed a model-matching protocol based on best structural alignment using selected scoring functions. As illustrated in **Figure 3**, the evaluation proceeded as follows:

1. **Score models against both reference structures** Let A = {a₁, a₂, …, a₅} and B = {b₁, b₂, …, b₅} be the two sets of predicted models. Each model *m* ∈ {A ∪ B} is scored against both reference structures v1 and v2 using a chosen structural metric, *S*(*m*, V):
  - *S_a_*_1_ = max{*S*(*m*, V1) for *m* ∈ A}
  - *S_a_*_2_ = max{*S*(*m*, V2) for *m* ∈ A}
  - *S_b_*_1_ = max{*S*(*m*, V1) for *m* ∈ B}
  - *S_b_*_2_ = max{*S*(*m*, V2) for *m* ∈ B}
  The highest of these four scores is identified.
2. **Assign sets to reference structures** Based on the highest-scoring model, the corresponding set is assigned to that reference structure.
  - If *S_a_*_1_ is the highest, assign set A to V1 and set B to V2.
  - If *S_a_*_2_ is the highest, assign set A to V2 and set B to V1.
  - If set A is assigned to V1, then B is assigned to V2 (or vice versa)
  - This assignment is fixed for scoring.
3. **Find best-matching models using the assigned reference state**
  - Within each set, identify the best-scoring model with respect to its assigned reference:
  - s_1_ = max{*S*(*m,* v1) for *m* ∈ assigned set for V1}
  - s_2_ = max{*S*(*m,* v2) for *m* ∈ assigned set for V2}
4. **Compute two-state score**
  - The two-state score is defined as S*= s_1_ + s_2_
  - For composite scores (e.g., Σ1–Σ4), s_1_ and s_2_ are the corresponding composite score for the best-scoring model to each reference state V1 and V2, as defined previously. The two state score is defined as Σ*=s_1_ + s_2_.
5. **Group Ranking** Predictor groups were then ranked based on their two-state scores, rewarding those that captured both alternate conformational states with high fidelity.

#### Molecular Graphics

Molecular visualization and preparation of graphical representations for figures was done using *PyMol* [45].

## Abbreviations

AF2: AlphaFold2 Multimer
cryoEM: cryogenic electron microscopy
GDT: Global Distance Test
LDDT: Local Difference Distance Test score, a superimposition independent metric of structure prediction accuracy
NMR: nuclear magnetic resonance spectroscopy
RDC: Residual Dipolar Coupling
RMSD: Root Mean Squared Deviation
SAXS: small angle X-ray scattering.

## Software and Data Availability

Data and scripts used in this work are available from our laboratory github site. https://github.rpi.edu/RPIBioinformatics/CASP16_TWO_STATE/

## Acknowledgements

This research was supported by grants R35-GM141818 (to GTM) and R01-GM100482 (to K. Fidelis, JM, and AK) from the National Institutes of Health, National Institute of General Medical Sciences.

## Author Contributions

ND, TAR, TLB, YJH, JM, AK, and GTM conceptualized the study. ND, TLB, TAR, JM, and AK carried out statistical analysis of protein and RNA structures. TLB and ND developed computer codes. All authors contributed in writing and editing the manuscript.

## Supplementary Tables

**Table S1.**
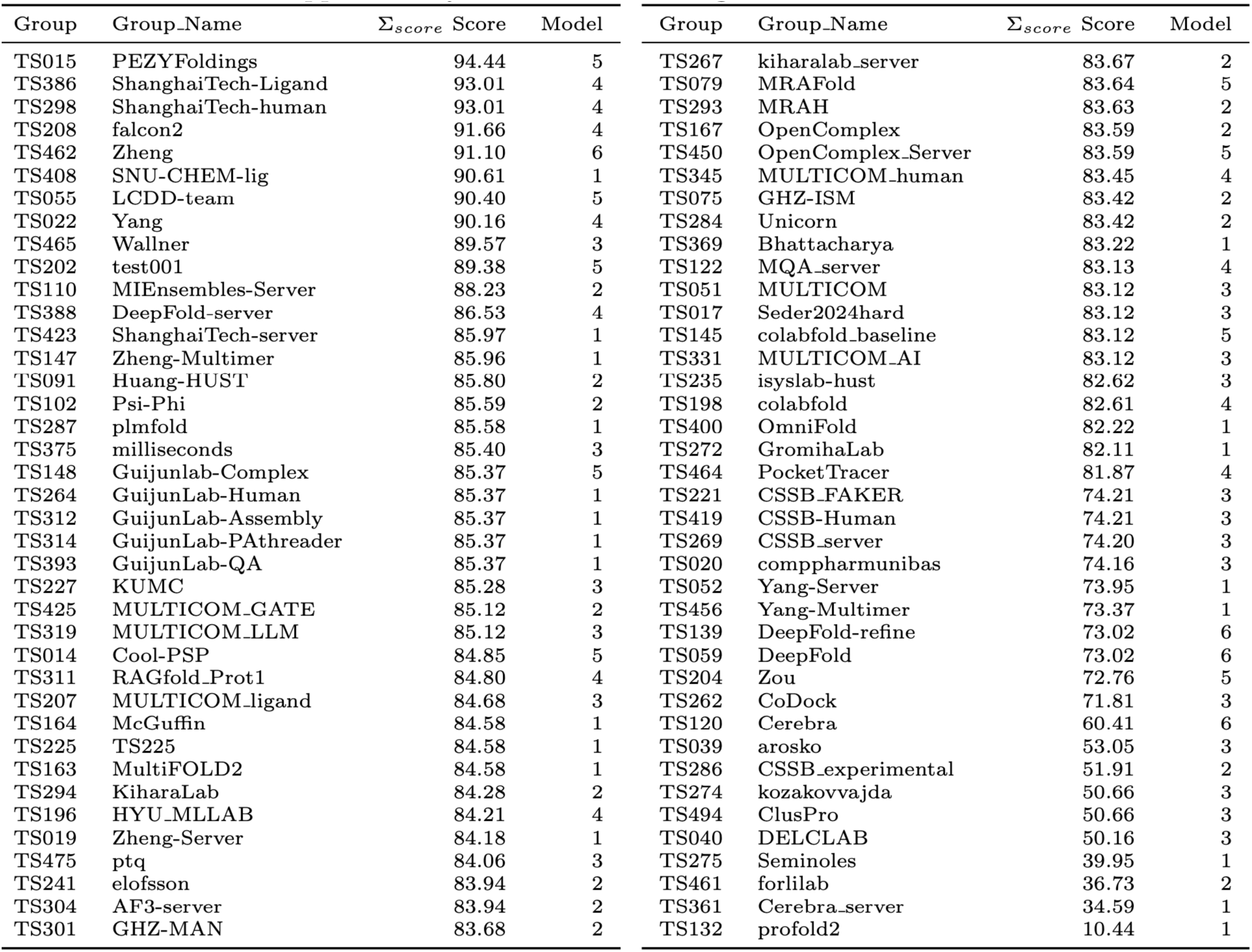
Ranking of all predictor groups for target T1214, based on best 𝜮𝟒 scoring model per group.

**Table S2.** Ranking of all predictor groups for target M1228, based on two-state TM-score.

| Group | Group_Name | Two-State_Score | V1_TMscore | V2_TMscore | V1_Model | V2_Model |
| --- | --- | --- | --- | --- | --- | --- |
| TS028 | NKRNA-s | 1.56 | 0.80 | 0.76 | TS028.v2.5 | TS028.v1.5 |
| TS462 | Zheng | 1.55 | 0.77 | 0.78 | TS462.v1.1 | TS462.v2.1 |
| TS481 | Vfold | 1.54 | 0.74 | 0.79 | TS481.v1.4 | TS481.v2.1 |
| TS286 | CSSB_experimental | 1.52 | 0.75 | 0.77 | TS286.v2.5 | TS286.v1.3 |
| TS091 | Huang-HUST | 1.50 | 0.71 | 0.79 | TS091.v2.1 | TS091.v1.2 |
| TS052 | Yang-Server | 1.50 | 0.69 | 0.81 | TS052.v2.3 | TS052.v1.1 |
| TS033 | Diff | 1.50 | 0.71 | 0.79 | TS033.v1.5 | TS033.v2.2 |
| TS022 | Yang | 1.50 | 0.70 | 0.80 | TS022.v1.4 | TS022.v2.2 |
| TS450 | OpenComplex_Server | 1.49 | 0.79 | 0.71 | TS450.v2.4 | TS450.v1.1 |
| TS167 | OpenComplex | 1.49 | 0.79 | 0.71 | TS167.v2.4 | TS167.v1.1 |
| TS304 | AF3-server | 1.49 | 0.71 | 0.78 | TS304.v1.4 | TS304.v2.5 |
| TS262 | CoDock | 1.49 | 0.80 | 0.70 | TS262.v1.1 | TS262.v2.3 |
| TS494 | ClusPro | 1.46 | 0.69 | 0.77 | TS494.v2.2 | TS494.v1.3 |
| TS274 | kozakovvajda | 1.46 | 0.69 | 0.77 | TS274.v2.2 | TS274.v1.3 |
| TS456 | Yang-Multimer | 1.46 | 0.67 | 0.78 | TS456.v2.3 | TS456.v1.1 |
| TS204 | Zou | 1.45 | 0.73 | 0.73 | TS204.v1.1 | TS204.v2.1 |
| TS241 | elofsson | 1.44 | 0.69 | 0.76 | TS241.v1.3 | TS241.v2.3 |
| TS110 | MIEnsembles-Server | 1.44 | 0.66 | 0.79 | TS110.v1.2 | TS110.v2.3 |
| TS294 | KiharaLab | 1.44 | 0.72 | 0.72 | TS294.v1.5 | TS294.v2.1 |
| TS369 | Bhattacharya | 1.40 | 0.72 | 0.69 | TS369.v2.5 | TS369.v1.2 |
| TS231 | B-LAB | 1.40 | 0.68 | 0.72 | TS231.v1.2 | TS231.v2.1 |
| TS208 | falcon2 | 1.37 | 0.70 | 0.67 | TS208.v2.5 | TS208.v1.1 |
| TS489 | Fernandez-Recio | 1.37 | 0.67 | 0.70 | TS489.v2.2 | TS489.v1.2 |
| TS323 | Yan | 1.33 | 0.65 | 0.68 | TS323.v1.1 | TS323.v2.1 |
| TS325 | 405 | 1.32 | 0.63 | 0.69 | TS325.v1.1 | TS325.v2.1 |
| TS159 | 406 | 1.32 | 0.63 | 0.69 | TS159.v1.1 | TS159.v2.1 |
| TS014 | Cool-PSP | 1.31 | 0.61 | 0.71 | TS014.v1.4 | TS014.v2.5 |

**Table S3.** Ranking of all predictor groups for target T1228, based on two-state GDT_TS scores.

| Group | Group_Name | Two-State_Score | V1_GDT_TS | V2_GDT_TS | V1_Model | V2_Model |
| --- | --- | --- | --- | --- | --- | --- |
| TS462 | Zheng | 116.15 | 63.11 | 53.04 | TS462.v1.3 | TS462.v2.5 |
| TS264 | GuijunLab-Human | 116.01 | 55.95 | 60.07 | TS264.v1.6 | TS264.v2.2 |
| TS147 | Zheng-Multimer | 115.68 | 62.31 | 53.37 | TS147.v1.1 | TS147.v2.1 |
| TS148 | Guijunlab-Complex | 114.51 | 54.45 | 60.07 | TS148.v2.3 | TS148.v1.3 |
| TS481 | Vfold | 112.45 | 61.80 | 50.66 | TS481.v1.3 | TS481.v2.5 |
| TS167 | OpenComplex | 112.41 | 57.07 | 55.34 | TS167.v1.1 | TS167.v2.3 |
| TS450 | OpenComplex_Server | 112.41 | 57.07 | 55.34 | TS450.v1.1 | TS450.v2.3 |
| TS022 | Yang | 112.13 | 57.21 | 54.92 | TS022.v1.4 | TS022.v2.5 |
| TS052 | Yang-Server | 111.89 | 59.50 | 52.39 | TS052.v1.4 | TS052.v2.2 |
| TS456 | Yang-Multimer | 110.72 | 58.33 | 52.39 | TS456.v1.3 | TS456.v2.2 |
| TS204 | Zou | 110.58 | 62.17 | 48.41 | TS204.v1.1 | TS204.v2.2 |
| TS241 | elofsson | 110.39 | 58.57 | 51.83 | TS241.v2.1 | TS241.v1.2 |
| TS294 | KiharaLab | 109.41 | 59.97 | 49.44 | TS294.v2.5 | TS294.v1.2 |
| TS091 | Huang-HUST | 109.18 | 54.54 | 54.63 | TS091.v1.2 | TS091.v2.5 |
| TS033 | Diff | 108.71 | 56.88 | 51.83 | TS033.v2.1 | TS033.v1.4 |
| TS262 | CoDock | 108.24 | 55.90 | 52.34 | TS262.v1.2 | TS262.v2.1 |
| TS212 | PIEFold_human | 107.58 | 55.10 | 52.48 | TS212.v2.4 | TS212.v1.2 |
| TS375 | milliseconds | 106.98 | 59.41 | 47.57 | TS375.v2.4 | TS375.v1.3 |
| TS028 | NKRNA-s | 106.69 | 51.31 | 55.38 | TS028.v1.2 | TS028.v2.5 |
| TS301 | GHZ-MAN | 106.65 | 55.95 | 50.70 | TS301.v1.3 | TS301.v2.4 |
| TS110 | MIEnsembles-Server | 105.81 | 52.25 | 53.56 | TS110.v1.2 | TS110.v2.5 |
| TS304 | AF3-server | 104.59 | 57.73 | 46.86 | TS304.v1.4 | TS304.v2.5 |
| TS331 | MULTICOM_AI | 103.98 | 56.65 | 47.33 | TS331.v1.2 | TS331.v2.4 |
| TS319 | MULTICOM_LLM | 103.98 | 56.65 | 47.33 | TS319.v1.2 | TS319.v2.4 |
| TS345 | MULTICOM_human | 103.98 | 56.65 | 47.33 | TS345.v1.3 | TS345.v2.2 |
| TS051 | MULTICOM | 103.98 | 56.65 | 47.33 | TS051.v1.3 | TS051.v2.2 |
| TS274 | kozakovvajda | 103.84 | 56.32 | 47.52 | TS274.v1.2 | TS274.v2.2 |
| TS494 | ClusPro | 103.84 | 56.32 | 47.52 | TS494.v1.2 | TS494.v2.2 |
| TS231 | B-LAB | 103.60 | 56.27 | 47.33 | TS231.v2.4 | TS231.v1.4 |
| TS314 | GuijunLab-Pathreader | 103.56 | 48.55 | 55.01 | TS314.v2.3 | TS314.v1.1 |
| TS286 | CSSB_experimental | 103.46 | 56.13 | 47.33 | TS286.v1.5 | TS286.v2.4 |
| TS425 | MULTICOM_GATE | 103.18 | 55.85 | 47.33 | TS425.v1.1 | TS425.v2.1 |
| TS198 | colabfold | 102.58 | 56.41 | 46.16 | TS198.v2.2 | TS198.v1.2 |
| TS419 | CSSB-Human | 102.29 | 55.52 | 46.77 | TS419.v2.4 | TS419.v1.3 |
| TS221 | CSSB_FAKER | 102.20 | 55.57 | 46.63 | TS221.v2.2 | TS221.v1.5 |
| TS369 | Bhattacharya | 102.11 | 54.82 | 47.28 | TS369.v1.2 | TS369.v2.1 |
| TS019 | Zheng-Server | 101.97 | 51.69 | 50.28 | TS019.v1.4 | TS019.v2.4 |
| TS475 | ptq | 101.12 | 58.43 | 42.70 | TS475.v2.5 | TS475.v1.2 |
| TS014 | Cool-PSP | 101.12 | 58.57 | 42.56 | TS014.v2.6 | TS014.v1.6 |
| TS287 | plmfold | 101.08 | 54.07 | 47.00 | TS287.v1.2 | TS287.v2.1 |
| TS267 | kiharalab_server | 100.05 | 54.68 | 45.37 | TS267.v1.1 | TS267.v2.5 |
| TS079 | MRAFold | 99.77 | 54.82 | 44.94 | TS079.v1.2 | TS079.v2.2 |
| TS293 | MRAH | 99.77 | 54.82 | 44.94 | TS293.v1.2 | TS293.v2.2 |
| TS489 | Fernandez-Recio | 99.67 | 55.48 | 44.20 | TS489.v1.2 | TS489.v2.2 |
| TS208 | falcon2 | 99.20 | 54.54 | 44.66 | TS208.v2.5 | TS208.v1.1 |
| TS196 | HYU_MLLAB | 98.74 | 53.37 | 45.37 | TS196.v1.5 | TS196.v2.5 |
| TS312 | GuijunLab-Assembly | 98.50 | 53.84 | 44.66 | TS312.v1.1 | TS312.v2.2 |
| TS112 | Seder2024easy | 97.99 | 54.35 | 43.63 | TS112.v1.4 | TS112.v2.3 |
| TS017 | Seder2024hard | 97.99 | 54.35 | 43.63 | TS017.v1.5 | TS017.v2.5 |
| TS145 | colabfold_baseline | 97.89 | 54.35 | 43.54 | TS145.v1.3 | TS145.v2.3 |
| TS465 | Wallner | 97.47 | 50.00 | 47.47 | TS465.v1.5 | TS465.v2.1 |
| TS235 | isyslab-hust | 96.63 | 50.80 | 45.83 | TS235.v1.1 | TS235.v2.3 |
| TS311 | RAGfold_Prot1 | 96.58 | 49.86 | 46.72 | TS311.v1.2 | TS311.v2.5 |
| TS163 | MultiFOLD2 | 96.02 | 53.23 | 42.79 | TS163.v1.5 | TS163.v2.2 |
| TS015 | PEZYFoldings | 95.55 | 51.12 | 44.43 | TS015.v2.2 | TS015.v1.5 |
| TS298 | ShanghaiTech-human | 95.04 | 54.35 | 40.68 | TS298.v1.1 | TS298.v2.3 |
| TS284 | Unicorn | 94.90 | 44.20 | 50.70 | TS284.v1.3 | TS284.v2.4 |
| TS075 | GHZ-ISM | 94.90 | 44.20 | 50.70 | TS075.v1.3 | TS075.v2.4 |
| TS122 | MQA_server | 94.90 | 44.20 | 50.70 | TS122.v1.3 | TS122.v2.4 |
| TS164 | McGuffin | 94.76 | 48.03 | 46.72 | TS164.v1.1 | TS164.v2.2 |
| TS325 | 405 | 94.29 | 49.34 | 44.94 | TS325.v1.1 | TS325.v2.1 |
| TS159 | 406 | 94.29 | 49.34 | 44.94 | TS159.v1.1 | TS159.v2.1 |
| TS269 | CSSB_server | 92.98 | 50.70 | 42.27 | TS269.v1.1 | TS269.v2.1 |
| TS059 | DeepFold | 91.53 | 46.91 | 44.62 | TS059.v2.3 | TS059.v1.3 |
| TS388 | DeepFold-server | 89.23 | 44.10 | 45.13 | TS388.v1.3 | TS388.v2.3 |
| TS023 | FTBiot0119 | 85.63 | 45.79 | 39.84 | TS023.v1.1 | TS023.v2.3 |
| TS139 | DeepFold-refine | 80.15 | 40.59 | 39.56 | TS139.v1.4 | TS139.v2.2 |
| TS120 | Cerebra | 76.08 | 38.34 | 37.73 | TS120.v1.4 | TS120.v2.1 |
| TS361 | Cerebra_server | 66.11 | 32.77 | 33.33 | TS361.v1.1 | TS361.v2.1 |

**Table S4.** Ranking of all predictor groups for target M1239, based on two-state TM-score.

| Group | Group_Name | Two-State_Score | V1_TMscore | V2_TMscore | V1_Model | V2_Model |
| --- | --- | --- | --- | --- | --- | --- |
| TS294 | KiharaLab | 1.45 | 0.81 | 0.64 | TS294_v1.1 | TS294_v2.3 |
| TS462 | Zheng | 1.41 | 0.75 | 0.66 | TS462_v2.3 | TS462_v1.1 |
| TS110 | MIEnsembles-Server | 1.39 | 0.73 | 0.66 | TS110_v2.5 | TS110_v1.1 |
| TS262 | CoDock | 1.37 | 0.74 | 0.63 | TS262_v1.3 | TS262_v2.4 |
| TS028 | NKRNA-s | 1.37 | 0.72 | 0.64 | TS028_v1.5 | TS028_v2.1 |
| TS235 | isyslab-hust | 1.35 | 0.67 | 0.68 | TS235_v2.4 | TS235_v1.5 |
| TS489 | Fernandez-Recio | 1.33 | 0.71 | 0.63 | TS489_v2.2 | TS489_v1.3 |
| TS481 | Vfold | 1.32 | 0.67 | 0.65 | TS481_v1.5 | TS481_v2.5 |
| TS204 | Zou | 1.29 | 0.64 | 0.65 | TS204_v2.1 | TS204_v1.5 |
| TS208 | falcon2 | 1.27 | 0.67 | 0.60 | TS208_v1.4 | TS208_v2.1 |
| TS325 | 405 | 1.27 | 0.64 | 0.63 | TS325_v1.2 | TS325_v2.2 |
| TS159 | 406 | 1.27 | 0.64 | 0.63 | TS159_v1.2 | TS159_v2.2 |
| TS286 | CSSB_experimental | 1.27 | 0.65 | 0.62 | TS286_v1.2 | TS286_v2.2 |
| TS033 | Diff | 1.25 | 0.65 | 0.61 | TS033_v2.4 | TS033_v1.2 |
| TS231 | B-LAB | 1.25 | 0.64 | 0.60 | TS231_v1.2 | TS231_v2.1 |
| TS091 | Huang-HUST | 1.23 | 0.64 | 0.59 | TS091_v2.2 | TS091_v1.5 |
| TS369 | Bhattacharya | 1.22 | 0.61 | 0.61 | TS369_v2.3 | TS369_v1.3 |
| TS167 | OpenComplex | 1.19 | 0.57 | 0.62 | TS167_v2.1 | TS167_v1.1 |
| TS450 | OpenComplex_Server | 1.19 | 0.57 | 0.62 | TS450_v2.1 | TS450_v1.1 |
| TS014 | Cool-PSP | 1.06 | 0.52 | 0.54 | TS014_v1.5 | TS014_v2.4 |
| TS272 | GromihaLab | 0.77 | 0.36 | 0.41 | TS272_v2.1 | TS272_v1.3 |
| TS241 | elofsson | 0.69 | 0.35 | 0.34 | TS241_v1.4 | TS241_v2.1 |
| TS304 | AF3-server | 0.66 | 0.33 | 0.33 | TS304_v2.4 | TS304_v1.1 |

**Table S5.** Ranking of all predictor groups for target T1239, based on two-state GDT-TS score.

| Group | Group_Name | Two-State_Score | V1_GDT-TS | V2_GDT-TS | V1_Model | V2_Model |
| --- | --- | --- | --- | --- | --- | --- |
| TS462 | Zheng | 119.79 | 70.81 | 48.98 | TS462.v2.2 | TS462.v1.1 |
| TS235 | isyslab-hust | 114.17 | 71.31 | 42.85 | TS235.v2.4 | TS235.v1.1 |
| TS419 | CSSB-Human | 114.04 | 71.10 | 42.94 | TS419.v2.1 | TS419.v1.5 |
| TS221 | CSSB-FAKER | 114.04 | 71.10 | 42.94 | TS221.v2.1 | TS221.v1.5 |
| TS028 | NKRNA-s | 113.49 | 71.36 | 42.13 | TS028.v2.2 | TS028.v1.2 |
| TS345 | MULTICOM_human | 113.32 | 71.23 | 42.09 | TS345.v1.4 | TS345.v2.5 |
| TS051 | MULTICOM | 113.32 | 71.23 | 42.09 | TS051.v1.4 | TS051.v2.5 |
| TS286 | CSSB_experimental | 112.64 | 69.87 | 42.77 | TS286.v2.1 | TS286.v1.2 |
| TS208 | falcon2 | 108.53 | 66.99 | 41.54 | TS208.v1.4 | TS208.v2.4 |
| TS425 | MULTICOM_GATE | 107.68 | 63.22 | 44.46 | TS425.v1.2 | TS425.v2.4 |
| TS019 | Zheng-Server | 106.22 | 54.02 | 52.20 | TS019.v1.5 | TS019.v2.5 |
| TS022 | Yang | 105.14 | 60.72 | 44.42 | TS022.v1.5 | TS022.v2.4 |
| TS204 | Zou | 104.76 | 61.23 | 43.53 | TS204.v1.5 | TS204.v2.2 |
| TS110 | MIEnsembles-Server | 104.72 | 63.01 | 41.71 | TS110.v1.1 | TS110.v2.2 |
| TS456 | Yang-Multimer | 104.50 | 60.21 | 44.29 | TS456.v1.3 | TS456.v2.2 |
| TS147 | Zheng-Multimer | 103.35 | 56.27 | 47.08 | TS147.v1.3 | TS147.v2.5 |
| TS167 | OpenComplex | 102.59 | 54.07 | 48.52 | TS167.v1.5 | TS167.v2.3 |
| TS450 | OpenComplex_Server | 102.59 | 54.07 | 48.52 | TS450.v1.5 | TS450.v2.3 |
| TS294 | KiharaLab | 101.71 | 59.58 | 42.13 | TS294.v1.1 | TS294.v2.2 |
| TS314 | GuijunLab-Pathreader | 101.54 | 58.77 | 42.77 | TS314.v2.5 | TS314.v1.2 |
| TS052 | Yang-Server | 100.05 | 55.93 | 44.12 | TS052.v1.5 | TS052.v2.2 |
| TS481 | Vfold | 99.67 | 56.57 | 43.10 | TS481.v2.5 | TS481.v1.2 |
| TS091 | Huang-HUST | 99.24 | 52.08 | 47.17 | TS091.v1.5 | TS091.v2.3 |
| TS319 | MULTICOM_LLM | 98.53 | 55.64 | 42.89 | TS319.v1.2 | TS319.v2.5 |
| TS262 | CoDock | 98.36 | 57.63 | 40.74 | TS262.v1.2 | TS262.v2.1 |
| TS033 | Diff | 97.93 | 53.94 | 43.99 | TS033.v2.2 | TS033.v1.1 |
| TS331 | MULTICOM_AI | 97.73 | 55.64 | 42.09 | TS331.v1.2 | TS331.v2.2 |
| TS325 | 405 | 97.47 | 55.21 | 42.26 | TS325.v1.2 | TS325.v2.2 |
| TS159 | 406 | 97.47 | 55.21 | 42.26 | TS159.v1.2 | TS159.v2.2 |
| TS369 | Bhattacharya | 97.17 | 54.07 | 43.10 | TS369.v1.4 | TS369.v2.4 |
| TS489 | Fernandez-Recio | 96.71 | 54.96 | 41.75 | TS489.v1.3 | TS489.v2.4 |
| TS298 | ShanghaiTech-human | 96.00 | 61.74 | 34.26 | TS298.v1.1 | TS298.v2.1 |
| TS212 | PIEFold_human | 95.91 | 55.21 | 40.69 | TS212.v2.1 | TS212.v1.3 |
| TS241 | elofsson | 95.14 | 54.32 | 40.82 | TS241.v1.1 | TS241.v2.5 |
| TS231 | B-LAB | 94.34 | 54.07 | 40.27 | TS231.v2.1 | TS231.v1.2 |
| TS304 | AF3-server | 94.04 | 50.89 | 43.15 | TS304.v2.5 | TS304.v1.2 |
| TS465 | Wallner | 92.97 | 48.14 | 44.84 | TS465.v1.4 | TS465.v2.2 |
| TS358 | PerezLab_Gators | 92.78 | 54.96 | 37.82 | TS358.v2.2 | TS358.v1.4 |
| TS293 | MRAH | 91.92 | 49.91 | 42.01 | TS293.v1.2 | TS293.v2.3 |
| TS079 | MRAFold | 91.92 | 49.91 | 42.01 | TS079.v1.2 | TS079.v2.3 |
| TS272 | GromihaLab | 91.33 | 50.09 | 41.24 | TS272.v1.1 | TS272.v2.1 |
| TS423 | ShanghaiTech-server | 91.08 | 51.91 | 39.17 | TS423.v2.4 | TS423.v1.1 |
| TS267 | kiharalab_server | 89.30 | 50.00 | 39.30 | TS267.v1.1 | TS267.v2.4 |
| TS148 | Guijunlab-Complex | 88.53 | 48.94 | 39.59 | TS148.v2.5 | TS148.v1.4 |
| TS287 | plmfold | 88.49 | 49.32 | 39.17 | TS287.v1.3 | TS287.v2.4 |
| TS198 | colabfold | 88.11 | 50.34 | 37.77 | TS198.v1.2 | TS198.v2.1 |
| TS375 | milliseconds | 87.73 | 47.16 | 40.57 | TS375.v2.3 | TS375.v1.4 |
| TS112 | Seder2024easy | 86.88 | 48.09 | 38.79 | TS112.v1.4 | TS112.v2.5 |
| TS014 | Cool-PSP | 86.84 | 49.49 | 37.35 | TS014.v2.3 | TS014.v1.3 |
| TS301 | GHZ-MAN | 86.84 | 46.65 | 40.19 | TS301.v2.4 | TS301.v1.1 |
| TS163 | MultiFOLD2 | 86.72 | 48.39 | 38.33 | TS163.v1.3 | TS163.v2.3 |
| TS040 | DELCLAB | 86.50 | 48.09 | 38.41 | TS040.v1.3 | TS040.v2.3 |
| TS145 | colabfold_baseline | 86.46 | 47.67 | 38.79 | TS145.v1.2 | TS145.v2.1 |
| TS023 | FTBiot0119 | 86.46 | 47.67 | 38.79 | TS023.v1.2 | TS023.v2.1 |
| TS264 | GuijunLab-Human | 86.20 | 47.54 | 38.66 | TS264.v2.2 | TS264.v1.5 |
| TS312 | GuijunLab-Assembly | 85.86 | 46.27 | 39.59 | TS312.v2.4 | TS312.v1.4 |
| TS122 | MQA_server | 85.78 | 46.65 | 39.13 | TS122.v1.3 | TS122.v2.2 |
| TS139 | DeepFold-refine | 84.76 | 45.93 | 38.83 | TS139.v1.6 | TS139.v2.6 |
| TS059 | DeepFold | 84.76 | 45.93 | 38.83 | TS059.v1.6 | TS059.v2.6 |
| TS017 | Seder2024hard | 84.52 | 48.05 | 36.46 | TS017.v1.5 | TS017.v2.4 |
| TS397 | smg_ulaval | 84.13 | 45.00 | 39.13 | TS397.v1.1 | TS397.v2.1 |
| TS164 | McGuffin | 83.83 | 45.47 | 38.37 | TS164.v2.2 | TS164.v1.3 |
| TS015 | PEZYFoldings | 83.79 | 45.00 | 38.79 | TS015.v1.4 | TS015.v2.1 |
| TS196 | HYU_MLLAB | 78.41 | 38.94 | 39.47 | TS196.v1.2 | TS196.v2.2 |
| TS388 | DeepFold-server | 75.91 | 37.92 | 37.99 | TS388.v1.3 | TS388.v2.2 |
| TS120 | Cerebra | 61.43 | 31.48 | 29.95 | TS120.v1.3 | TS120.v2.2 |
| TS475 | ptq | 49.20 | 49.20 | 0.00 | TS475.v2.1 | N/A<sup>1</sup> |
| TS351 | digiwiser-ensemble | 48.73 | 24.45 | 24.28 | TS351.v1.1 | TS351.v2.1 |
| TS269 | CSSB_server | 48.18 | 48.18 | 0.00 | TS269.v2.4 | N/A<sup>1</sup> |
| TS075 | GHZ-ISM | 46.65 | 46.65 | 0.00 | TS075.v2.4 | N/A<sup>1</sup> |
| TS284 | Unicorn | 46.65 | 46.65 | 0.00 | TS284.v2.4 | N/A<sup>1</sup> |
| TS311 | RAGfold_Prot1 | 42.25 | 42.25 | 0.00 | TS311.v1.1 | N/A<sup>1</sup> |
| TS361 | Cerebra_server | 31.40 | 31.40 | 0.00 | TS361.v1.2 | N/A<sup>1</sup> |
| TS105 | PFSC-PFVM | 4.20 | 4.20 | 0.00 | TS105.v1.3 | N/A<sup>1</sup> |

<sup>1</sup> Model either not submitted or not assessed

**Table S6.**
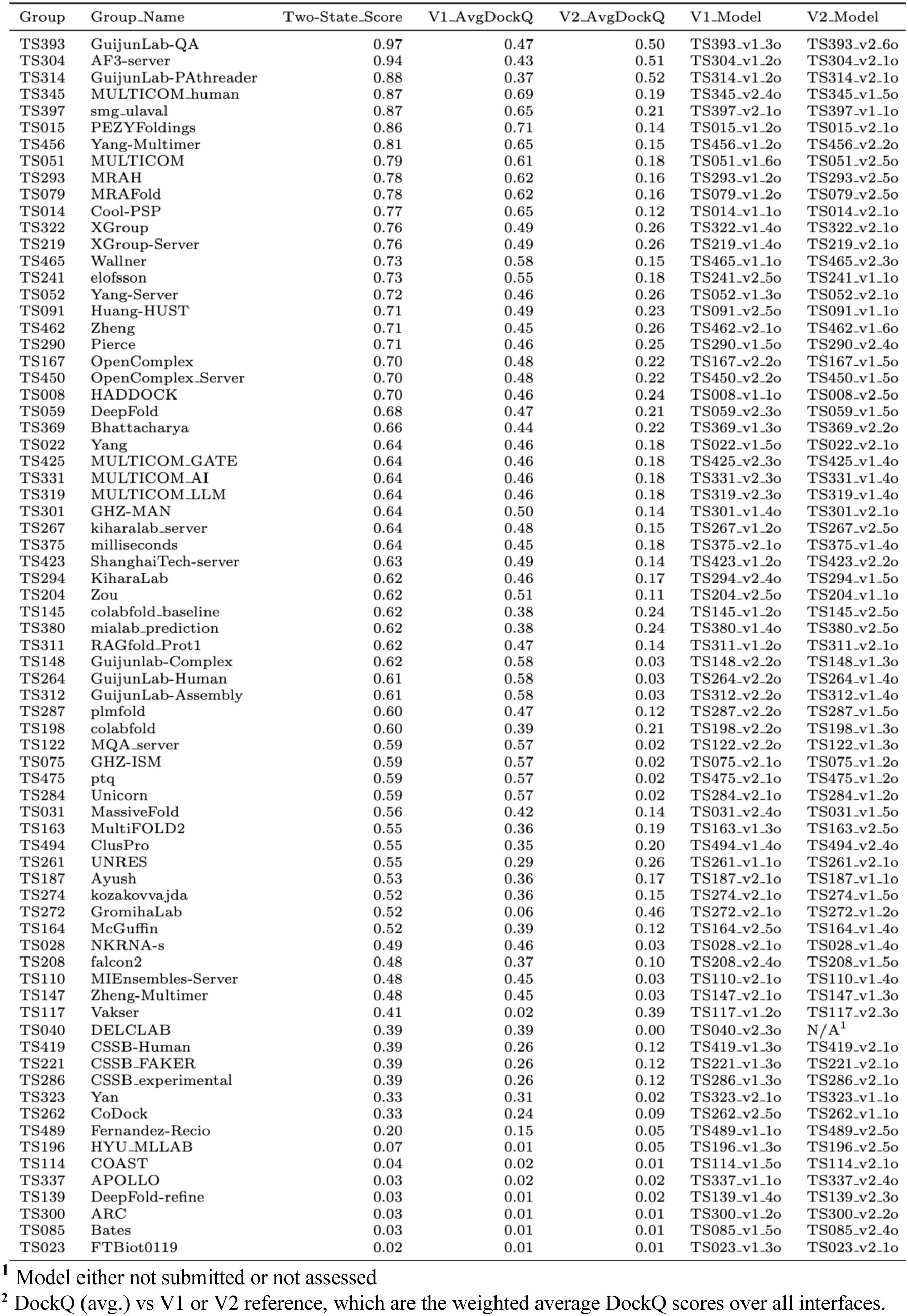
Ranking of all predictor groups for target T1249, based on two-state <DOCKQ> score.

**Table S7.** Ranking of all predictor groups for target R1203, based on two-state 𝜮4 global/local composite scores.

| Group | Group_Name | Two-State_Score | V1-Composite_Score_4 | V2-Composite_Score_4 | V1_Model | V2_Model |
| --- | --- | --- | --- | --- | --- | --- |
| TS304 | AF3-server | 85.08 | 50.07 | 35.01 | TS304.v1.3 | TS304.v2.2 |
| TS241 | elofsson | 85.08 | 50.07 | 35.01 | TS241.v1.3 | TS241.v2.2 |
| TS235 | isyslab-hust | 83.60 | 46.42 | 37.18 | TS235.v1.2 | TS235.v2.5 |
| TS435 | RNAFOLDX | 77.05 | 47.74 | 29.31 | TS435.v1.5 | TS435.v2.6 |
| TS369 | Bhattacharya | 73.39 | 40.37 | 33.02 | TS369.v1.2 | TS369.v2.5 |
| TS006 | RNA_Dojo | 70.31 | 30.80 | 39.51 | TS006.v1.1 | TS006.v2.3 |
| TS272 | GromihaLab | 67.08 | 31.19 | 35.89 | TS272.v1.4 | TS272.v2.1 |
| TS231 | B-LAB | 66.58 | 28.13 | 38.45 | TS231.v1.1 | TS231.v2.5 |
| TS159 | 406 | 59.61 | 31.73 | 27.89 | TS159.v1.2 | TS159.v2.1 |
| TS238 | BRIQX | 49.74 | 11.68 | 38.06 | TS238.v1.2 | TS238.v2.5 |
| TS294 | KiharaLab | 42.72 | 10.72 | 31.99 | TS294.v1.4 | TS294.v2.1 |
| TS267 | kiharalab_server | 42.25 | 19.02 | 23.23 | TS267.v1.1 | TS267.v2.3 |
| TS167 | OpenComplex | 42.16 | 9.21 | 32.95 | TS167.v1.5 | TS167.v2.2 |
| TS156 | SoutheRNA | 40.91 | 25.12 | 15.79 | TS156.v1.1 | TS156.v2.4 |
| TS052 | Yang-Server | 39.88 | 19.93 | 19.95 | TS052.v1.2 | TS052.v2.4 |
| TS028 | NKRNA-s | 36.51 | 24.46 | 12.05 | TS028.v1.3 | TS028.v2.2 |
| TS165 | dfr | 35.44 | 11.21 | 24.23 | TS165.v1.3 | TS165.v2.5 |
| TS400 | OmniFold | 35.24 | 0.00 | 35.24 | N/A<sup>1</sup> | TS400.v2.1 |
| TS110 | MIEnsembles-Server | 35.06 | 7.94 | 27.12 | TS110.v1.2 | TS110.v2.5 |
| TS189 | LCBio | 34.89 | 23.45 | 11.44 | TS189.v1.5 | TS189.v2.1 |
| TS417 | GuangzhouRNA-meta | 34.68 | 9.97 | 24.71 | TS417.v1.5 | TS417.v2.4 |
| TS286 | CSSB_experimental | 34.08 | 18.77 | 15.31 | TS286.v1.1 | TS286.v2.3 |
| TS276 | FrederickFolding | 33.62 | 0.00 | 33.62 | N/A<sup>1</sup> | TS276.v2.1 |
| TS317 | GuangzhouRNA_AI | 33.16 | 9.19 | 23.97 | TS317.v1.5 | TS317.v2.4 |
| TS448 | dNAfold | 30.11 | 20.49 | 9.62 | TS448.v1.1 | TS448.v2.5 |
| TS169 | thermomaps | 28.57 | 12.37 | 16.20 | TS169.v1.5 | TS169.v2.2 |
| TS462 | Zheng | 27.64 | 7.94 | 19.70 | TS462.v1.2 | TS462.v2.1 |
| TS183 | GuangzhouRNA-human | 26.84 | 16.07 | 10.77 | TS183.v1.5 | TS183.v2.2 |
| TS481 | Vfold | 26.72 | 9.62 | 17.10 | TS481.v1.4 | TS481.v2.5 |
| TS358 | PerezLab_Gators | 25.81 | 11.55 | 14.27 | TS358.v1.2 | TS358.v2.1 |
| TS261 | UNRES | 25.29 | 9.56 | 15.73 | TS261.v1.1 | TS261.v2.3 |
| TS338 | GeneSilico | 25.14 | 10.67 | 14.47 | TS338.v1.5 | TS338.v2.3 |
| TS456 | Yang-Multimer | 23.94 | 13.31 | 10.63 | TS456.v1.2 | TS456.v2.4 |
| TS094 | SimRNA-Server | 22.53 | 13.25 | 9.28 | TS094.v1.2 | TS094.v2.3 |
| TS063 | RNApolis | 22.33 | 10.21 | 12.13 | TS063.v1.4 | TS063.v2.3 |
| TS450 | OpenComplex_Server | 18.83 | 9.97 | 8.86 | TS450.v1.2 | TS450.v2.4 |
| TS298 | ShanghaiTech-human | 9.18 | 9.18 | 0.00 | TS298.v1.1 | N/A<sup>1</sup> |
| TS423 | ShanghaiTech-server | 9.18 | 9.18 | 0.00 | TS423.v1.1 | N/A<sup>1</sup> |
| TS367 | AIR | 9.14 | 0.00 | 9.14 | N/A<sup>1</sup> | TS367.v2.1 |
| TS306 | GeneSilicoRNA-server | 6.35 | 0.00 | 6.35 | N/A<sup>1</sup> | TS306.v2.1 |
| TS208 | falcon2 | 4.34 | 4.34 | 0.00 | TS208.v1.1 | N/A<sup>1</sup> |
| TS403 | mmagnus | 2.86 | 0.00 | 2.86 | N/A<sup>1</sup> | TS403.v2.1 |

<sup>1</sup> Model either not submitted or not assessed

**Table S8.**
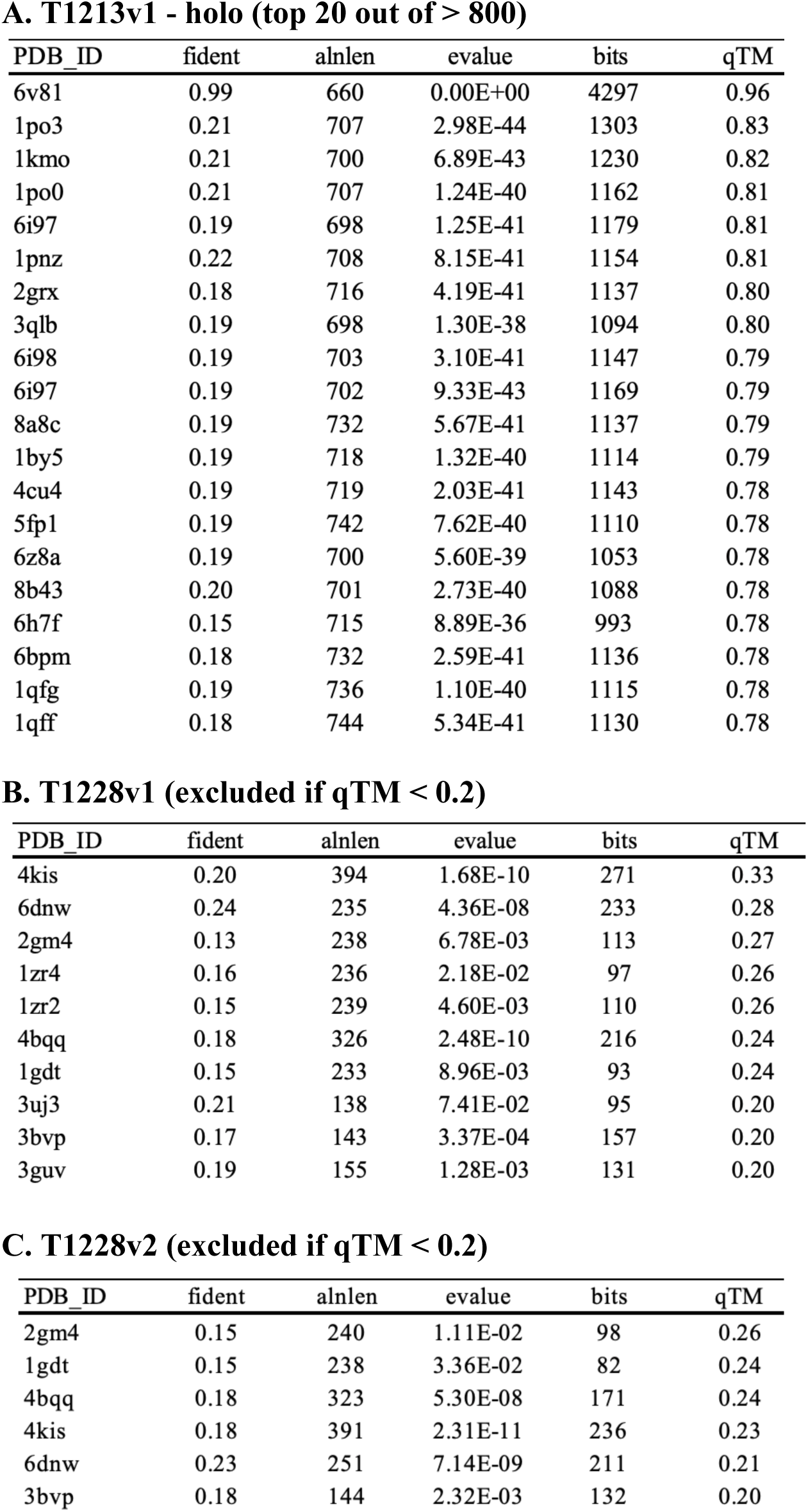
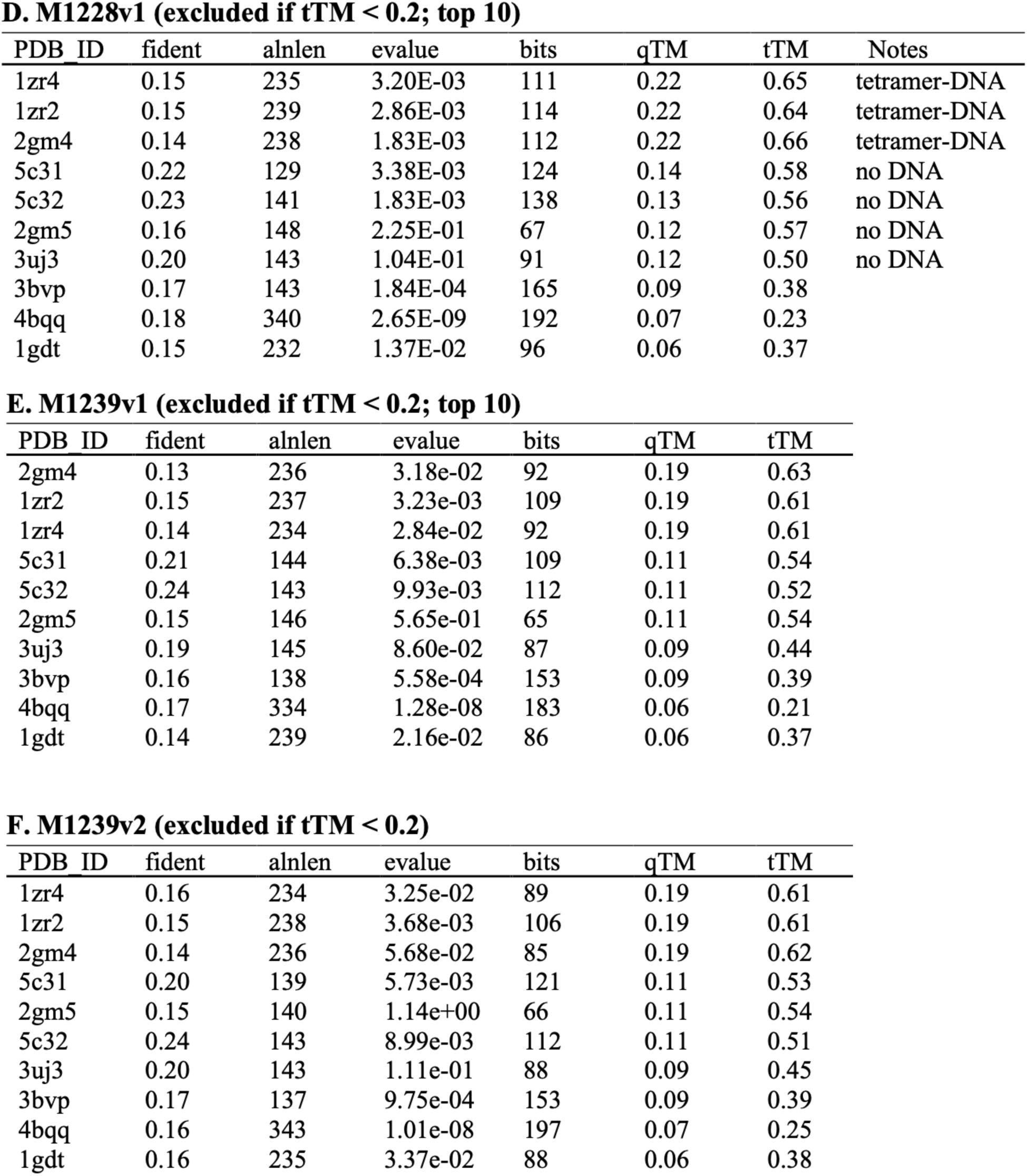
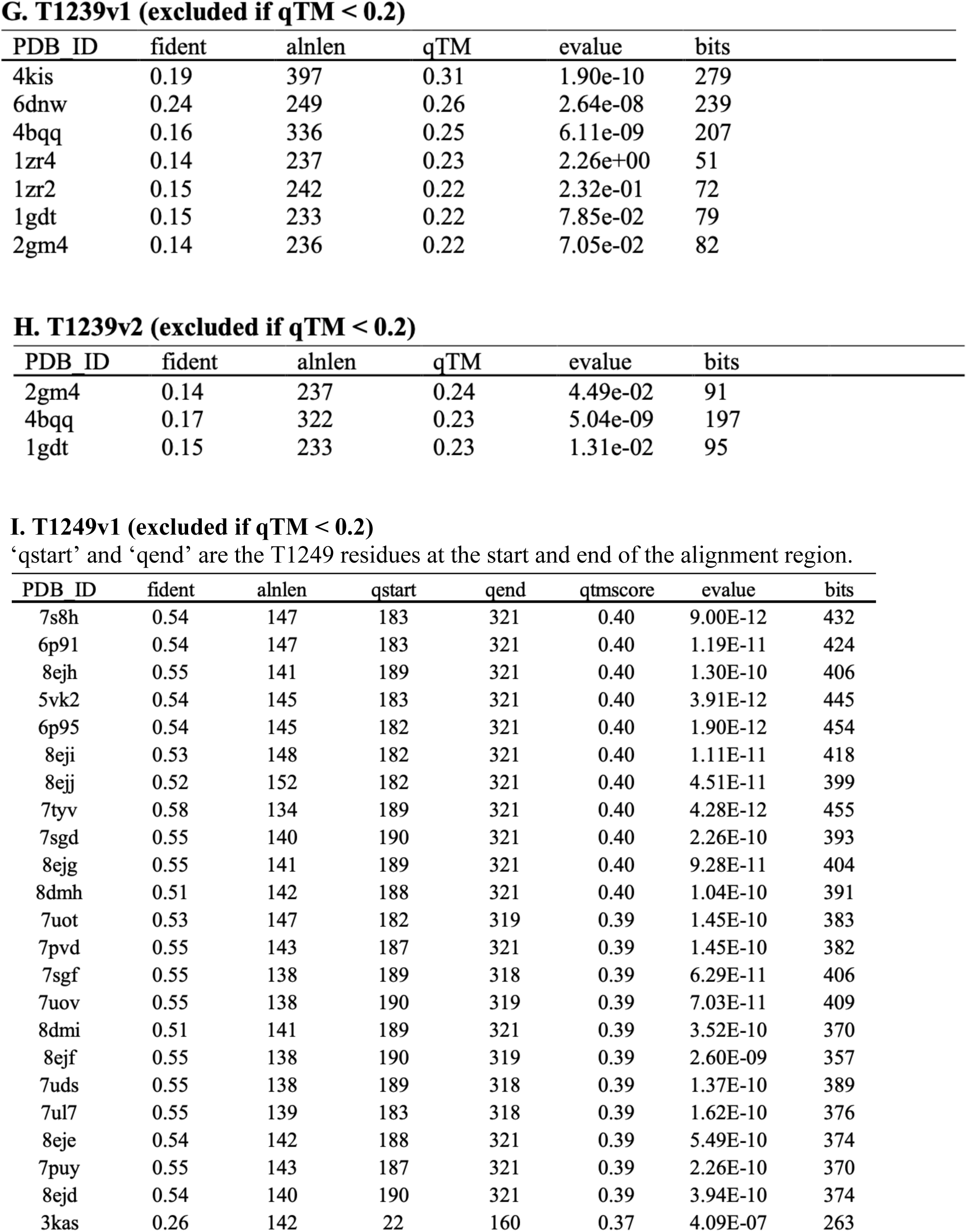
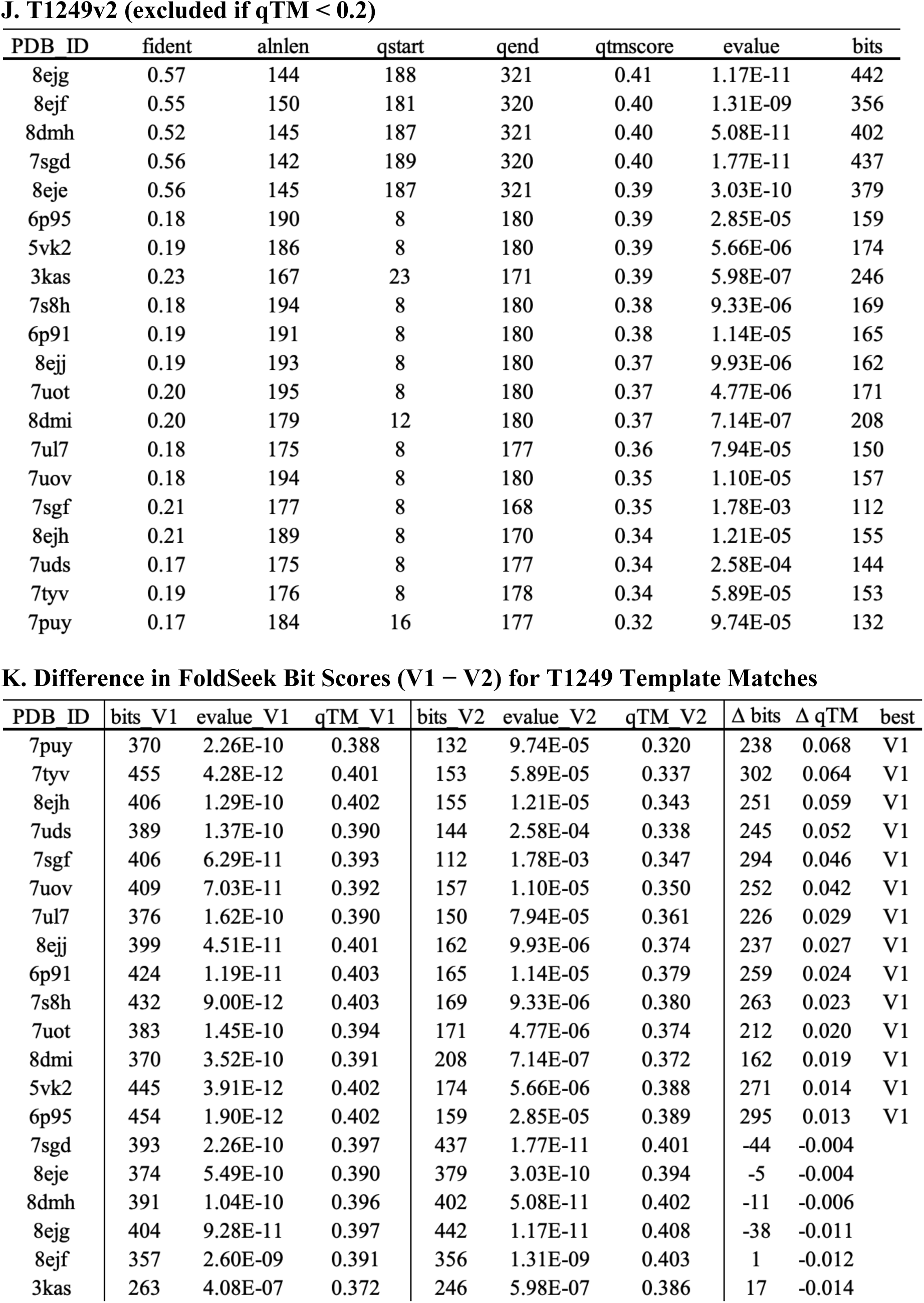
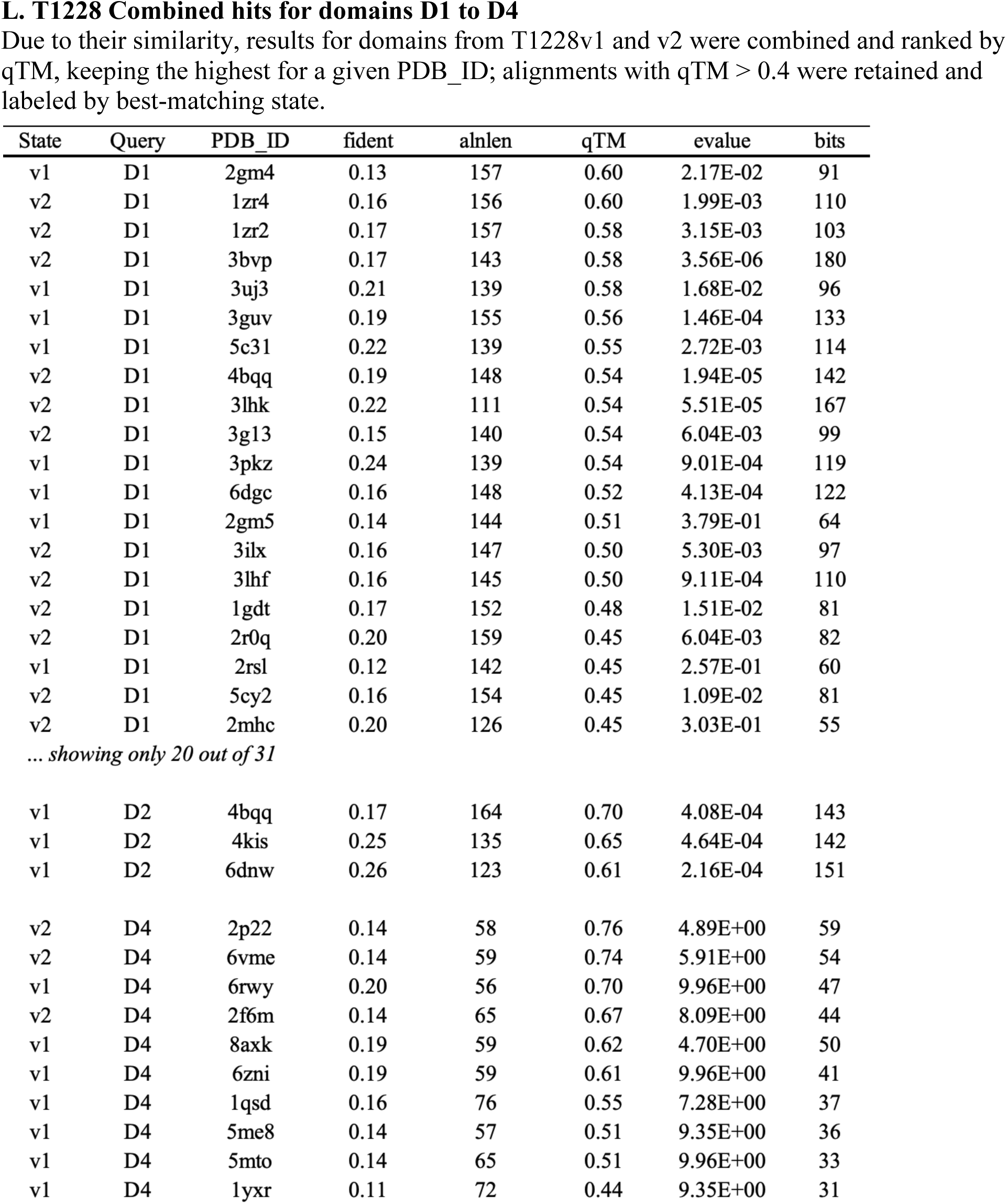
FoldSeek output summary tables by target. Homologs ranked by qTM score, with tTM (if available) or bitscore as the tie-breakers. Only the best hit per PDB ID was retained. ‘fident’ is the fraction of identical matched residues after structural alignment with the given alignment length, ‘alnlen’.

## Supplementary Figures

**Fig. S1.**
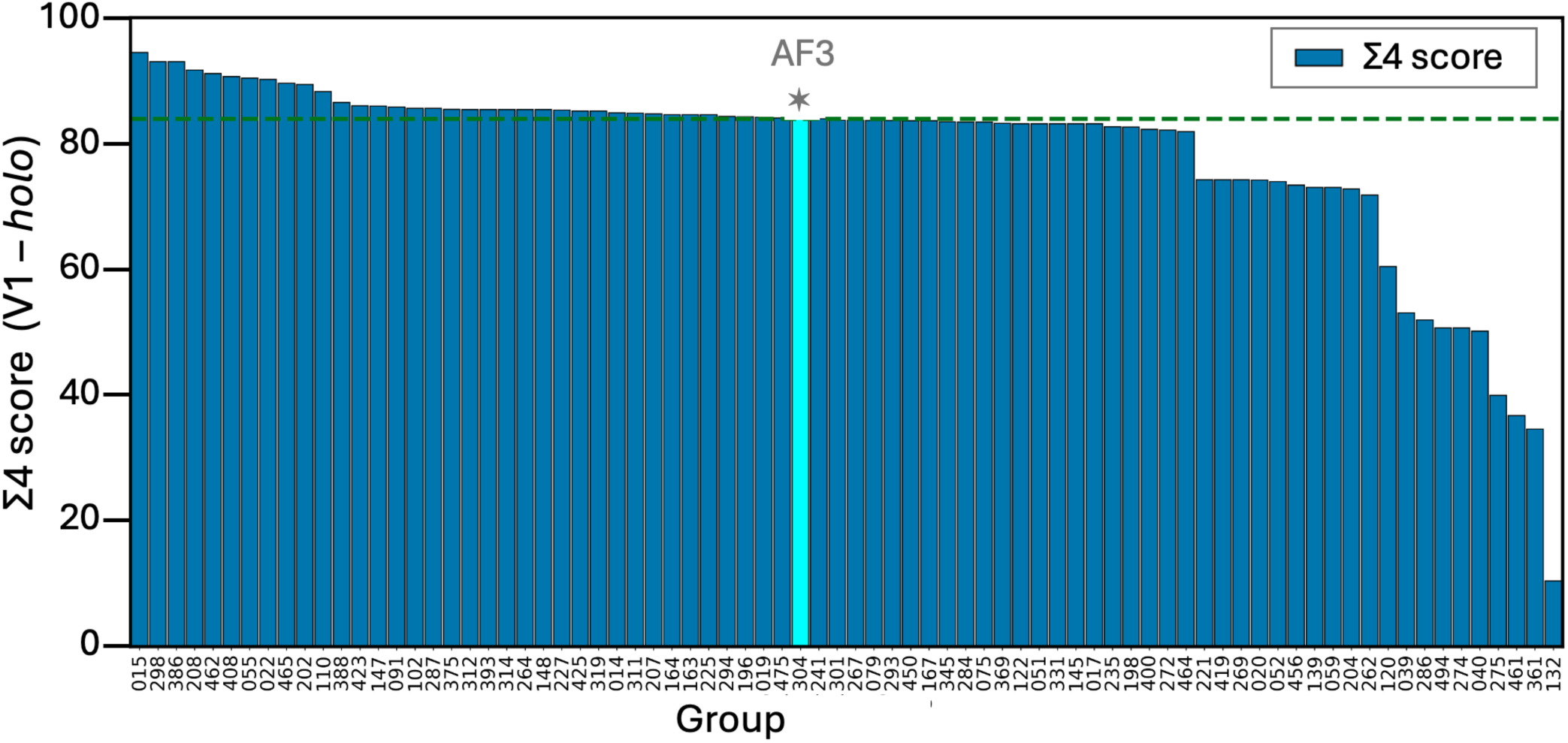
Expanded version of panels 1C for target T1214. Composite Σ4 scores for T1214 holo (V1). Values for best AF3-server (304) models are indicated by * and vertical lt. blue histogram column.

**Fig. S2.**
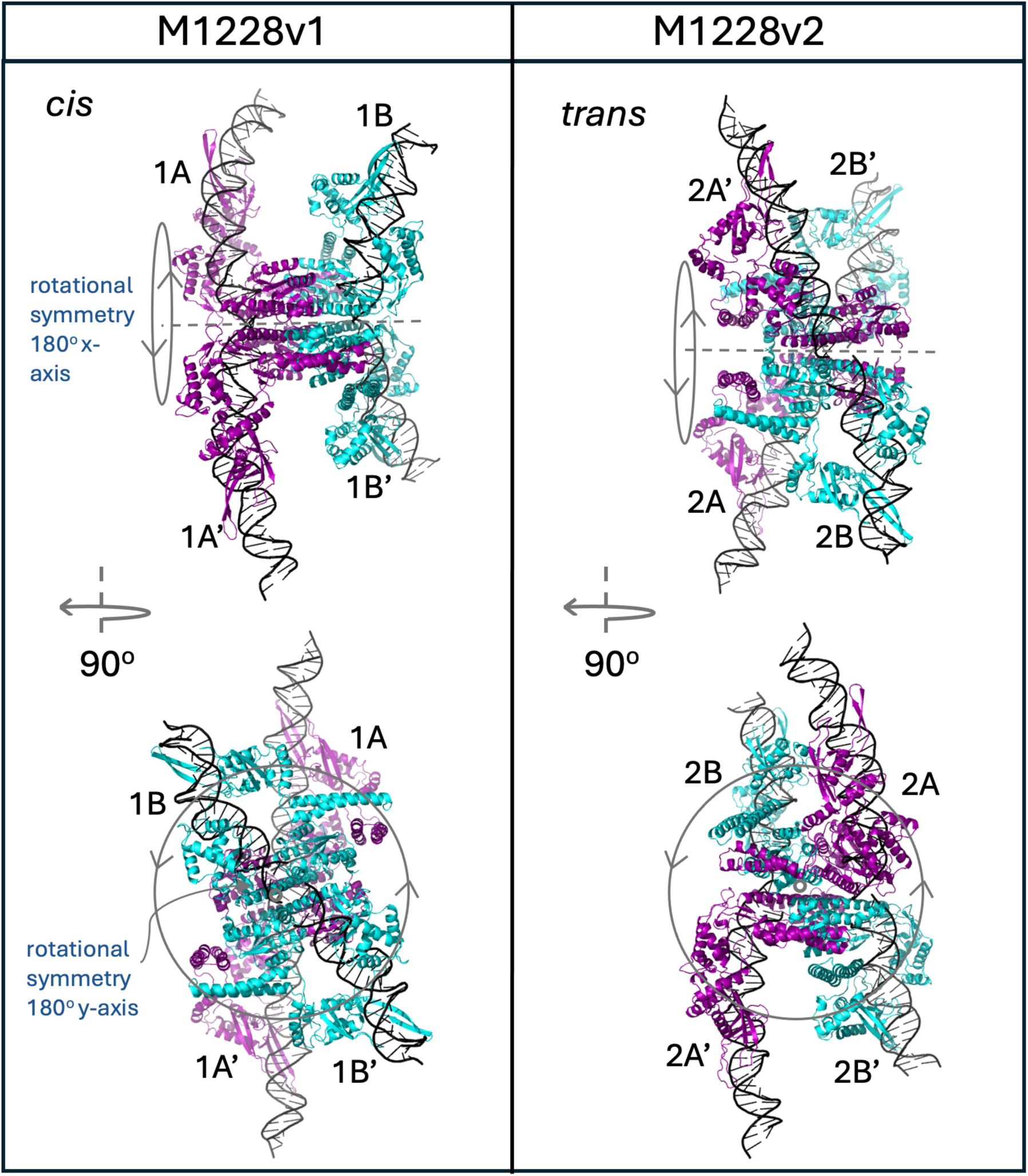
Symmetry in target M1228. Reference structures of M1228 in two conformational states (V1 and V2), each forming tetrameric assemblies with four DNA fragments. Chains are colored by conformation (“V-shaped” in magenta and “compact-shaped” in cyan), and subunits are labeled to highlight interdomain rearrangements between conformational states. A grey dashed line marks the axis of symmetry of the complex, which is different for V1 vs V2. Lower structures show the symmetry of each conformation as seen from after a 90° z-axis rotation (side-view).

**Fig. S3.**
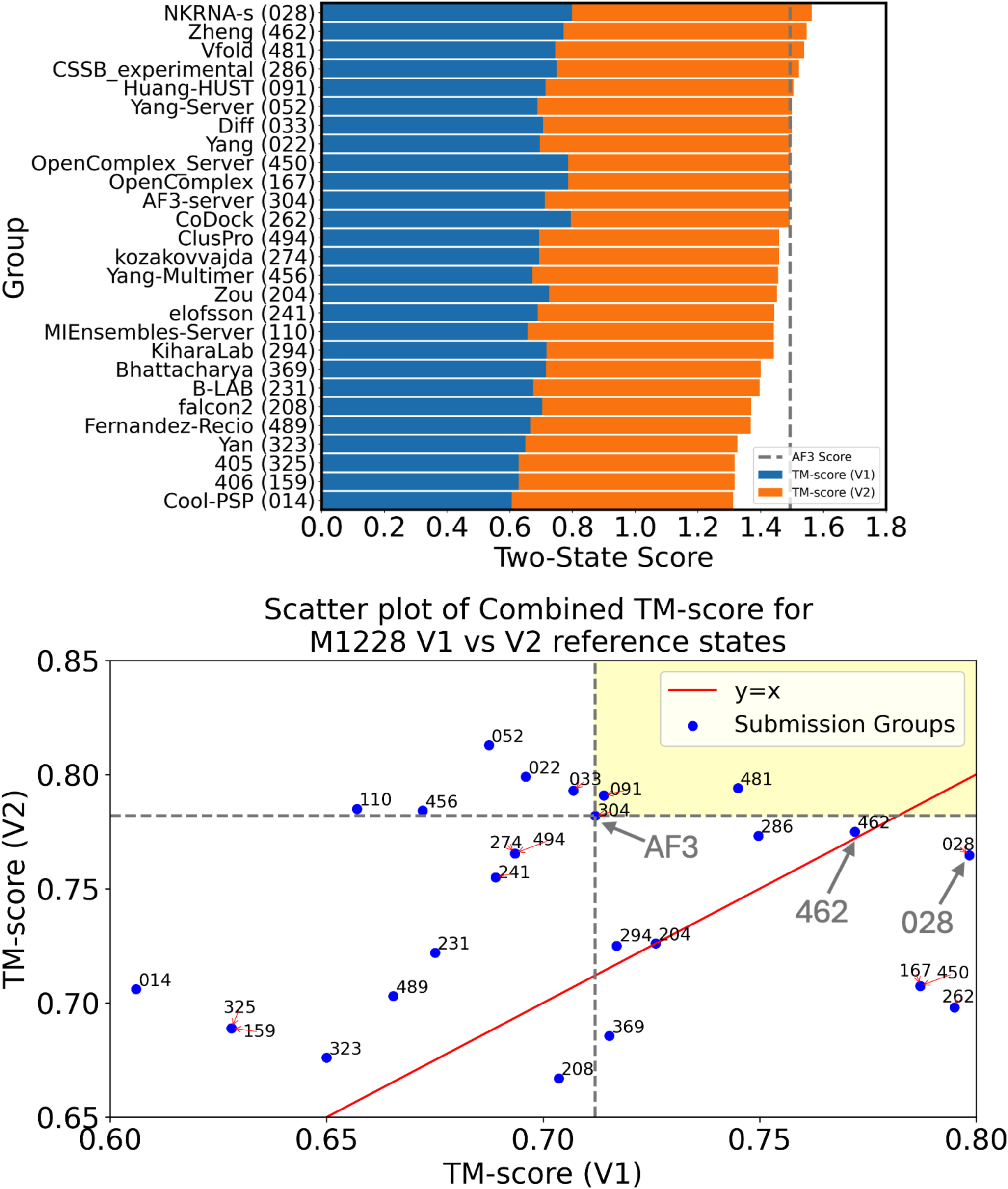
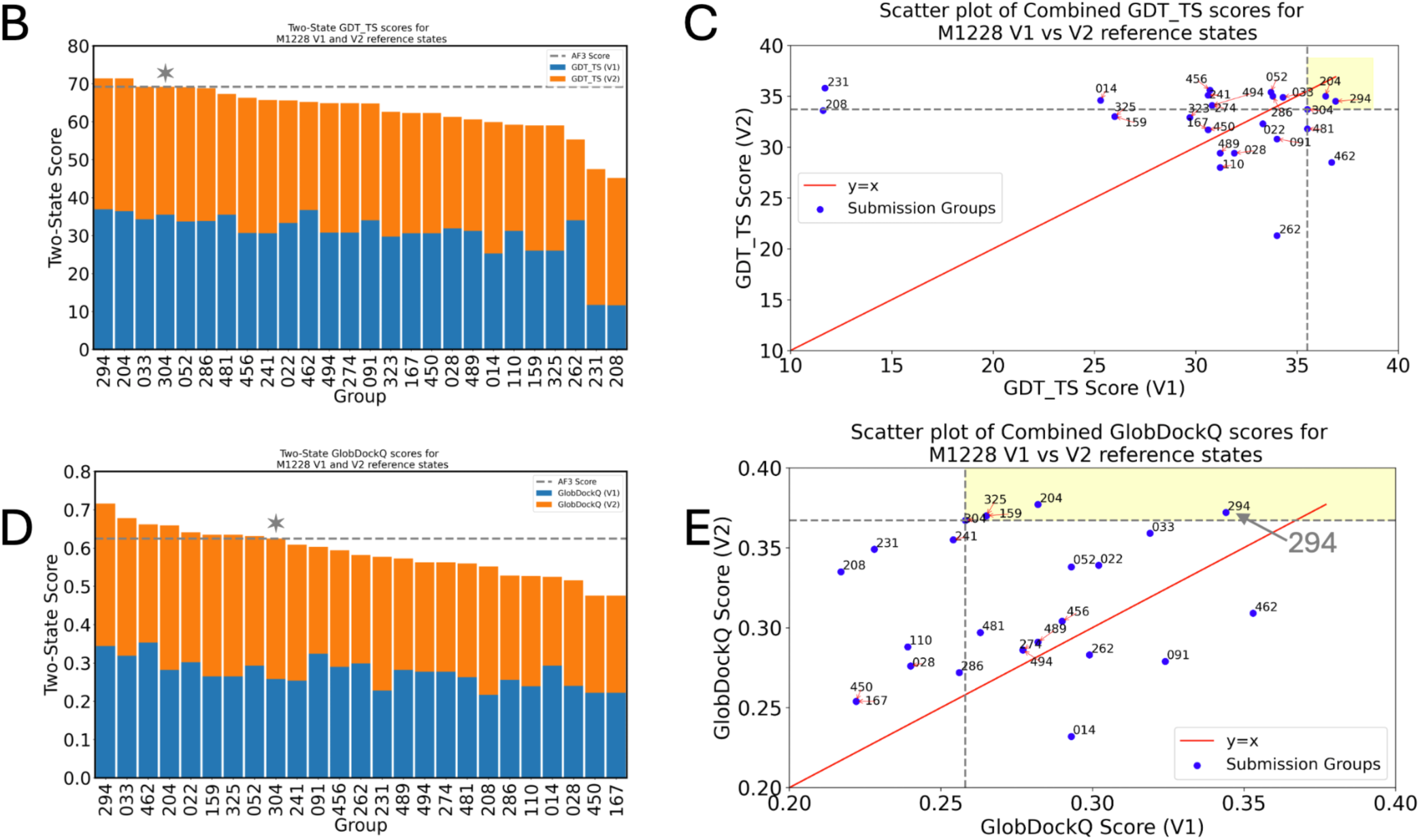
Expanded versions of panels 2B and 2C for target M1228. (B) Composite/aggregate two-state GDT_TS scores plot for all best models ranked by predictor groups to evaluate agreement with both V1 and V2 states. (C) Scatterplot showing the best GDT_TS scores across predictor groups each group’s best models plotted against reference states V1 (x-axis) and V2 (y-axis). (D-E) Same plots as B and C for Global DockQ. Values for best AF3-server (304) models are indicated by * and horizontal dashed lines in panels B and D.

**Fig. S4.**
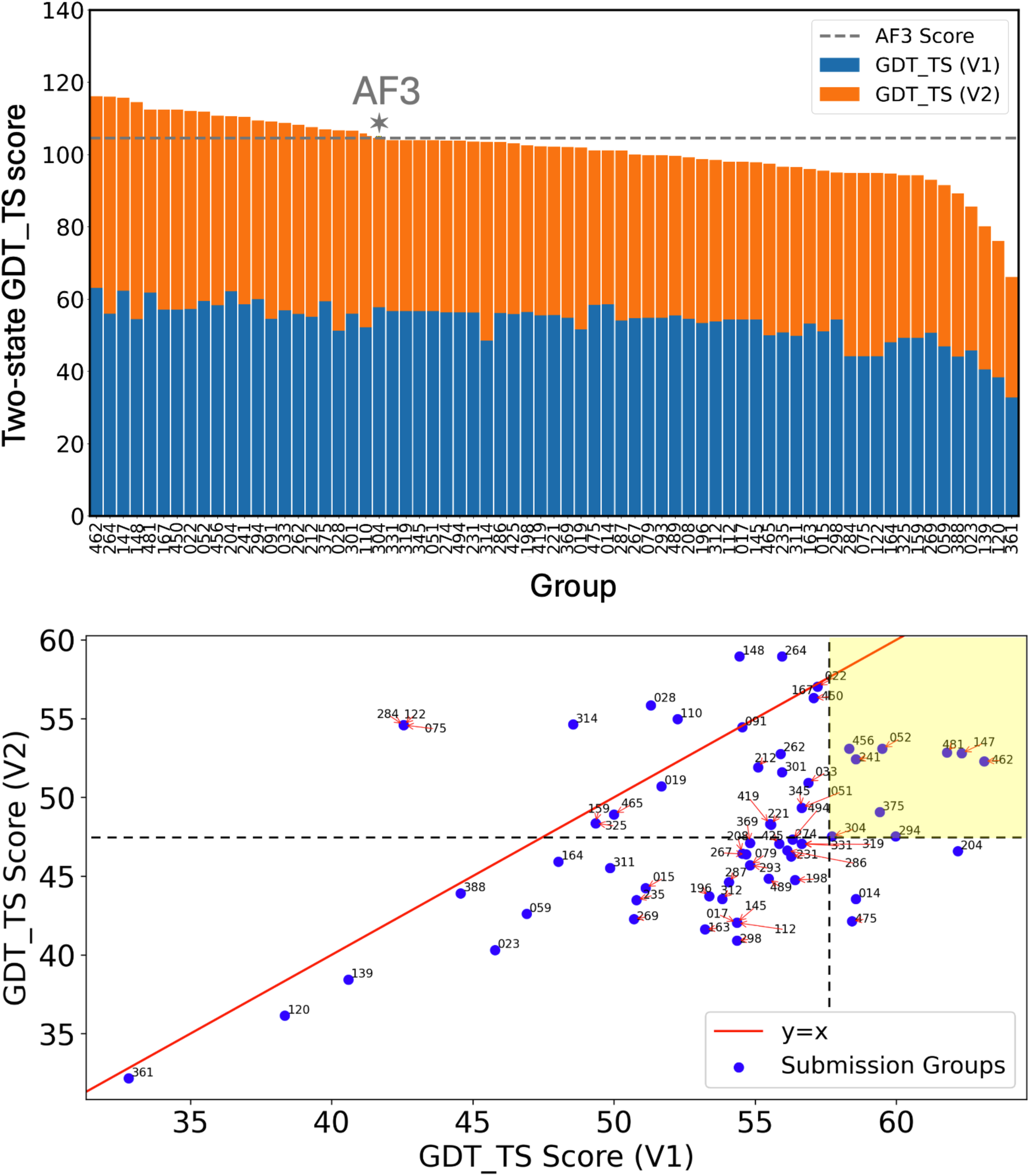
Expanded versions of panels 4C and GDT_TS scatterplot for target T1228. Values for best AF3-server (304) models are indicated in upper panel by * and horizontal dashed line.

**Fig. S5.**
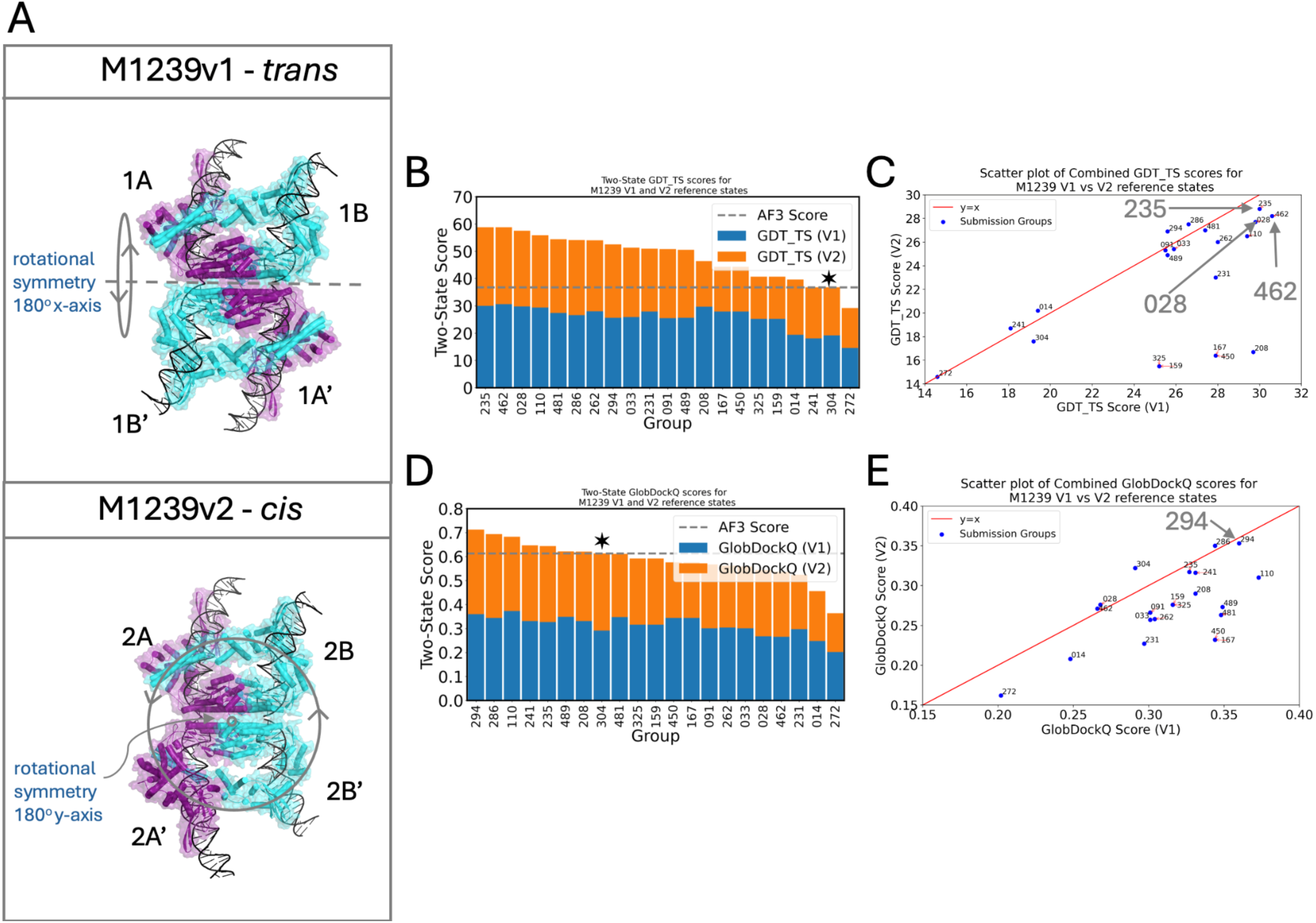
Symmetry in target M1239 and additional two-state assessments. (A) Reference structures of M1239 in two conformational states (V1 and V2), each forming tetrameric assemblies with four DNA fragments. Chains are colored by domain (magenta and cyan), and subunits are labeled to highlight interdomain rearrangements between states. Rotational symmetry axes are marked, which are different for V1 vs V2. (B) Combined two-state GDT_TS scores for best models ranked by predictor groups. (C) Scatterplot of the best GDT_TS scores across predictor groups for both conformational states. (D-E) Same plots as B and C for Global DockQ. Values for best AF3-server (304) models are indicated in panels B and D by * and horizontal dashed line.

**Fig. S6.**
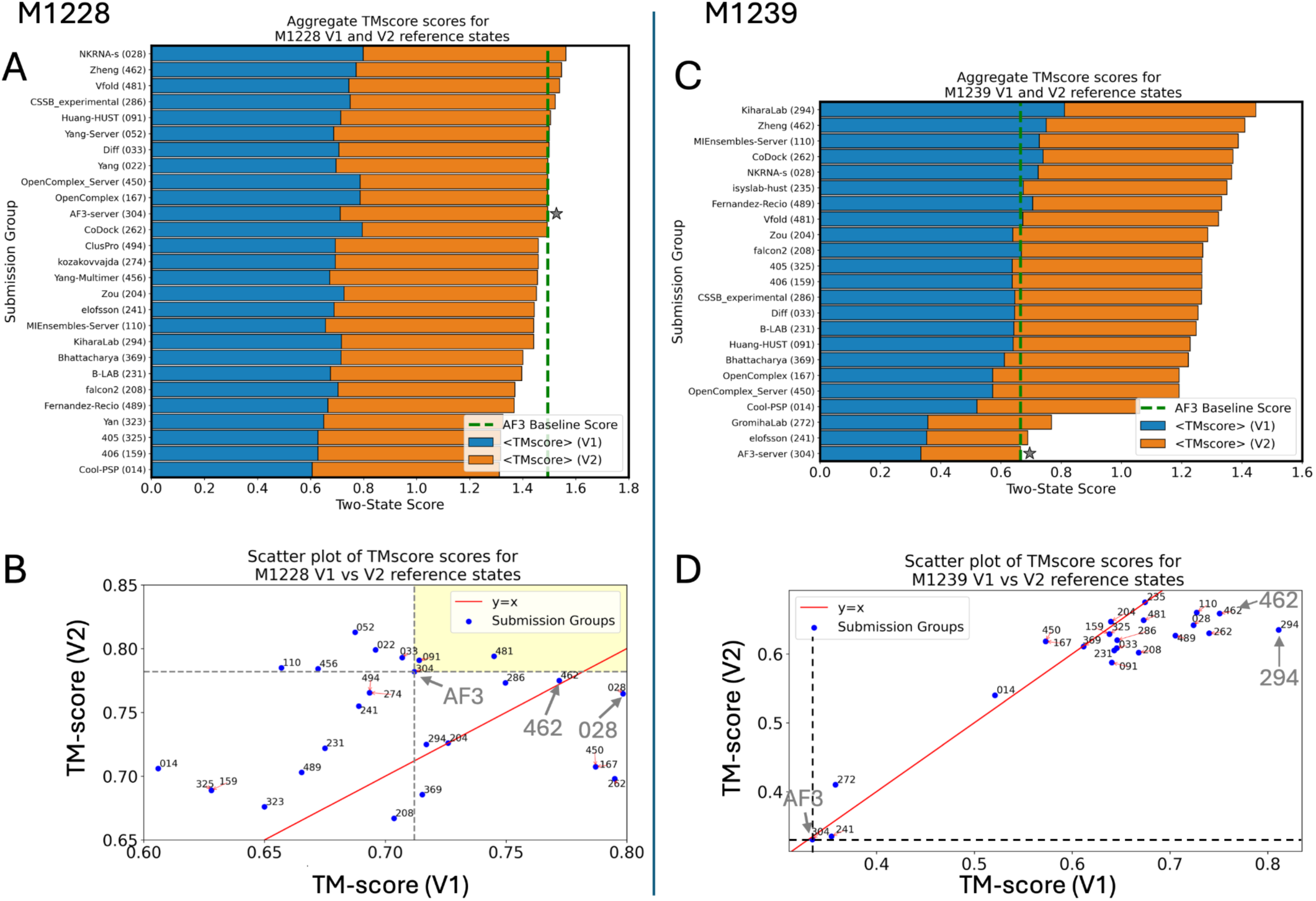
Two state TM-scores plots and scatterplots for M1228 and M1239. Values for best AF3-server (304) models are indicated in panels A and C by a star and vertical dashed line.

**Fig. S7.**
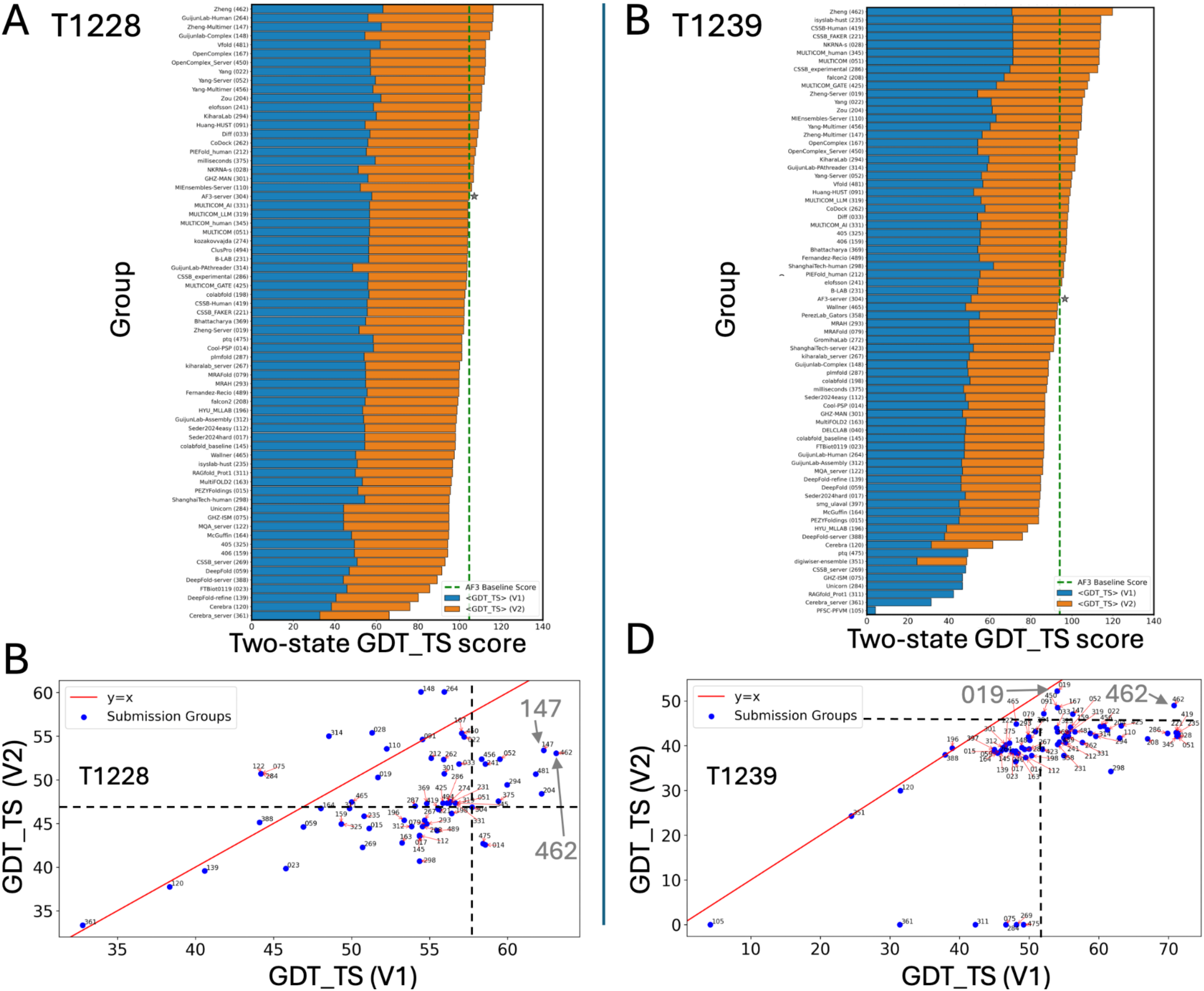
Two state GDT_TS scores plots and scatterplots for T1228 and T1239. Values for best AF3-server (304) models are indicated in panels A and B by * and a vertical dashed line.

**Fig. S8.**
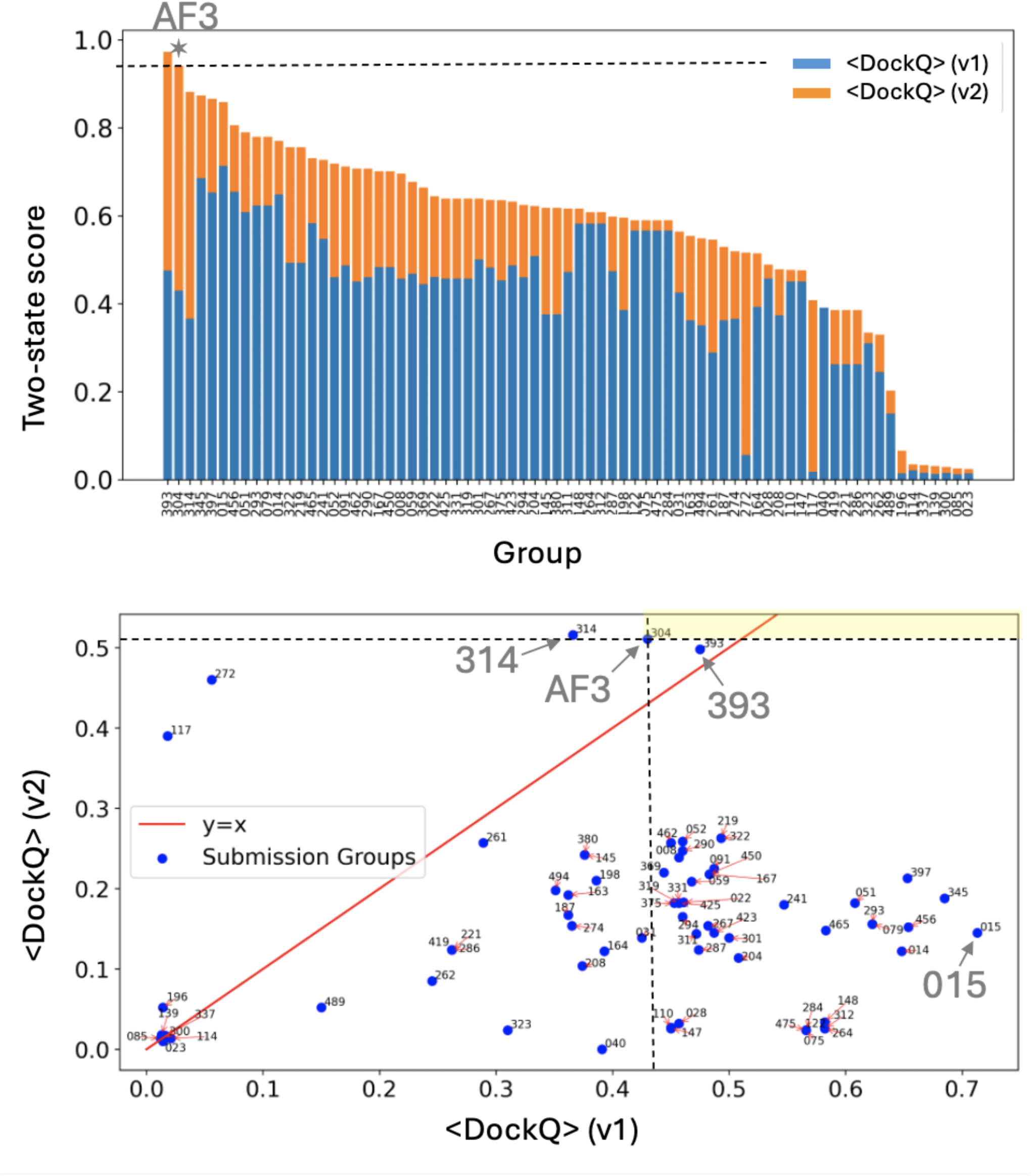
Expanded versions of panels 6B and 6C for target T1249. Values for best AF3-server (304) models are indicated in top panel by * and horizontal dashed line.

**Fig. S9.**
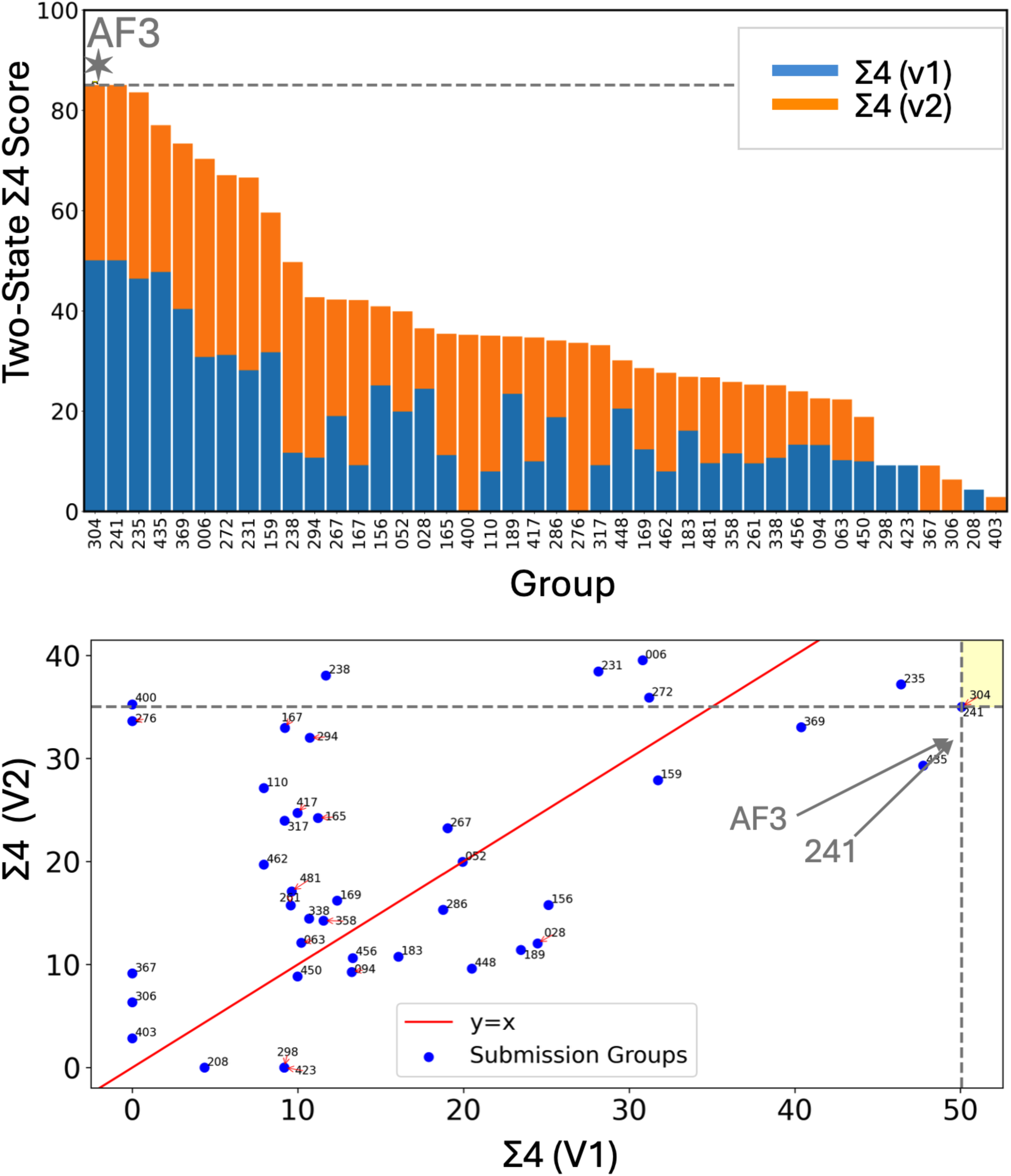
R1203 - Expanded versions of panels 7C and 7D. Values for best AF3-server (304) models are indicated in top panel by * and horizontal dashed line.

**Fig. S10.**
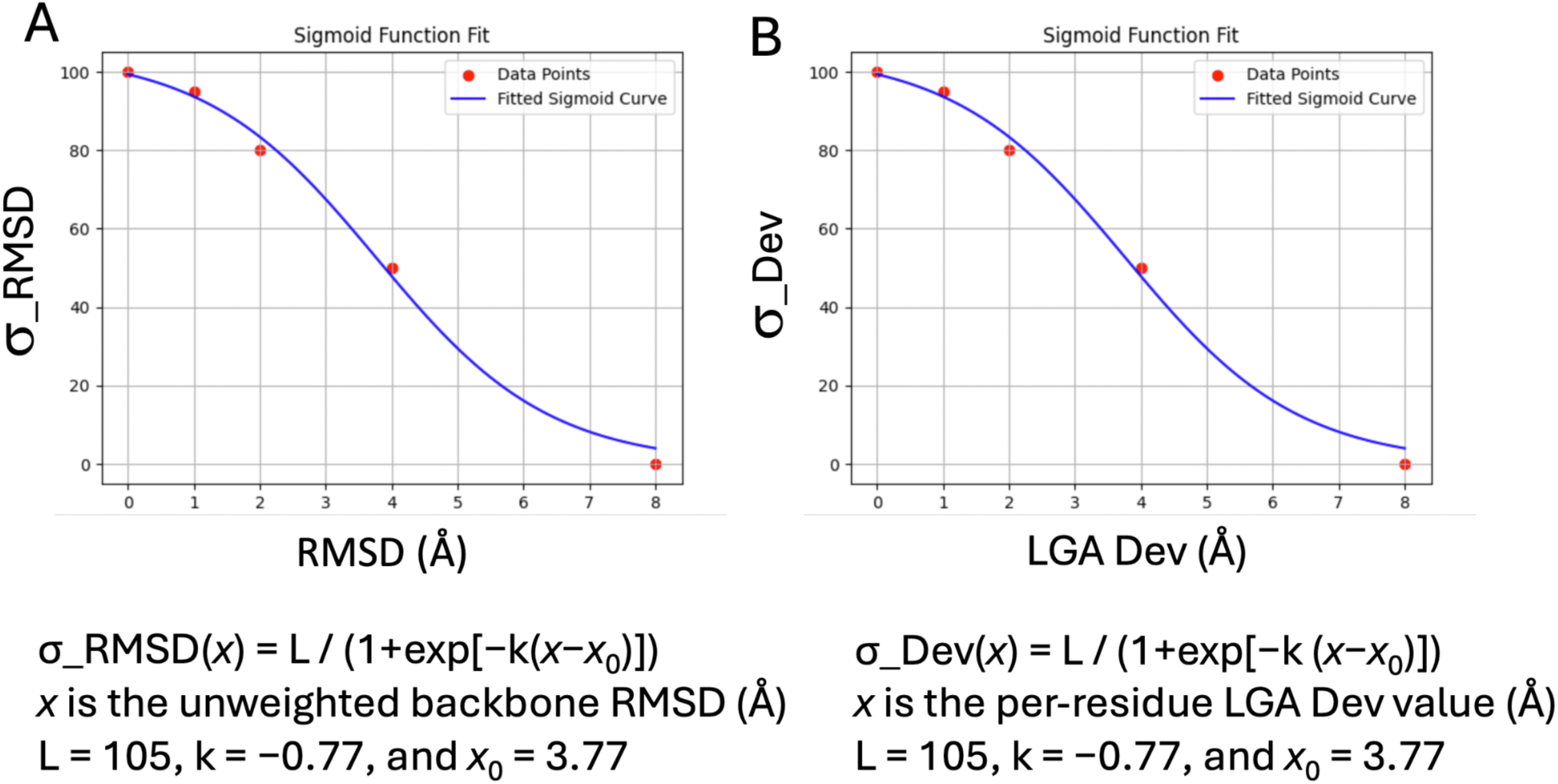
Sigmoid-weighted scoring functions. (A) Fitting of RMSD values to a sigmoid function. (B) Fitting local LGA Dev values to sigmoid function.

**Figure S11.**
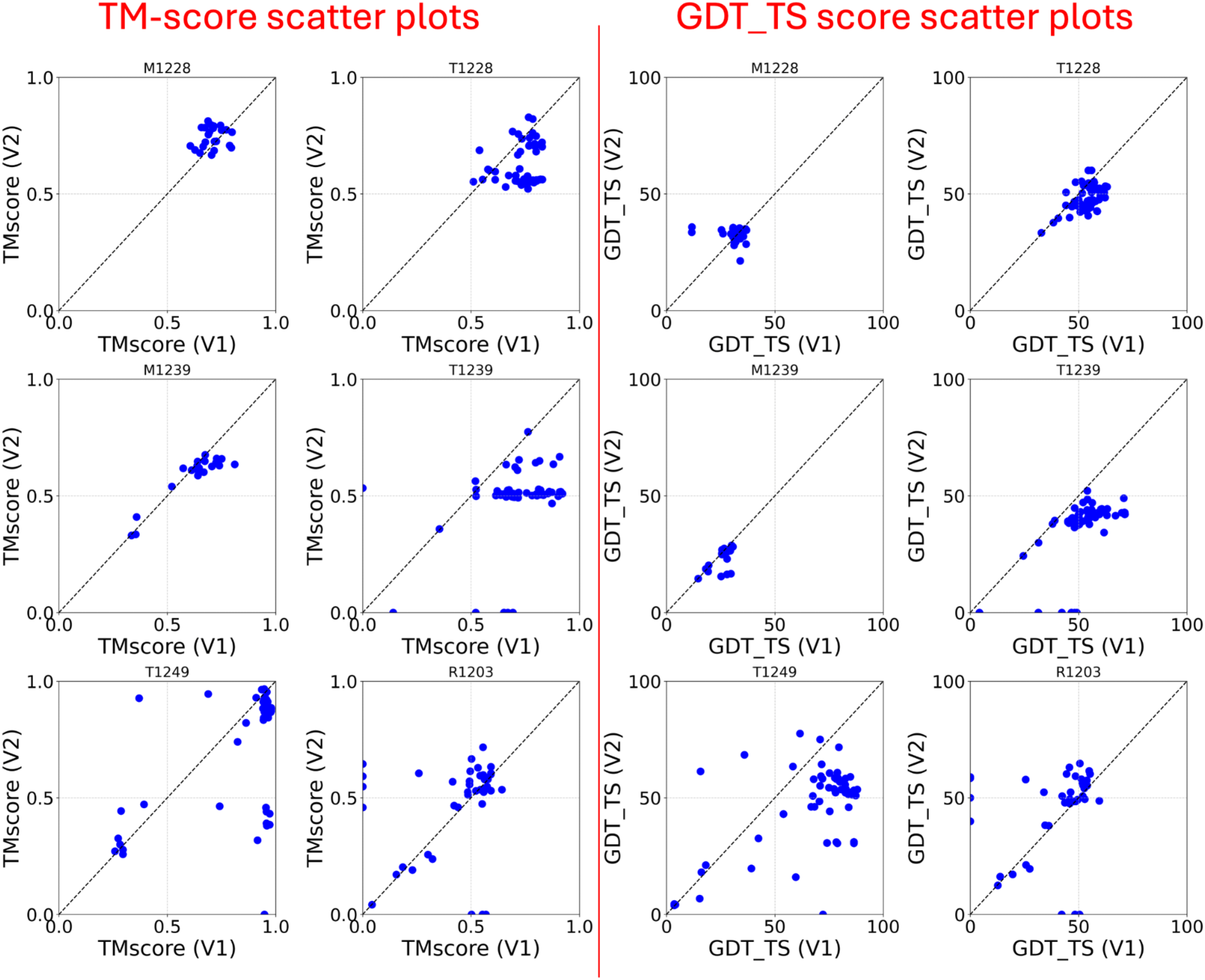
TM and GDT-TS V1 vs V2 scatter plots for the six of the seven most-successfully modeled targets. These plots illustrate the range of RM and GDT_TS scores obtained for the most successful target. While some predictors submitted models with relatively high TM scores (left panels), the corresponding GDT-TS scores (right panels) are generally lower than obtained for typical CASP16 single-state targets. Target 1214 is not included because only one state (holo) was modeled.

## Supplementary Text - Top-performing methods in Ensembles experiment of CASP16

### Summaries of some top-performing modeling methods used for the several most successful targets and target pairs

These summaries were copied from the author provided CASP16 abstracts (https://predictioncenter.org/casp16/doc/CASP16_Abstracts.pdf) and then edited. To emphasize this point the text edited from submitted abstracts is indented.

#### Target T1214holo

The six top-performing groups for T1214holo were PEZYFoldings (015), ShanghaiTech-ligand (386), ShanghaiTech-human (298), falcon2(208), Zheng (462), and SNU-CHEM-lig (408). Despite using different strategies, all captured critical structural rearrangements in the ROIs, with PEZYFoldings and ShanghaiTech-human achieving the highest S4 scores.

**PEZYFoldings (015)** employed an AlphaFold-Multimer v2.3-based pipeline augmented with extensive sequence searches, PLM-based MSA construction (via ESM2), and template selection through PLM-derived alignments. Post-modeling, they used OpenMM for relaxation and a custom rescoring method based on per-residue pLDDT thresholds to refine and rank structures. For large or complex targets, modular modeling and reassembly were applied with visual inspection in PyMOL. Despite not explicitly modeling the ligand for T1214, their method accurately captured ligand-induced conformational changes via robust MSA context and dropout sampling strategies. This text is an edited version of the published CASP16 abstract.
**falcon2 (208)** generated ensembles using AlphaFold3 with at least eight seeds per target and clustered predictions based on TM-score to identify distinct conformers. FoldSeek and ChatGPT (GPT-4) were used to retrieve templates and infer biological assembly strategies. For holo structure prediction, they used AlphaFold3 output and ranked structures via DENOISer energy scoring. Their ligand-free models still captured key ROIs and conformational shifts, reflecting the power of structural diversity and AI-guided inference. This text is an edited version of the published CASP16 abstract.
**ShanghaiTech-human (298) and ShanghaiTech-ligand (386)** both performed among the top five for T1214. However, no specific methodological details were found in the abstract document or supporting materials, limiting further analysis of their strategy. This text is an edited version of the published CASP16 abstract.
**Zheng (462)** is described below.
**SNU-CHEM-lig (408)** focused on ligand-aware modeling using receptor conformational sampling (AlphaFold2 + 3Di), flexible docking via GALigandDock, and deep learning-based scoring with DENOISer. For T1214, their pipeline explicitly addressed the ligand-binding loop flexibility and produced a model with sub-angstrom ligand RMSD. Their multi-tool strategy incorporated conformational benchmarking and re-ranking across several in-house and third-party docking algorithms, emphasizing pocket adaptation. This text is an edited version of the published CASP16 abstract.

##### Summary

These diverse approaches reflect the multiple pathways to success in modeling ligand-induced conformational changes. PEZYFoldings’ MSA-driven ensemble modeling, falcon2’s AI-integrated sampling strategy, and SNU-CHEM-lig’s ligand-centric docking protocol each independently converged on accurate predictions. Their complementary strengths suggest that integrating sequence-based sampling, structural clustering, and ligand-aware docking could further enhance multistate prediction capabilities.

#### M/T Targets: M1228v1/v2, M1239v1/v2, T1228v1/v2, T1239v1/v2

Based on TM two-state scores, the top-performing groups for M1228 and M1239 were: NKRNA-s (028), Zheng (462), Vfold (481), CSSB-experimental (286), Huang-HUST (091), and KiharaLab (294), Zheng (462), MIEnsembles-Server (110), CoDock (262), NKRNA-s (028), respectively. Based on GDT-TS two-state scores, the top-performing groups for T1228 and T1239 were: Zheng (462), GuijunLab-Human (264), Zheng-Multimer (147), Guijunlab-Complex (148), Vfold (481), and Zheng (462), isyslab-hust (235), CSSB-human (419), CSSB_FAKER (221), NKRNA-s (028), MULTICOM_human (Group 345), MULTICOM (051), respectively. The one group that consistently scored at or near the top (though marginally better than other groups) was Zheng (462). All of these best model two-state scores were (sometimes marginally) higher the corresponding scores for AF3-server (304).

**NKRNA-s (Group 028):** The NKRNA-s group developed DeepProtNA for modeling protein-DNA and protein-RNA complexes. For protein-protein and protein-nucleic acid complexes, they used a modified DMFold-Multimer pipeline. MSAs were built using DeepMSA with large metagenomic databases. For heteromers and hybrids, domain-level MSAs were merged into chain-level alignments. Templates were obtained using LOMETS4, which could process complex assemblies. AlphaFold2 variants with extended decoy generation and different dropout settings were used for modeling. Their pipeline addressed stoichiometry estimation and complex model ranking, making it well-suited for hybrid targets involving protein and nucleic acid components. This text is an edited version of the published CASP16 abstract.
**Zheng (462)** employed a robust, modular approach for both protein monomers and multimeric complexes. For monomeric targets, they used D-I-TASSER, a Monte Carlo-based folding simulation pipeline guided by restraints predicted through deep learning—including residue-residue distances, torsion angles, and hydrogen bond networks. Their MSA construction pipeline was based on DeepMSA2 with enhancements for domain-level alignment merging. This text is an edited version of the published CASP16 abstract.
**VFold (481)**, although traditionally focused on RNA structure prediction, extended its capabilities to hybrid and protein-only targets in CASP16. For these tasks, they used AlphaFold3 to predict structural candidates and applied AMBER-based energy minimization for refinement. They incorporated ITScore, a knowledge-based scoring function, to evaluate the quality of models, particularly for protein-nucleic acid and protein-protein interface accuracy. This method proved especially useful for assessing large quaternary complexes where native-like packing and docking geometries were critical. Their pipeline showed flexibility in adapting to hybrid assemblies by integrating structural prediction, docking evaluation, and empirical energetic refinement. This text is an edited version of the published CASP16 abstract.
**CSSB-experimental (286)** used a newly developed approach, UltimateMSA, to generate MSAs by integrating data from custom databases built from public domain multiple databases (nr, tsa_nr, and env_nr). MSAs were constructed using HHblits and MMseqs. They used an in-house protein structure prediction model incorporating elements from RoseTTAFold2 and AlphaFold-Multimer, in parallel with ColabFold. Models were relaxed using Rosetta FastRelax and re-ranked using a composite score of AF2Rank multiplied by iPTM and AFRank. This text is an edited version of the published CASP16 abstract.
**Huang-HUST (091)** utilized a template-guided ensemble docking strategy for multimeric protein targets. Their receptor models were generated using AlphaFold-Multimer and Modeller, forming diverse structural ensembles to accommodate conformational variability. When high-confidence templates were available (e.g., from homologs like FecA for T1214), ligand and subunit poses were inferred using LSAlign and refined through docking tools such as XDock, with final model selection guided by ITScore. This pipeline was particularly effective in capturing alternate binding pocket conformations and modeling ligand-induced domain motions within multimeric proteins. Their strategy demonstrated the value of combining structural homology with ensemble-based conformational sampling and docking refinement. This text is an edited version of the published CASP16 abstract.
**KiharaLab (Group 294).** KiharaLab used template-based modeling followed by MD ensemble generation for protein targets. Specifically for T1200 and T1300, MODELLER generated initial models using known PDB templates, which were further refined with 500 ns MD simulations using the Desmond engine and OPLS4 force field. The resulting 1000 trajectory frames per target were filtered and superimposed to preserve the reference structure topology. This ensemble was submitted as final predictions. The method highlights structure stabilization and refinement via MD simulation for both monomers and multimeric assemblies. This text is an edited version of the published CASP16 abstract.
**MIEnsembles-Server (110)** is designed for modeling a wide range of targets, including protein monomers, nucleic acids (DNA or RNA), nucleic acid-nucleic acid complexes, protein-protein complexes, protein-nucleic acid complexes, ensemble targets, and model quality assessment (QA) targets. Nucleic acid-related targets are predicted using the newly developed deep learning method, DeepProtNA, which integrates pre-trained language model embeddings, multiple sequence alignments (MSA), predicted secondary structures, and structural templates as inputs for modified Evoformer blocks. Protein monomer and multimer targets were modeled using a modified version of DMFold. Ensemble targets are predicted using a clustering approach based on model structural similarity, and QA targets are assessed by selecting the best DMFold model as a reference. This text is an edited version of the published CASP16 abstract.
**CoDock (Group 262):** CoDock focused on ligand-binding prediction, but their method includes hybrid protein-ligand and protein-DNA modeling. They began by searching for homologous PDB structures, retrieving native ligands and clustering them with the provided ones. For docking, they used CoDock-Ligand and AutoDock Vina, accounting for conformational flexibility. For unknown or apo targets (e.g., L5001), blind docking was used. Ligand pose selection incorporated clustering and CNN-based scoring. For DNA targets, AlphaFold3 was used to predict the receptor (protein or hybrid structure), followed by docking. This text is an edited version of the published CASP16 abstract.
**GuijunLab-Human (264)** used Guijunlab-complex, AlphaFold2, AlphaFold3, and HDOCK [24] to predict monomer and complex structure. These models were then used to determine inter-chain distances which were input into DeepAssembly [23], together with the monomer structures, to predict the complex structure. Models were selected using our in-house model quality assessment method. This text is an edited version of the published CASP16 abstract.
**Zheng-Multimer (147)**. For complexes, Zheng applied a modified AlphaFold2 architecture that incorporated paired MSAs, complex-specific template detection through LOMETS4, and extended sampling. Multimeric predictions involved optimizing both stoichiometry and conformation through a scoring function combining pTM and ipTM metrics. This hybrid pipeline allowed Zheng to generalize across diverse target types and achieve top-tier results on many ensemble targets. This text is an edited version of the published CASP16 abstract.
**Guijunlab-Complex (148)** is a pipeline for protein multimer structure modeling using paired MSA construction based on deep learning. A sequence-based deep learning model was used to predict protein-protein structural similarity (pSS-score) and inter-action probability (pIA-score). These predicted scores were used to efficiently construct diverse paired MSAs for multimer prediction by integrating them with species information. Subsequently, we fed the paired MSAs into AlphaFold-Multimer and a previously developed DeepAssembly to generate a series of multimer structures. In-house multi-model quality assessment methods were used for model scoring and selection. This text is an edited version of the published CASP16 abstract.
**isyslab-hust (235)** experimented with AlphaFold2, AlphaFold3, and EsmFold, tuning parameters like num_recycles and random seeds. Structures with low pLDDT scores were refined using Rosetta, and final model selection was based on both pLDDT and TM-score diversity. Their workflow primarily targeted protein monomers but also included AlphaFold3-based complex modeling for selected targets. This text is an edited version of the published CASP16 abstract.
**CSSB-human (419)** took a human-in-the-loop approach, combining structure prediction (ColabFold, AlphaFold, RoseTTAFold) with literature-informed decisions. They stratified targets by confidence: high-confidence models were retained without modification; medium-confidence ones underwent MSA refinement or template search; and low-confidence ones were handled manually. For multimers, complex templates and docking (e.g., GalaxyTongDock, HADDOCK) were used where literature data guided interfaces. Final models were ranked using AF2Rank and refined with Rosetta FastRelax.This text is an edited version of the published CASP16 abstract.
**CSSB_FAKER (221)** employed a structure-augmented MSA refinement strategy to improve AlphaFold-based predictions for protein monomers and multimers (excluding DNA/RNA complexes). They began with ColabFold models, selected high-confidence structures using AF2Rank, and iteratively enhanced MSAs using Foldseek and ProteinMPNN (targeting low pLDDT residues). This process was repeated five times, with final models ranked by AF2Rank and relaxed using energy minimization. For Phase 2, MassiveFold predictions were used and clustered via TM-score to ensure diversity. This text is an edited version of the published CASP16 abstract.
**MULTICOM_human (Group 345) and MULTICOM (051).** Cheng and colleagues applied the latest MULTICOM4, protein structure prediction system, an upgrade of MULTICOM3 [48,49], built on top of AlphaFold2 [4], AlphaFold-Multimer [50], and AlphaFold3 [51] to generate a large number of tertiary structural models. Sequences were searched against various sequence databases using HHblits [52] and JackHMMER [53] to generate a diverse set of MSAs. In addition to using the structural templates identified by the default AlphaFold2, the MSA generated from UniRef was used to search an inhouse template database curated from Protein Data Bank (PDB) to identify alternative templates. For targets extracted from multimeric assemblies, they used AlphaFold-Multimer-derived chains. The structural models were evaluated using multiple quality assessment (QA) methods including GATE, GCPNet-EMA, EnQA, average pairwise similarity score and DeepRank3 methods (DeepRank3_Cluster, DeepRank3_SingleQA, DeepRank3_SingleQA_lite). Their multi-pronged approach ensured a diverse set of high-confidence protein monomer and complex structures. This text is an edited version of the published CASP16 abstract.

##### Summary

Across these groups, commonalities include extensive and comprehensive MSA construction/refinement using tools like HHblits, MMseqs2, or Foldseek, the widespread use of AlphaFold2/AlphaFold-Multimer and increasingly AlphaFold3, and the adoption of confidence metrics (pLDDT, TM-score, AF2Rank) for model selection. Several groups also used structural template detection (LOMETS, Foldseek) and ensemble or human-guided ranking strategies. ProteinMPNN, Rosetta, and MD simulations were selectively used for refinement. Differences lie in the degree of automation (e.g., MULTICOM vs. CSSB-human), MSA enhancement strategies (e.g., NKRNA-s’s domain MSA assembly vs. CSSB_FAKER’s structure-based augmentation), and incorporation of external literature or experimental insights (notably in CSSB-human and CoDock). Not all groups worked on DNA/protein hybrids, but NKRNA-s provided the most comprehensive treatment of such systems using DeepProtNA. Groups with no available or relevant information for monomer/multimer protein or hybrid targets were not summarized here.

#### Target T1249 v1/v2

The five top-performing groups for T1249 based on two-state average interface <DOCKQ> scores were GuijunLab-QA (393), AF3-server (304), GuijunLab-PAthreader (314), Multicom-human (345), smg.ulaval (397).

**GuijunLab-QA (393)** focused on model quality assessment across their various prediction groups using advanced single- and multi-model evaluation strategies. Their primary methodology (QA1) used a global-local consensus approach—globally, they modeled protein structures at multiple abstraction levels with a multi-layer GNN; locally, they assessed interfaces using protein language model embeddings combined with AlphaFold2 representations through attention-based modules. These assessments fed into downstream structural alignment-based consensus methods. For complex assemblies and trimeric units, their QA4 and QA5 pipelines integrated structural clustering, scoring via GraphCPLMQA, and Voronoi-based local interface evaluations. Models from various predictors (including MassiveFold and AlphaFold3) were pooled and iteratively scored to select high-confidence submissions. This text is an edited version of the published CASP16 abstract.
**GuijunLab-PAthreader (314)** employed a robust multi-step protocol integrating enhanced MSA generation, remote homology-based template detection, and refined model quality assessment. For MSA generation, they combined traditional database searches (Uniclust30, UniRef90, MGnify, BFD) with structure-informed searches from AlphaFold-predicted models using Foldseek. This yielded 11 diverse MSAs per target. For template recognition, the group’s PAthreader method utilized structural alignment against PAcluster80 (a structure-clustered DB) with a three-track alignment approach. Their DeepMDisPre tool predicted distance restraints which, along with structural profiles, were processed by a convolutional self-attention network to yield DMScore, used for ranking templates. AlphaFold2 was run on combinations of MSAs and templates to generate structural models. These were ranked using both AlphaFold2’s intrinsic scoring and their in-house model quality assessment to select the best five predictions. This text is an edited version of the published CASP16 abstract.
**AF3-server (304)** group submitted both manual (Elofsson - 241) and automated predictions using AlphaFold3. For trimeric and other multimeric targets, the team tested multiple stoichiometries ranging from monomers to hexamers, guided by AlphaFold3’s pTM and PAE confidence metrics. If the stoichiometry was unclear, the models were screened for ranking scores and evaluated for assembly consistency. When targets were too large to run on AlphaFold3 directly, a fragmentation and reassembly strategy was used to reconstruct full models. Manual interventions included structure inspection and clustering to select diverse submissions. This text is an edited version of the published CASP16 abstract.
**MULTICOM_Human (345)** extended the MULTICOM4 framework built on AlphaFold2 and AlphaFold3 to generate large ensembles of models for trimeric and multimeric predictions. They emphasized massive sampling using diverse MSA/template combinations, with up to 300,000 models per target in some cases. Their model selection strategy combined automated ranking metrics—plDDT, GATE (graph-based accuracy estimation), EnQA, GCPNet-EMA, and DeepRank—with human intervention. Trimeric monomers extracted from multimer targets were further refined and ranked based on consensus from multiple quality metrics. The group’s extensive sampling and multi-pronged ranking allowed them to submit highly confident models for complex assemblies including trimers. This text is an edited version of the published CASP16 abstract.
**smg_ulaval (397).** Unfortunately, no abstract or methodological details were provided for the smg_ulaval group in the CASP16 documents reviewed. As such, we cannot comment on their specific protocol for trimeric protein prediction.

##### Summary

Among the top performing groups, GuijunLab-PAthreader (314), GuijunLab-QA (393), AF3-server (304), and MULTICOM_Human (345) reported careful handling of trimeric protein units, albeit with different emphases. GuijunLab groups focused heavily on model quality assessment and structure-guided MSA enhancement. AF3-server prioritized efficient stoichiometry prediction with practical interventions for large systems, while MULTICOM_Human (345) excelled in massive model generation and quality-ranking frameworks combining AI and human expertise.

#### RNA target R1203

The five top-performing groups for R1203 based on two-state 𝛴4 global/local composite scores were AF3-server (304), elofsson (241), isyslab-hust (235), RNAFOLDX (435), and Bhattachayra (369).

**AF3-server** is described above.

**elofsson (241)** submitted manual predictions based on models generated with AlphaFold3 server [51]. Quality estimations were performed using pDockQ2, pDockQ23, and Pcons4. They also extend their work to RNA structure prediction, integrating secondary structure from the Ribonanza challenge with AlphaFold3, HelixFold3, and RosettaFoldNA for 3D modeling. This text is an edited version of the published CASP16 abstract.
**isyslab-hust (235)** did not describe RNA-specific protocols in detail, they used a diverse ensemble of structure prediction tools—AlphaFold2, AlphaFold3, and EsmFold—to model biomolecular structures, likely including RNA. Their pipeline relied on extensive sampling: varying AlphaFold2 parameters such as num_recycles (up to 80) and random seeds (100 values), followed by Rosetta-based refinement of low-confidence models. Final submissions were filtered using pLDDT and TM-score metrics to ensure diversity and structural reliability. Given that they ranked among the top groups for the RNA target R1203, their approach suggests effective indirect handling of RNA targets through careful sampling and rigorous quality filtering, although RNA-specific tools or modeling choices were not explicitly outlined. This text is an edited version of the published CASP16 abstract.
**RNAFOLDX** (435) is a hierarchical approach for fully automated RNA tertiary structure prediction, combining structural templates by threading alignments and inter-nucleotide geometries predicted by a deep residual convolutional neural network. The method consists of four main steps: (1) RNA template detection by threading method, (2) multiple sequence alignment generation, (3) inter-nucleotide geometries prediction, and (3) structure selection, optimization, and refinement. This text is an edited version of the published CASP16 abstract.
**Bhattachayra (369)** integrates AlphaFold3 and AlphaFold2, using AF3 models as templates with custom MSAs in AF2 (ColabFold), applying AF2_ptm and AF2_multimer_v3 weights with tuned recycles for monomeric and multimeric targets, while default AF2 handled very large inputs (>5,000 residues). In parallel, they developed a scoring-guided RNA structure pipeline (lociPARSE) to rank ensembles from diverse RNA predictors, and an integrative protein-RNA complex predictor (ProRNA3D-single) combining language-model embeddings with structural methods. Final submissions were confidence- or score-ranked and refined with PyRosetta, with lociPARSE and ProRNA3D-single released as open-source tools. This text is an edited version of the published CASP16 abstract.

##### Summary

The top performing group was AlphaFold3. Most other top-performing groups built on AlphaFold2, but did not perform quite as well at the AlphaFold3-server. **RNAFOLDX** (435) utilized a distinct convolutional neural network for modeling, with performance rivaling AlphaFold3.

## References

1. A. Kryshtafovych, T. Schwede, M. Topf, K. Fidelis, J. Moult. “Critical Assessment of Methods of Protein Structure Prediction (Casp)-Round Xiv,” Proteins 89 (2021): 1607–1617. 10.1002/10.1002/prot.26237.

2. A. Kryshtafovych, T. Schwede, M. Topf, K. Fidelis, J. Moult. “Critical Assessment of Methods of Protein Structure Prediction (Casp)-Round Xv,” Proteins 91 (2023): 1539–1549. 10.1002/10.1002/prot.26617.

3. A. Kryshtafovych, G. T. Montelione, D. J. Rigden, S. Mesdaghi, E. Karaca, J. Moult. “Breaking the Conformational Ensemble Barrier: Ensemble Structure Modeling Challenges in Casp15,” Proteins 91 (2023): 1903–1911. 10.1002/10.1002/prot.26584.

4. J. Jumper, R. Evans, A. Pritzel, et al. “Highly Accurate Protein Structure Prediction with Alphafold,” Nature 596 (2021): 583–589. 10.1002/10.1038/s41586-021-03819-2.

5. A. C. McBride, F. Yu, E. H. Cheng, et al. “Predicting Pose Distribution of Protein Domains Connected by Flexible Linkers Is an Unsolved Problem,” bioRxiv (2025): 2025.2004.2027.650885. 10.1002/10.1101/2025.04.27.650885.

6. A. Elofsson, R. C. Kretsch, M. Magnus, G. T. Montelione. “Engaging the Community: Casp Special Interest Groups,” Proteins (2025): 10.1002/10.1002/prot.26833.

7. G. T. Montelione. Casp-Sig on Conformational Ensembles of Proteins. 2025; https://www.youtube.com/@CASPSIGConformationalEnsembles..

8. T. A. Ramelot, R. Tejero, G. T. Montelione. “Representing Structures of the Multiple Conformational States of Proteins,” Curr Opin Struct Biol 83 (2023): 102703. 10.1002/10.1016/j.sbi.2023.102703.

9. A. C. Gibbs, R. Steele, G. Liu, B. A. Tounge, G. T. Montelione. “Inhibitor Bound Dengue Ns2b-Ns3pro Reveals Multiple Dynamic Binding Modes,” Biochemistry 57 (2018): 1591–1602. 10.1002/10.1021/acs.biochem.7b01127.

10. A. G. Palmer III. “Enzyme Dynamics from Nmr Spectroscopy,” Accounts of Chemical Research 48 (2015): 457–465..

11. P. Vallurupalli, G. Bouvignies, L. E. Kay. “Studying “Invisible” Excited Protein States in Slow Exchange with a Major State Conformation,” Journal of the American Chemical Society 134 (2012): 8148–8161. 10.1002/10.1021/ja3001419.

12. I. Bertini, C. Del Bianco, I. Gelis, et al. “Experimentally Exploring the Conformational Space Sampled by Domain Reorientation in Calmodulin,” Proc Natl Acad Sci U S A 101 (2004): 6841–6846. 10.1002/10.1073/pnas.0308641101.

13. Y. Qi, J. W. Martin, A. W. Barb, et al. “Continuous Interdomain Orientation Distributions Reveal Components of Binding Thermodynamics,” Journal of Molecular Biology 430 (2018): 3412–3426. 10.1002/10.1016/j.jmb.2018.06.022.

14. C. A. Brosey, J. A. Tainer. “Evolving Saxs Versatility: Solution X-Ray Scattering for Macromolecular Architecture, Functional Landscapes, and Integrative Structural Biology,” Current Opinion in Structural Biology 58 (2019): 197–213. 10.1002/10.1016/j.sbi.2019.04.004.

15. M. Hammel, J. A. Tainer. “X-Ray Scattering Reveals Disordered Linkers and Dynamic Interfaces in Complexes and Mechanisms for DNA Double-Strand Break Repair Impacting Cell and Cancer Biology,” Protein Science 30 (2021): 1735–1756. 10.1002/10.1002/pro.4133.

16. F. Munder, M. Voutsinos, K. Hantke, H. Venugopal, R. Grinter. “High-Affinity Pqq Import Is Widespread in Gram-Negative Bacteria,” bioRxiv (2024): 2024.2006.2004.597491. 10.1002/10.1101/2024.06.04.597491.

17. M. C. M. Smith. “Phage-Encoded Serine Integrases and Other Large Serine Recombinases,” Microbiology Spectrum 3 (2015): 10.1128/microbiolspec.mdna1123-0059-2014. https://doi.org/10.1002/doi:10.1128/microbiolspec.mdna3-0059-2014.

18. H. Shin, Y. Pigli, T. Peña Reyes, J. R. Fuller, F. J. Olorunniji, P. A. Rice. “Structural Basis of Directionality Control in Large Serine Integrases,” bioRxiv (2025): 2025.2001.2003.631226. 10.1002/10.1101/2025.01.03.631226.

19. W. Li, S. Kamtekar, Y. Xiong, G. J. Sarkis, N. D. F. Grindley, T. A. Steitz. “Structure of a Synaptic Γδ Resolvase Tetramer Covalently Linked to Two Cleaved Dnas,” Science 309 (2005): 1210–1215. 10.1002/doi:10.1126/science.1112064.

20. C. S. Trejo, R. S. Rock, W. M. Stark, M. R. Boocock, P. A. Rice. “Snapshots of a Molecular Swivel in Action,” Nucleic Acids Research 46 (2018): 5286–5296. 10.1002/10.1093/nar/gkx1309.

21. M. van Kempen, S. S. Kim, C. Tumescheit, et al. “Fast and Accurate Protein Structure Search with Foldseek,” Nature Biotechnology 42 (2024): 243–246. 10.1002/10.1038/s41587-023-01773-0.

22. S. Kamtekar, R. S. Ho, M. J. Cocco, et al. “Implications of Structures of Synaptic Tetramers of Γδ Resolvase for the Mechanism of Recombination,” Proceedings of the National Academy of Sciences 103 (2006): 10642–10647. 10.1002/doi:10.1073/pnas.0604062103.

23. Y. Xia, K. Zhao, D. Liu, X. Zhou, G. Zhang. “Multi-Domain and Complex Protein Structure Prediction Using Inter-Domain Interactions from Deep Learning,” Communications Biology 6 (2023): 1221. 10.1002/10.1038/s42003-023-05610-7.

24. Y. Yan, D. Zhang, P. Zhou, B. Li, S.-Y. Huang. “Hdock: A Web Server for Protein–Protein and Protein–DNA/Rna Docking Based on a Hybrid Strategy,” Nucleic Acids Research 45 (2017): W365–W373. 10.1002/10.1093/nar/gkx407.

25. R. C. Kretsch, R. Albrecht, E. S. Andersen, et al. “Functional Relevance of Casp16 Nucleic Acid Predictions as Evaluated by Structure Providers,” bioRxiv (2025): 2025.2004.2015.649049. 10.1002/10.1101/2025.04.15.649049.

26. H. Cohen-Dvashi, M. Katz, R. Diskin. “Metal-Induced Conformational Changes in the Sabiá Virus Spike Complex,” Nature Microbiology (2025): 10.1002/10.1038/s41564-025-02075-8.

27. S. Basu, B. Wallner. “Dockq: A Quality Measure for Protein-Protein Docking Models,” PLoS One 11 (2016): e0161879. 10.1002/10.1371/journal.pone.0161879.

28. J. Fernandes, B. Jayaraman, A. Frankel. “The Hiv-1 Rev Response Element,” RNA Biology 9 (2012): 6–11. 10.1002/10.4161/rna.9.1.18178.

29. M. Ojha, L. Hudson, A. Photenhauer, et al. “Structural Plasticity of Rre Stem-Loop Ii Modulates Nuclear Export of Hiv-1 Rna,” bioRxiv (2025): 2025.2002.2020.639096. 10.1002/10.1101/2025.02.20.639096.

30. R. C. Kretsch, A. M. Hummer, S. He, et al. “Assessment of Nucleic Acid Structure Prediction in Casp16,” bioRxiv (2025): 2025.2005.2006.652459. 10.1002/10.1101/2025.05.06.652459.

31. D. Del Alamo, D. Sala, H. S. McHaourab, J. Meiler. “Sampling Alternative Conformational States of Transporters and Receptors with Alphafold2,” Elife 11 (2022): 10.1002/10.7554/eLife.75751.

32. T. Saldano, N. Escobedo, J. Marchetti, et al. “Impact of Protein Conformational Diversity on Alphafold Predictions,” Bioinformatics 38 (2022): 2742–2748. 10.1002/10.1093/bioinformatics/btac202.

33. H. K. Wayment-Steele, A. Ojoawo, R. Otten, et al. “Predicting Multiple Conformations Via Sequence Clustering and Alphafold2,” Nature 625 (2024): 832–839. 10.1002/10.1038/s41586-023-06832-9.

34. D. Chakravarty, J. W. Schafer, E. A. Chen, et al. “Alphafold Predictions of Fold-Switched Conformations Are Driven by Structure Memorization,” Nat Commun 15 (2024): 7296. 10.1002/10.1038/s41467-024-51801-z.

35. T. Xie, J. Huang. “Can Protein Structure Prediction Methods Capture Alternative Conformations of Membrane Transporters?,” J Chem Inf Model 64 (2024): 3524–3536. 10.1002/10.1021/acs.jcim.3c01936.

36. P. Bryant, F. Noe. “Structure Prediction of Alternative Protein Conformations,” Nat Commun 15 (2024): 7328. 10.1002/10.1038/s41467-024-51507-2.

37. Y. Kalakoti, B. Wallner. “Afsample2 Predicts Multiple Conformations and Ensembles with Alphafold2,” Commun Biol 8 (2025): 373. 10.1002/10.1038/s42003-025-07791-9.

38. D. Chakravarty, M. Lee, L. L. Porter. “Proteins with Alternative Folds Reveal Blind Spots in Alphafold-Based Protein Structure Prediction,” Current Opinion in Structural Biology 90 (2025): 102973. 10.1002/10.1016/j.sbi.2024.102973.

39. M. Lazou, O. Khan, T. Nguyen, et al. “Predicting Multiple Conformations of Ligand Binding Sites in Proteins Suggests That Alphafold2 May Remember Too Much,” Proc Natl Acad Sci U S A 121 (2024): e2412719121. 10.1002/10.1073/pnas.2412719121.

40. G. V. T. Swapna, N. Dube, M. J. Roth, G. T. Montelione. “Memorization Bias Impacts Modeling of Alternative Conformational States of Solute Carrier Membrane Proteins with Methods from Deep Learning.,” bioRxiv (2024): 10.1002/10.1101/2024.07.15.603529.

41. A. Fadini, M. Li, A. J. McCoy, et al. “Alphafold as a Prior: Experimental Structure Determination Conditioned on a Pretrained Neural Network,” bioRxiv (2025): 10.1002/10.1101/2025.02.18.638828.

42. A. Kryshtafovych, B. Monastyrskyy, K. Fidelis. “Casp Prediction Center Infrastructure and Evaluation Measures in Casp10 and Casp Roll,” Proteins: Structure, Function, and Bioinformatics 82 (2014): 7–13. 10.1002/10.1002/prot.24399.

43. A. Kryshtafovych, M. Milostan, L. M.F., e. al. “Updates to the Casp Infrastructure in 2024.,” Proteins (2025): 10.1002/ 10.1002/prot.70042.

44. C. Zhang, M. Shine, A. M. Pyle, Y. Zhang. “Us-Align: Universal Structure Alignments of Proteins, Nucleic Acids, and Macromolecular Complexes,” Nature Methods 19 (2022): 1109–1115. 10.1002/10.1038/s41592-022-01585-1.

45. W. L. DeLano. The Pymol Molecular Graphics System, Version 1.2r3pre, Schrödinger, Llc. In:2025..

46. A. Zemla. “Lga: A Method for Finding 3d Similarities in Protein Structures,” Nucleic Acids Research 31 (2003): 3370–3374. 10.1002/10.1093/nar/gkg571.

47. V. Mariani, M. Biasini, A. Barbato, T. Schwede. “Lddt: A Local Superposition-Free Score for Comparing Protein Structures and Models Using Distance Difference Tests,” Bioinformatics 29 (2013): 2722–2728. 10.1002/10.1093/bioinformatics/btt473.

48. J. Liu, Z. Guo, T. Wu, R. S. Roy, C. Chen, J. Cheng. “Improving Alphafold2-Based Protein Tertiary Structure Prediction with Multicom in Casp15,” Communications Chemistry 6 (2023): 188. 10.1002/10.1038/s42004-023-00991-6.

49. J. Liu, Z. Guo, T. Wu, et al. “Enhancing Alphafold-Multimer-Based Protein Complex Structure Prediction with Multicom in Casp15,” Communications Biology 6 (2023): 1140. 10.1002/10.1038/s42003-023-05525-3.

50. R. Evans, M. O’Neill, A. Pritzel, et al. “Protein Complex Prediction with Alphafold-Multimer,” bioRxiv (2021): 2021.2010.2004.463034. 10.1002/10.1101/2021.10.04.463034.

51. J. Abramson, J. Adler, J. Dunger, et al. “Accurate Structure Prediction of Biomolecular Interactions with Alphafold 3,” Nature 630 (2024): 493–500. 10.1002/10.1038/s41586-024-07487-w.

52. M. Remmert, A. Biegert, A. Hauser, J. Söding. “Hhblits: Lightning-Fast Iterative Protein Sequence Searching by Hmm-Hmm Alignment,” Nature Methods 9 (2012): 173–175. 10.1002/10.1038/nmeth.1818.

53. L. S. Johnson, S. R. Eddy, E. Portugaly. “Hidden Markov Model Speed Heuristic and Iterative Hmm Search Procedure,” BMC Bioinformatics 11 (2010): 431. 10.1002/10.1186/1471-2105-11-431.

